# Reinstating targeted protein degradation with DCAF1 PROTACs in CRBN PROTAC resistant settings

**DOI:** 10.1101/2023.04.09.536153

**Authors:** Martin Schröder, Martin Renatus, Xiaoyou Liang, Fabian Meili, Thomas Zoller, Sandrine Ferrand, Francois Gauter, Xiaoyan Li, Fred Sigoillot, Scott Gleim, Marie-Therese Stachyra, Jason Thomas, Damien Begue, Peggy Lefeuvre, Rita Andraos-Rey, BoYee Chung, Renate Ma, Seth Carbonneau, Benika Pinch, Andreas Hofmann, Markus Schirle, Niko Schmiedberg, Patricia Imbach, Delphine Gorses, Keith Calkins, Bea Bauer-Probst, Magdalena Maschlej, Matt Niederst, Rob Maher, Martin Henault, John Alford, Erik Ahrne, Greg Hollingworth, Nicolas H. Thomä, Anna Vulpetti, Thomas Radimerski, Philipp Holzer, Claudio R. Thoma

## Abstract

Targeted protein degradation (TPD) of neo-substrates with proteolysis targeting chimeras (PROTACs) or molecular glues has emerged as a key modality in exploring new biology as well as designing new drug candidates where catalytic inhibition is neither efficacious nor an option. TPD is mediated through harnessing E3 ligases and redirecting them to ubiquitinate *de novo* target proteins for subsequent proteasomal degradation. Until recently, E3 ligase chemical matter available for mediating TPD has been limited to a relatively low number of ligases, considering that over 600 E3 ligases are encoded by the human genome. In addition, the most utilized ligase for TPD approaches, CRBN, has been observed to be downregulated in settings of acquired resistance to immunomodulatory inhibitory drugs (IMiDs). IMiDs are molecular glues that target IKZF transcription factors to CRBN for degradation. Resistance is potentially accelerated by non-essentiality of CRBN for cell viability. Here we investigated if the essential E3 ligase receptor DCAF1 can be harnessed for TPD utilizing a potent, non-covalent DCAF1 binder. We show that this binder, selective for the CRL4^DCAF1^ E3 ligase complex, can be functionalized into an efficient DCAF1-BRD9 PROTAC. Chemical and genetic rescue experiments confirm specific degradation via the CRL4^DCAF1^ E3 ligase. We further highlight the versatility of DCAF1 for TPD by developing a DCAF1-dasatininb PROTAC targeting multiple cytosolic and membrane bound tyrosine kinases. We expand these findings towards Bruton’s tyrosine kinase (BTK) selective PROTACs and through extensive optimization and characterization efforts share key observations that led to a potent and selective DCAF1-BTK PROTAC (DBt-10). Finally, with this PROTAC DBt-10, we show rescue of BTK degradation in a BTK-dependent, CRBN-degradation-resistant cell line and provide a rationale for E3 ligase swap to overcome CRBN mediated resistance.

## INTRODUCTION

Targeted protein degradation (TPD) to eliminate pathological proteins through the cellular disposal system, the ubiquitin-proteasome pathway, has been actively pursued for more than a decade as a therapeutic principle with multiple protein degraders in clinical development (Békés et al., 2022). Rather than inhibiting proteins, TPD mediates elimination thereof through modification with ubiquitin chains tagging them for degradation. On a molecular level, this is enabled by two classes of molecules; both induce proximity between an E3 ligase and the pathological target protein (Ciechanover, 2005). The first class are bivalent molecules also known as proteolysis targeting chimeras (PROTACs) that induce proximity through two independent binding events mediated by two separate binding moieties connected by a linker ((Sakamoto et al., 2001) and reviewed by Hughes et al. (Hughes & Ciulli, 2017)). The second class of molecules, known as molecular glues, intercalate between the E3 ligase and the targeted protein. This phenomenon was first described in plants as the mode of action (MoA) of the plant hormone auxin (Tan et al., 2007), and later described in humans with the breakthrough discovery of the MoA of the immunomodulatory inhibitory drug (IMiD) thalidomide that binds to the E3 ligase receptor CRBN (Ito et al., 2010). Instead of inhibiting the ligase, IMiD binding was accompanied by a gain-of-function phenotype through a molecular glue MoA leading to the recruitment, ubiquitination and subsequent degradation of Ikaros transcription factors (IKZF) (Krönke et al., 2014; Lu et al., 2014). Key structural insights about the IMiD binding mode and its ability to glue proteins to CRBN (Fischer et al., 2014; Petzold et al., 2016) paved the way for the use of IMiDs, both as handles for PROTACs (Lu et al., 2015; Winter et al., 2015) as well as molecular glues for *de novo* target proteins (Bonazzi et al., 2023) with first candidates in clinical trials (“A Phase 1/2 Trial of ARV-471 Alone and in Combination With Palbociclib (IBRANCE®) in Patients With ER+/HER2-Locally Advanced or Metastatic Breast Cancer,” ; “Study of Safety and Efficacy of DKY709 Alone or in Combination With PDR001 in Patients With Advanced Solid Tumors,”) (e.g. candidates ARV-471, trial NCT04072952 or DKY709, trial NCT03891953).

A common key asset for both classes of TPD molecules are binders to E3 ligases to enable recruitment of *de novo* substrates. Even though there are ca. 600 E3 ligases encoded in the human genome, until now only a handful have been successfully engaged with non-covalent binders for TPD. CRBN is the most widely used E3 which is hijacked by IMiD derivatives. The second most described ligase for TPD, the von Hippel-Lindau protein (VHL) (Buckley et al., 2012), is already less frequently used in PROTACs as well as in clinical development candidates compared to CRBN (Bondeson et al., 2015). In addition, to the CRBN and VHL E3 ligase binders that can mediate TPD at low and sub nM potencies (Bondeson et al., 2015; Krieger et al., 2023; Zorba et al., 2018), there are few additional non-covalent ligase binders that can match these potencies in TPD applications. Other promising non-covalent E3 ligase binders with characterized but less potent PROTACs for TPD of neo-substrates have been described for KEAP1 (Wei et al., 2021), MDM2 (Schneekloth et al., 2008), and IAP (Tinworth et al., 2019). In addition, multiple chemical proteomics approaches using electrophiles have enabled the discovery of additional tractable ligases for TPD (Pinch et al., 2022) such as DDB1 and CUL4 associated factor 16 (DCAF16) (Zhang et al., 2019), RNF4 (Ward et al., 2019) and more recently DCAF1 (also known as Vpr binding protein VprBP) (Tao et al., 2022). However, even though they are great tools to discover novel E3 ligases for TPD these chemical entities lack the full potential of the catalytic nature seen for non-covalent ligase binders since it is limited by the half-life of the ligase receptor. Furthermore, the covalent occupancy of the E3 ligase receptor might block degradation of natural substrates and as such may be more prone to on-target toxicity. A recent publication described reversible ligands binding to the WD40 domain of DCAF, highlighting further the potential of developing non-covalent DCAF1-based PROTACs(Li et al., 2023).

The most widely used IMiDs for TPD are thalidomide and its next-generation derivatives lenalidomide and pomalidomide, which have been approved as 1^st^ line treatment for multiple myeloma (MM). MM is a plasma cell malignancy that is dependent on IKZF transcription factors (reviewed in (Zuo & Liu, 2022)). Of note, thalidomide and its analogue lenalidomide have been approved by the FDA for the treatment of MM many years before their TPD MoA was described. Unfortunately, a recurring problem in IMiD-treated MM patients is emergence of resistance. About 30% of the IMiD-refractory patients bear alterations in CRBN levels, with copy loss in most cases, and lower CRBN levels result in reduced IMiD-mediated IKZF degradation. CRBN is a non-essential gene based on genome-wide CRISPR KO studies (Shalem et al., 2014; Tsherniak et al., 2017; Wang et al., 2014), which raises the question if loss of, or mutations in CRBN might be a liability for CRBN-targeting PROTACs?. Therefore, targeting essential E3 ligases could represent a potential strategy to circumvent or at least delay the emergence of therapy resistance (Hanzl et al., 2023).

Here, we describe the discovery of PROTACs based on a novel and specific binder to the E3 ligase receptor DCAF1 (Vulpetti et al. submitted). DCAF1 is an essential WD40 repeat (WDR) domain containing E3 ligase receptor of the Cullin RING ligase (CRL) 4 subfamily (Hrecka et al., 2007), and using two different PROTAC prototypes, one directed towards the nuclear protein BRD9 and the other towards tyrosine kinases, we demonstrate utility of DCAF1 for targeted protein degradation.

Finally, we extend our studies by using a specific Bruton’s tyrosine kinase (BTK) inhibitor to degrade BTK, a target protein with approved inhibitors to treat various blood-borne cancers (Shirley, 2022). We show that DCAF1 can be harnessed for BTK degradation in preclinical settings of resistance to CRBN PROTACs.

## RESULTS

### Discovery and characterization of a selective DCAF1 E3 ligase receptor binder

Successful discovery of ligands for WDR domain proteins such as EED (Huang et al., 2022) and WDR5 (Ding et al., 2023) (Guarnaccia et al., 2021) shows that WDR domains can be targeted (reviewed in (Schapira et al., 2017)). Therefore, amongst the 600 E3 ligases we focused on the 48 E3 ligases that contain WDR domains as putative recognition motifs (Supplementary Table 1). In addition, as a rationale to have potentially a higher bar for emerging resistance mechanisms on the ligase side, we analyzed this subset of 48 E3 receptors for essentiality by extracting the Demeter2 dependency scores from the DepMAP database (McDonald et al., 2017; Tsherniak et al., 2017).

Interestingly only three E3 ligase family members show pan-lethality upon gene depletion: DCAF1 (aka VPRBP), DDB1 and PRPF19 (Fig. 1A). Since many CRL E3 ligase receptors are well-characterized with known substrates and degradation activities such as the FBXW and DCAF subfamilies of CRL1 and CRL4 receptors, respectively, we focused on DCAF1, a CRL4 receptor with only one WDR domain, for ligand discovery to enable TPD of *de novo* substrates. Potentially the adaptor DDB1 would have also been an intriguing candidate, as it bridges CUL4A/B with the DCAF receptors. However, we felt that the three WDR domains of DDB1 and their positioning might not enable optimal target presentation for ubiquitination. Notwithstanding, the discovery of molecular glues for DDB1 targeting CDK12-cyclinK proves tractability of this adaptor (Słabicki et al., 2020). The Pre-mRNA processing factor 19 (PRPF19) belongs to the U-box family of E3 ligases with critical roles in DNA damage response (DDR) and splicing and was also shown to mediate lysine 63-linked ubiquitin chains (K63) (Song et al., 2010).

**Figure 1.**
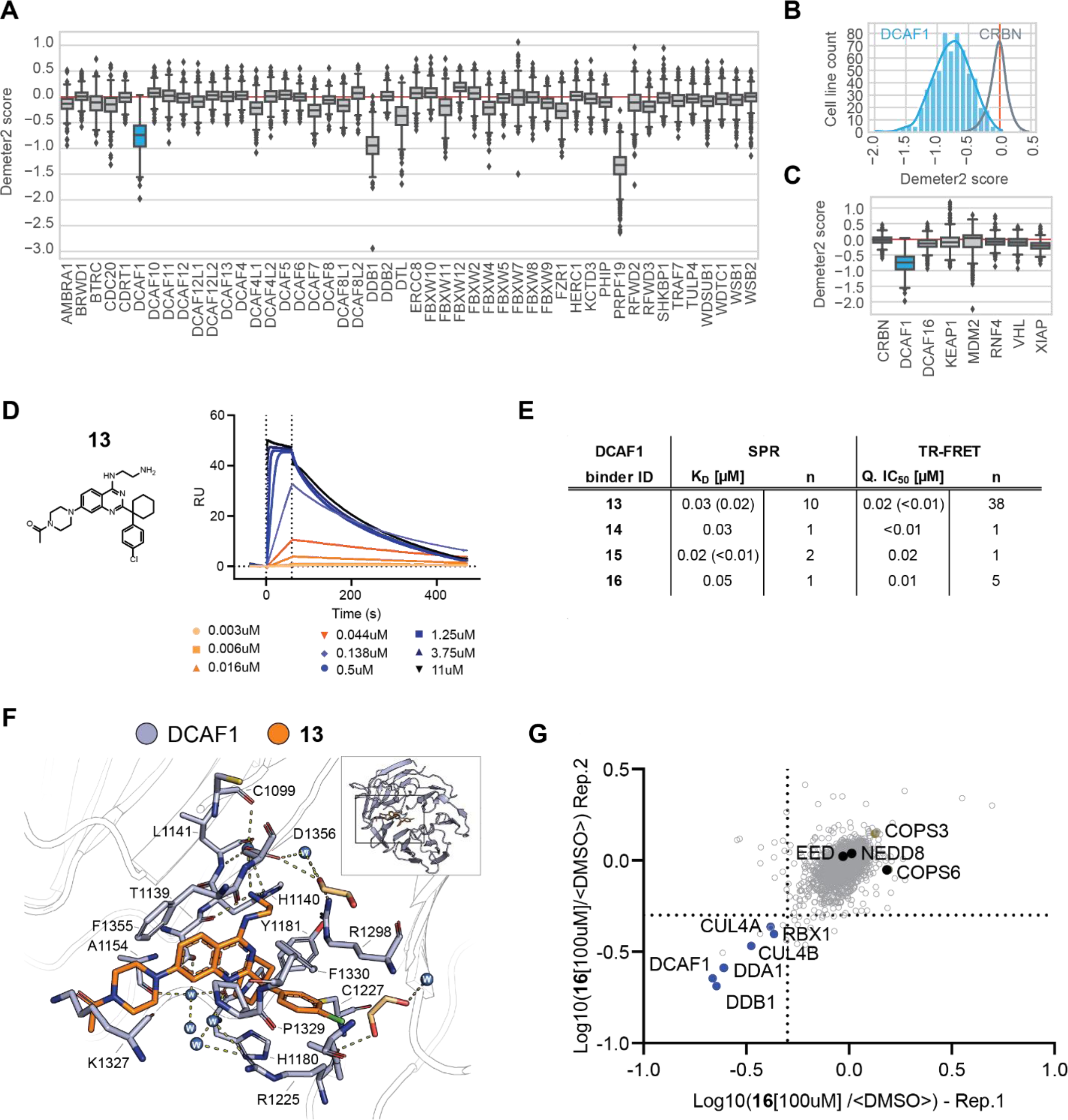
DCAF1 ligand characterization for targeted protein degradation. (A) RNAi gene effect score (Demeter2) extracted for WDR domain containing E3 ligases. (B) Histograms depicting Demeter2 scores per # of cell lines for CRBN (grey) and DCAF1 (blue). (C) Demeter2 scores for selected E3 ligases. (D) Structure of DCAF1 binder VHF543 and SPR sensorgrams with surface immobilized DCAF1(WDR) and (**13**) as analyte. (E) Affinity (KD) determined by SPR and IC50 determined by DCAF1 TR-FRET for selected DCAF1 binders (F) Binding mode of (**13**) (orange) in DCAF1 (blue, pdb ID: 8OO5). The inlet with the cartoon depiction of DCAF1 highlights the location of the binding site in the WD40 domain of DCAF1. Shown in the remaining part is detailed view of selected amino acids interacting with the compound. Hydrogen bonds are indicated as dashes. (G) Dot plot depicting competition from (**16**) beads of significant (**13**) interacting proteins determined by proteomics from 2 independent replicates 1 and 2. Dotted line depicts cut-off at 50% competition.

Interestingly, DCAF1 is frequently hi-jacked by Vpr and Vpx, virion-associated proteins encoded by lentiviruses, such as the human immunodeficiency virus (HIV) (Hrecka et al., 2007). Mechanistically they act as a protein-glue to recruit cellular restriction factors to DCAF1 for ubiquitination and subsequent degradation, setting a precedent for *de novo* substrate degradation via DCAF1 (Schwefel et al., 2014). Furthermore, DCAF1 has been shown to be a preferred receptor in Cullin4 ligase assemblies and ranks as the second most abundant CRL4 receptor in CRL4 E3 ligases behind CRBN (Reichermeier et al., 2020). In addition, various DCAF1 substrates have been proposed such as e.g. MCM10 (Kaur et al., 2012) or FOXM1 (Wang et al., 2017).

A potential lack of genetic dependency on CRBN in cancer cells as exemplified by the Demeter2 score might explain rapidly emergence of resistance in patients (Fig. 1B). Therefore, we hypothesized that the essentiality of DCAF1 might provide a higher bar for the occurrence of resistance. Furthermore, comparison of genetic dependency scores between DCAF1 and other ligase receptors that have been liganded for TPD approaches highlighted the potential of DCAF1 as an essential gene with respect to its Demeter2 score (Fig. 1C).

Our DCAF1 binder discovery strategy and medicinal chemistry campaign is described in an accompanying manuscript (Vulpetti et al. submitted). In summary, WDR binders from a previous hit-discovery campaign for EED (Huang et al., 2022) have been screened and optimized using NMR, computer-aided drug design (CADD), biophysics and structure activity relationship (SAR) approach. This resulted in the discovery of a scaffold that occupies the DCAF1 WDR donut-hole pocket (Vulpetti et al. submitted). Here, we further characterized the potent DCAF1 binder (**13**), which binds to DCAF1 with a Kd by SPR of <50 nM (Fig. 1D and E), an X-ray structure confirmed the same binding mode in the WDR donut-hole and helped to determine the piperazine as a potential site for an exit vector (Fig. 1F). Indeed, extension by PEG3 or PEG6 chains (cpd (**14**) and (**15**), Supplementary Fig. 1A) did not alter binding to DCAF1 (Fig. 1E) and X-ray crystallography confirms overlapping binding to the DCAF1 donut-hole pocket like the parental binder (**13**) and highlighted the extension of the PEG chain and accessibility outside of the pocket (Supplementary Fig. 1B) allowing functionalization of this DCAF1 binding scaffold for synthesis of proteolysis targeting chimeras (PROTACs) and further compound specificity studies. To determine cellular compound binding specificity, we coupled compound (16) to beads using azide-alkyne click-chemistry (Supplementary Fig. 1C) and show that alkyne derivatization of (**13**) (resulting in cpd (**16**)) does not alter DCAF1 binding (Fig. 1E). 293T cell lysate incubation with these (**16**)-beads and comparison to beads in competition with (**16**) to block specific compound interacting protein complexes from binding, enabled the discovery of the whole core CRL4^DCAF1^ complex including CUL4A/B, DDB1, DDA1 and RBX1 (Fig. 1G and Supplementary Table 2). Interestingly, neither EED whose early scaffolds led to the discovery of cpd (**13**) nor any other WDR containing protein (besides DDB1, a known DCAF1 interactor) showed significant depletion >50% from beads, proving high-specificity of our DCAF1 binding scaffold.

### Generation and chemical and in-depth genetic validation of a prototype DCAF1-BRD9 PROTAC

Since DCAF1 is mainly localized to the nucleoplasm (*The Human Protein Atlas-DCAF1*; Thul et al., 2017; Uhlén et al., 2015), we wanted to test a first DCAF1 PROTAC against a nuclear target that has been successfully degraded with a TPD MoA. The non-canonical BAF (ncBAF) complex member BRD9 has been shown to be an attractive therapeutic target for e.g. synovial sarcoma (Brien et al., 2018; Michel et al., 2018) and has been shown to be amenable for PROTAC-mediated degradation hi-jacking CRBN (Remillard et al., 2017). Here we synthesized a prototype DCAF1-BRD9 PROTAC (**DBr-1**) by coupling the published BRD9 bromodomain binder BI-9564 (Martin et al., 2016) via a piperidine and aliphatic carbon linker to the piperazine of our DCAF1 binding scaffold (**13**) resulting in **DBr-1** (Fig. 2A). Using SPR we confirmed binding of **DBr-1** to DCAF1 with a steady-state Kd of 209 nM, which is approximately 2-fold higher compared to the DCAF1 binder (**13**) having a K_d_ of 93 nM. Furthermore, we show ternary complex formation between DCAF1, **DBr-1** and BRD9 by SPR, observing a typical bell-shaped behavior when reaching saturating ligand concentrations, a “hook-effect” consistent with respective bimolecular interactions dominating at high **DBr-1** concentrations (Douglass et al., 2013) (Fig. 2B).

**Figure 2.**
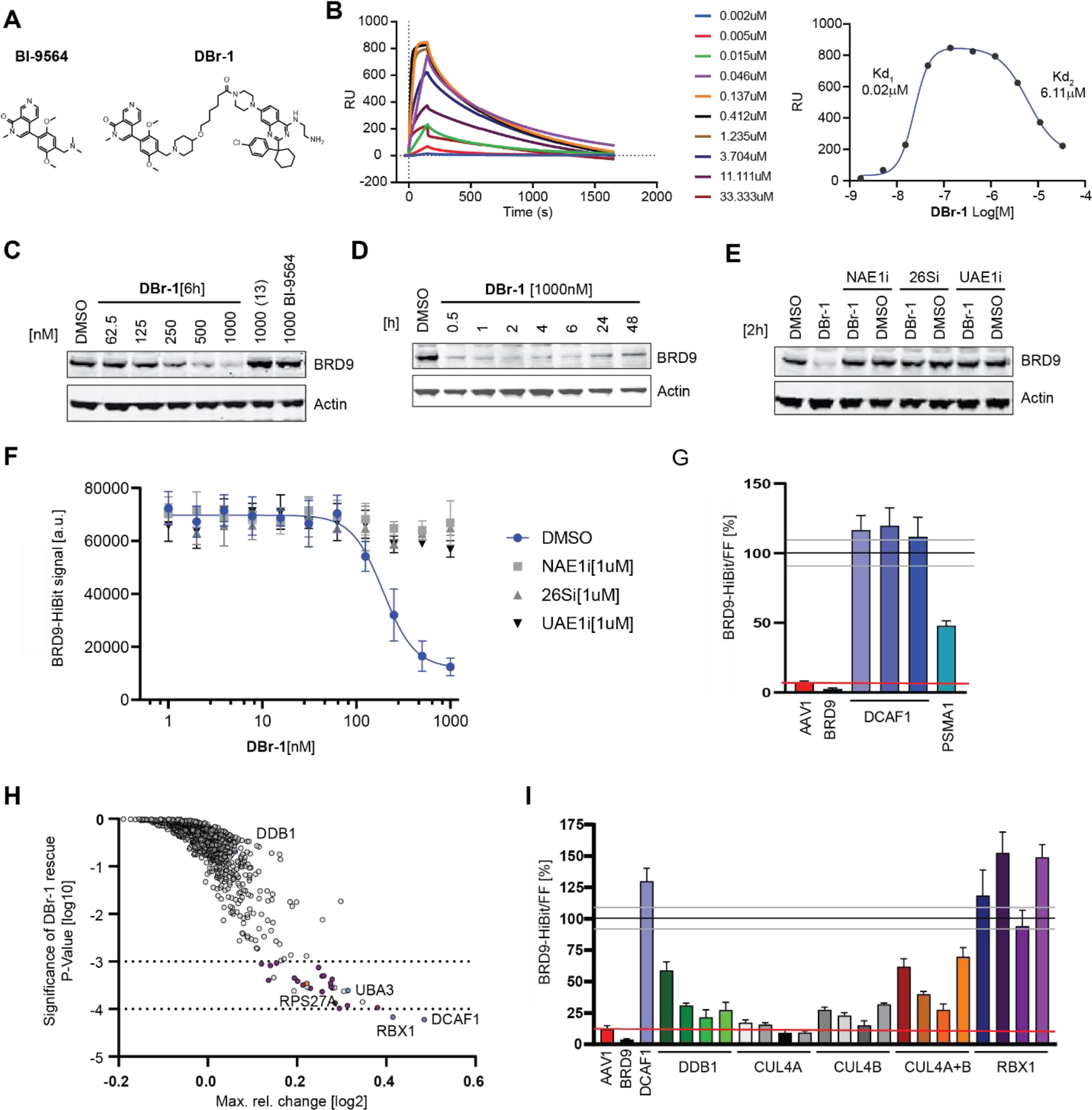
DCAF1-BRD9 PROTAC characterization and in-depth genetic validation. A) Compound structures of BRD9 BD binder BI-9564 and corresponding DCAF1-BRD9 PROTAC (**DBr-1**). B) SPR sensorgrams with surface immobilized DCAF1(WDR) and (**DBr-1**) as analyte in the presence of 0.2uM BRD9-BD(130-250) (left panel) and response blotted against (**DBr-1**) concentration (right panel). Displayed values indicate apparent EC50 values of the two separated sigmoidal transitions, respectively. C) Immunoblot analysis of HEK293 BRD9-HiBit/FF/CAS9 cells treated for 6h with (**DBr-1**) at various doses as well as (**13)** and BI-9564 at 1000nM. D) Immunoblot analysis of HEK293 BRD9-HiBit/FF/CAS9 cells treated with 1000nM (**DBr-1**) for various time points. E) Immunoblot analysis of HEK293 BRD9-HiBit/FF/CAS9 cells pretreated with NEDD8 E1 inhibitor (NAE1i) [1000nM], proteasome inhibitor Bortezomib (26Si) [1000nM] and Ubiquitin E1 inhibitor (UAE1i) [1000nM] for 2h, followed by (**DBr-1**) treatment [1000nM] for additional 2h. F) BRD9-HiBiT signal detection of samples pretreated as in (E), followed by 2h (**DBr-1**) treatment at various doses, DC50 = 193nM. Data represents mean ± SEM from n=4 replicates. G) Relative BRD9-HiBit vs Firefly signal ratio normalized to non-treated DMSO ctrl after 2h of (**DBr-1**) treatment [1000nM]. Indicated gene editing with sgRNA has been performed as described 6d before treatment. Data represents mean ± SEM from n=4 replicates. H) Ubiquitin sgRNA sublibrary rescue scores from (**DBr-1**) treatment plotted as significance of rescue P-value (y-axis) vs max. rel. change. Dotted lines at -4 and -3 P-value indicate strong and weaker hits with a false-discovery rate of 7%. I) Relative BRD9-HiBit vs Firefly signal ratio normalized to non-treated DMSO ctrl after 2h of (**DBr-1**) treatment [1000nM]. Indicated gene editing with individual sgRNAs or a combination of two guides for *CUL4A* and *CUL4B (CUL4A+B)* has been performed as described 6d before treatment.

At 1000 nM **DBr-1** exhibited about 90% BRD9 protein loss after 6h incubation of HEK293 cells, and thus sub-µM potency as estimated from a dose-response curve (DC_50_ approx. 500nM). No BRD9 protein reduction was observed incubating the individual binders at highest concentration for either DCAF1 WDR site (**13**) or the BRD9 BD site **BI-9564** (Fig. 2C). A time course with a fixed concentration of 1000 nM **DBr-1** showed rapid degradation (90% degradation already after 30 min) that is maintained for more than 6 hours with slow rebound after 24h and 48h hours (Fig. 2D). We observe full rescue of DBr-1 mediated degradation using either a chemical inhibitor against the ubiquitin-activating E1 (UAE1i, TAK-243), the Nedd8-activating E1 (NAE1i, MLN4924) or a proteasome inhibitor (26Si, bortezomib) (Fig. 2E), demonstrating specificity for the ubiquitin-mediated CRL proteasome pathway. To further confirm and study the underlying ubiquitin pathway components that are hi-jacked by **DBr-1**, we engineered CAS9 and Firefly luciferase expressing HEK293 cells that have been engineered at the BRD9 locus to express BRD9-HiBIT (BRD9-HiBIT CAS9/FF) with the aim to enable sgRNA mediated knock-out studies and its impact on BRD9-HiBit levels post **DBr-1** treatment. Incubating these BRD9-HiBit CAS9/FF cells for 2 hours with **DBr-1** confirmed efficient degradation with a DC_50_% of 193 nM. Rescue of BRD9-HiBit levels with NAE1i, 26Si and UAE1i confirmed degradation specific loss-of-signal (Fig. 2F). Using 3 combinations of two individual small guide RNA (sgRNA) against *DCAF1*, one ctrl sgRNA against *BRD9*, one sgRNA against a proteasome pathway ctrl gene *PSMA1* as well as a nonspecific sgRNA *AAV1,* we genetically confirm full rescue with all 3 *DCAF1* sgRNA mixes and as such dependence on DCAF1 as the E3 ligase receptor, as predicted (Fig. 2G).

To obtain a more complete picture of ubiquitin pathway components involved in **DBr-1** mediated degradation of BRD9, we performed a CRISPR knock-out rescue screen with an arrayed ubiquitin-sublibrary (943 target genes, 3-4 pooled sgRNAs per gene, Thermo Fisher, Supplementary Table 3). The **DBr-1** rescue screen was performed as outlined in Supplementary Fig. 2A. In summary, genetic BRD9-HiBit degradation rescue was measured after a short 2h pulse of **DBr-1** in HEK293 BRD9-HiBIT CAS9/FF cells, 5 days after gene editing by sgRNAs. For hit calling we set a threshold based on a minimal Firefly signal > log10(1.25), and this threshold was based on signal measurements of control wells containing non-infected cells selected with puromycin. The Firefly signal served as control for well-based variation of cell number and was used to normalize the BRD9-HiBit signals on a plate basis using a non-linear regression 3^rd^ order polynominal curve (Supplementary Fig. 2B – E). From this normalized data we calculated a robust z-score per gene (Supplementary Figure F) to feed into a gene-level RSA statistical mode (Zeng et al., 2019) to calculate gene effect scores as max. rel. change and significance P-values (Supplementary Table 4). We could confirm sgRNAs against *DCAF1* as the strongest rescuer, in addition we found *RBX1* the RING-box protein 1 a key component of the CRL4^DCAF1^ complex as another strong hit (Fig. 2H). DDB1 and CUL4A and B, two other key components of the CRL4^DCAF1^ E3 ligase did not rescue from **DBr-1** degradation in our screen setting. CUL4A and B did not pass the Firefly signal threshold together with 117 other genes (including PSMA1 we previously used as ctrl, Supplementary Fig. 2F). For further hit-calling, we defined a threshold P-value of < log10(−3), based on a randomized activity run (Supplementary Fig. 2G, STAR Methods). Based on this threshold, we discovered 30 genes that rescued from **DBr-1** mediated degradation with a false discovery rate (FDR) of 7% (Supplementary Table 5). In contrast, all the genes that did not pass the Firefly signal threshold did not pass the P-value threshold of < log10(−3) as opposed to all the plate-based *DCAF1* ctrl sgRNAs (Supplementary Fig. 2F). Interestingly, *UBA3* and NAE1, as well as 20 proteasomal subunit gene knockouts rescue **DBr-1** mediated degradation as previously confirmed with chemical inhibitors NAE1i and 26Si (Supplementary Table 5 and Fig. 2H). Another interesting hit was also *RPS27A*, the ubiquitin and ribosomal protein S27A gene, that expresses a fusion protein between RPS27A and ubiquitin. *RPS27A* is one of the cellular sources of ubiquitin expression, together with *UBB* (P-Val log10(−1.2))*, UBC* (P-Val log10(−0.6)) and *UBA52* (P-Val log10(−2.1 (Luo et al., 2022; Redman & Rechsteiner, 1989). Based on this data and the redundant nature of the cellular ubiquitin sources we hypothesize that in these HEK293 cells *RPS27A* is the main source of ubiquitin expression.

To validate RBX1 and follow up on the other CRL4^DCAF1^ core components CUL4A, CULB and DDB1 that did not score in the rescue screen, we designed 4 sgRNAs per gene to test for rescue individually or in the case of the redundant CUL4A and CULB (Sang et al., 2015) also in combination. We could confirm full rescue from **DBr-1** degradation with all four individual sgRNAs against *RBX1*; for *DDB1*, we saw only partial rescue with 1 guide >50%. Interestingly, only combination of *CUL4A* and *CUL4B* targeting guides led to rescue >50%, confirming the redundant nature of these two Cullins (Fig. 2I). In summary, with these chemical and in-depth genetic rescue experiments we present CRL4^DCAF1^ and proteasome-mediated degradation with our first sub-µM DCAF1-BRD9 PROTAC **DBr-1** and highlight that the E3 ligase receptor DCAF1 can be specifically targeted to mediate degradation of the nuclear protein BRD9.

### DCAF1 PROTACs mediate degradation of tyrosine kinases

After successfully degrading BRD9, a nuclear BD-containing protein, we explored whether DCAF1 PROTACs could also mediate degradation of kinases. To cast a wider net, we synthesized another prototype PROTAC targeting the tyrosine-kinase subfamily by coupling dasatinib (Lombardo et al., 2004) to our DCAF1 binder using a similar piperidine aliphatic carbon linker as for BRD9 (Fig. 3A). Dasatinib binds to >30 kinases (Rix et al., 2007) and has been shown to inhibit (Kitagawa et al., 2013) multiple tyrosine kinases of the ABL, DDR, TEC (tyrosine kinase expressed in hepatocellular carcinoma), SRC (sarcoma kinase) and EPH subfamilies at low nM concentrations. Interestingly, amongst these kinase targets, we find kinases that preferentially localize to the plasma membrane (tyrosine receptor kinases EPH and non-receptor tyrosine kinases TEC and LYN), nucleoplasm (ABL1), cytosol (CSK) or cytosol and nucleoplasm (e.g. LIMK2) (*The Human Protein Atlas*; Thul et al., 2017), further enabling us to study if DCAF1 shows particular preferences towards cellular localization of its targets. We first tested our DCAF1-Dasatinib PROTAC by Western blotting against four non-receptor tyrosine kinase candidates C-Src Tyrosine Kinase (CSK), Lck/Yes-Related Protein Tyrosine Kinase (LYN), LIM domain kinase 2 (LIMK2) and Abelson Protein Tyrosine kinase 1 (ABL1). Interestingly, we observed significant reduction in protein levels for CSK and LIMK2, but not for LYN or ABL1 after 6h in a proteasome (26Si) and CRL dependent manner (NAE1i), neither the individual DCAF1 binder nor dasatinib impacted protein levels (Fig. 3B). Compared to a prototype CRBN-dasatinib degrader **CDa-1** that shows degradation of all the four probed kinases, we also see reduced potencies towards the two degraded kinases for the DCAF1 PROTAC **DDa-1**compared to the CRBN PROTAC **CDa-1**. However, we cannot conclude that recruiting tyrosine kinases to CRBN leads to better degradation based on only two prototype PROTACs, and with only limited exploration of linkerology carried out for both.

**Figure 3.**
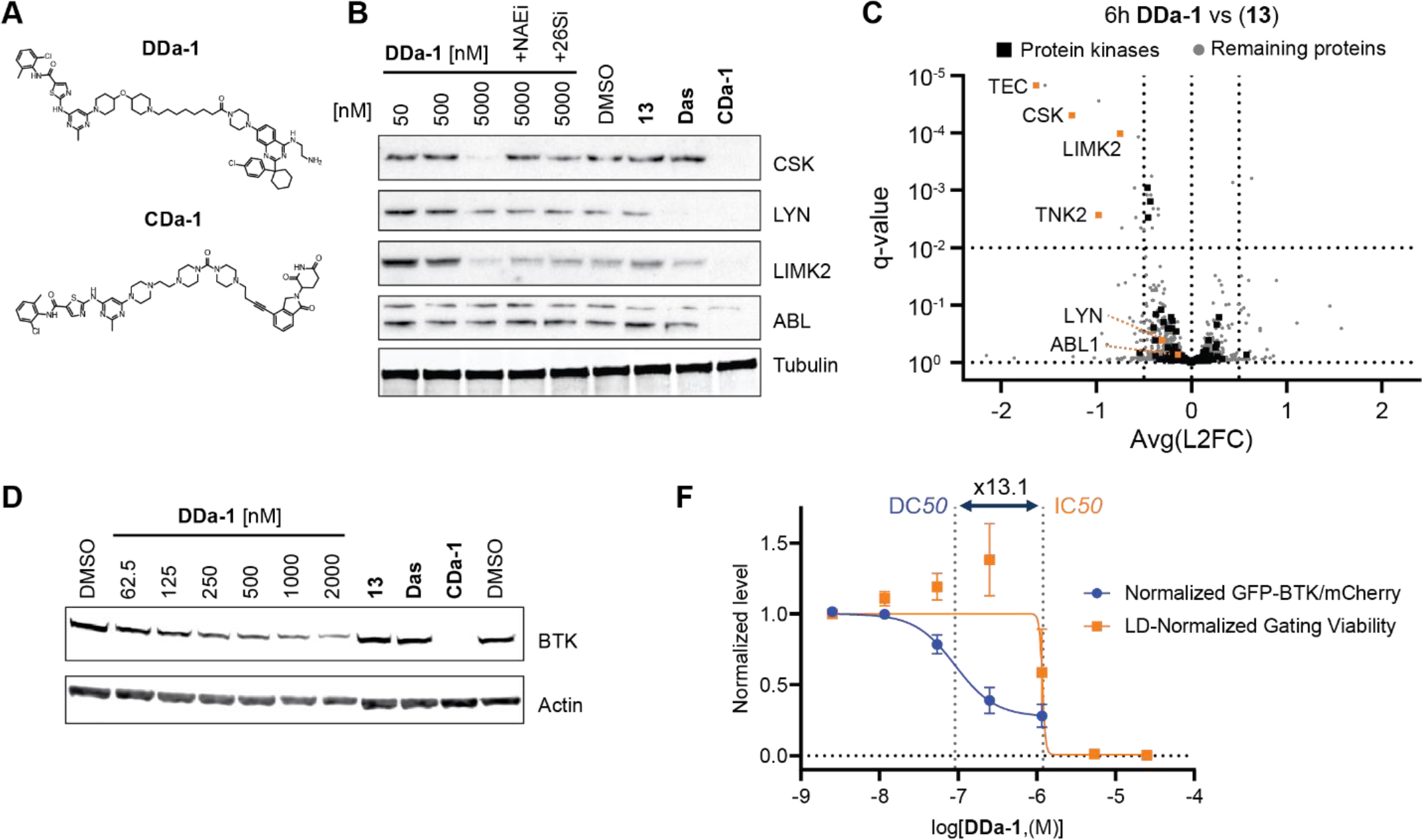
DCAF1-Dasatinib PROTAC degrades multiple tyrosine kinases. A) Compound structures of the DCAF1-Dasatinib PROTAC (**DDa-1**) and a CRBN-Dasatinib ctrl PROTAC (**CDa-1).** B) Immunoblot analysis of HEK293T cells treated for 6h with (**DDa-1**) at 50, 500 and 5000 nM as well as (**13**) and Dasatinib (**Das**) at a dose of 5000nM as control. Indicated co-treatments with NEDD8 E1 inhibitor (**NAE1i)** [1000nM] and proteasome inhibitor Bortezomib (**26Si**) [1000nM] are indicated. CRBN-Dasatinib degrader (**CDa-1**) [50nM] served as an internal ctrl degrader. C) Whole proteomics profile comparing differential protein levels between DCAF1-Dasatinib PROTAC (**DDa-1**) vs DCAF1 binder (**13**) plotted as Avg log2 fold change (L2FC, x-axis) versus Avg statistical significance P value (pval, y-axis). Horizontal dotted line indicates pval 10^−2^, vertical lines indicate a log2 fold change of 0.5 D) Immunoblot analysis of TMD8 cells treated for 24h with (**DDa-1**) at various doses as well as (**13**) and Dasatinib (**Das**) at a dose of 2000nM as control. CRBN-Dasatinib degrader (**CDa-1**) [50nM] served as an internal ctrl degrader. E) Degradation of BTK-GFP in TMD8 BTK-GFP/mCh cells after 24h (**DDa-1**) treatment displayed as normalized rel. change of the ratio between BTK-GFP and mCherry (mCh) signals (blue curve), DC_50_ = 0.09 µM. Viability after 24h (**DDa-1**) treatment is displayed as rel. change in cellular distribution between viable and apoptotic FSC/SSC gate, GI_50_ = 1.20 µM. The viability window as ratio between GI_50_/ DC_50_ = 13.1 is indicated with dotted vertical lines.

To obtain insights into global protein abundance changes, we analyzed HEK293 cells after 6h of treatment with **DDa-1** by LC/MS-based proteomics. We were able to confirm downregulation of the two previously studied kinases CSK and LIMK2 and further discovered two additional non-receptor tyrosine kinases, TEC and TNK2, which showed significant protein level reduction (Fig. 3C and Supplementary Table 6). Interestingly both, TEC and TNK2 are reported to preferentially localize to the plasma membrane (*The Human Protein Altlas-TNK2*; *The Human Protein Atlas-TEC*; Thul et al., 2017).

Bruton’s tyrosine kinase (BTK), another TEC family member, has been successfully targeted in a variety of blood cancers with the covalent BTK inhibitor ibrutinib in the clinic (Alu et al., 2022; Ran et al., 2022; Wen et al., 2021). Furthermore, BTK has been shown to be an attractive target for PROTAC mediated degradation hi-jacking the ligase CRBN (Buhimschi et al., 2018; Zorba et al., 2018) and with advanced TPD molecules in clinical trials (such as e.g. NX-2127, (Mato et al., 2022)). Since BTK is not expressed in HEK293 cells but has been shown to be bound and inhibited by dasatinib (Hantschel et al., 2007; Rix et al., 2007) we wanted to test if our dasatinib-DCAF1 PROTAC is active against this kinase. To this aim, we probed the BTK-expressing diffuse large B-cell lymphoma (DLBCL) cell line TMD8 (Tohda et al., 2006) with **DDa-1** at various concentrations and could confirm BTK degradation with sub-µM potency (Fig. 3D).To further explore BTK degradation in a more quantitative higher-throughput profiling system we engineered TMD8 cells stably expressing bicistronic BTK-GFP/mCherry construct (referred to as TMD8 BTK-GFP/mCh), ratiometric analysis of GFP to mCherry signal ratio changes allows a quantitative measurement of degradation and uncoupling of degradation from transcriptional and translational effects at equilibrium (Emanuele et al., 2011; Yen et al., 2008). TMD8 DLBCL cells have been shown to be dependent on BTK and hence sensitive to BTK inhibition (Kuo et al., 2016) and degradation (Lim et al., 2023).

To ensure that our engineered TMD8 BTK-GFP/mCh cells still respond to BTK perturbations, we used a BTK inhibitor BTKi (**17**) (WO2019/186343 A1) and compared CYLD tumor suppressor accumulation (Young & Staudt, 2012). Reassuringly, parental TMD8 and engineered TMD8 BTK-GFP/mCh cells gave a very similar biological response to BTK inhibition after 24 hours (Supplementary Fig. 3A).

Considering the sensitivity of TMD8 BTK-GFP/mCh cells to BTK perturbations, we explored potential signal changes upon impacts on cellular fitness to ensure confidence in measuring BTK degradation. We observed that the TMD8 BTK-GFP/mCh cells show some response to on-target BTK inhibition when measuring forward and sideward scattering by flow cytometry (FSC and SSC) of single cells after BTKi (**17**) treatment within the 24h timeframe we use to measure degradation effects. At very high BTKi (**17**) concentrations [25µM] we observe significant population shifts in FSC and SSC. By using two apoptotic markers for early and late apoptosis we could confirm that the apoptotic cells almost exclusively localize to the FSC/SSC-shifted population (Supplementary Fig. 3B). Therefore, our gating strategy to score for degradation focused on the viable cell population as defined by the FSC/SSC-scatter population observed in DMSO-treated cells. In addition, we set a minimal viability threshold of 25% viable cells and excluded treatments from degradation calculation due to the low number of viable cells observed (Supplementary Table 7, “Max dose w/ Gating Viability >25%”).

Having established this additional cell viability read-out when measuring BTK-GFP degradation allowed us to calculate a parameter to confidently score degradation. Plotting degradation as change of BTK-GFP/mCherry signals and viability as change in viable to apoptotic cells allows calculation of the fold change of viability GI_50_ *versus* degradation DC_50_ (Supplementary Table 7, “V/D [GI_50_ Viability / DC_50_ Degradation]”) and distinguish from on-target (as seen for BTKi (**17**)), as well as potential off-target effects on cellular fitness that could impair the degradation read-out especially at toxic concentrations. BTKi (**17**) has a V/D score of 0.3 whereas the very potent degrader **CDa-1** has a score of 2124.8; when testing our DCAF1 binder (**13**) we observed loss of viability at concentrations of >5µM with a V/D score of 0.2 (Supplementary Table 7, Supplementary Fig. 3C and D). Finally, testing **DDa-1** in this system validated our findings on endogenous BTK degradation (Figure 3E, Supplementary Table 7: DC_50_=0.09 µM and V/D=13.1).

### Discovery of a potent BTK-specific DCAF1 PROTAC

Having established the BTK-GFP/mCh TMD8 cell system to measure BTK degradation and shown successful BTK degradation via DCAF1 using a dasatinib-based PROTAC, we investigated the potential to develop more specific DCAF1-directed BTK PROTACs. To this end, we synthesized three DCAF1 PROTACs with our specific BTK binder (**17**) using PEG-based linear linkers (**DBt-3-5**, Fig. 4A). All PROTACs retained affinity to DCAF1, although with ca. 1.3 - 10 fold reduced IC50 for **DBt-3** and **DBt-4** compared to (**13**). However, the presence of BTK increased the binding potency towards DCAF1 for these two compounds, suggesting ternary complex formation facilitated by cooperativity (α>1, calculated as α = IC50^DCAF1^/IC50^DCAF1/BTK^) (Fig. 4B, Supplementary Fig. 4A). In contrast, **DBt-5** with a longer linker showed no impaired binding to DCAF1 compared to (13) and the presence of BTK decreased the potency for DCAF1 binding.

**Figure 4.**
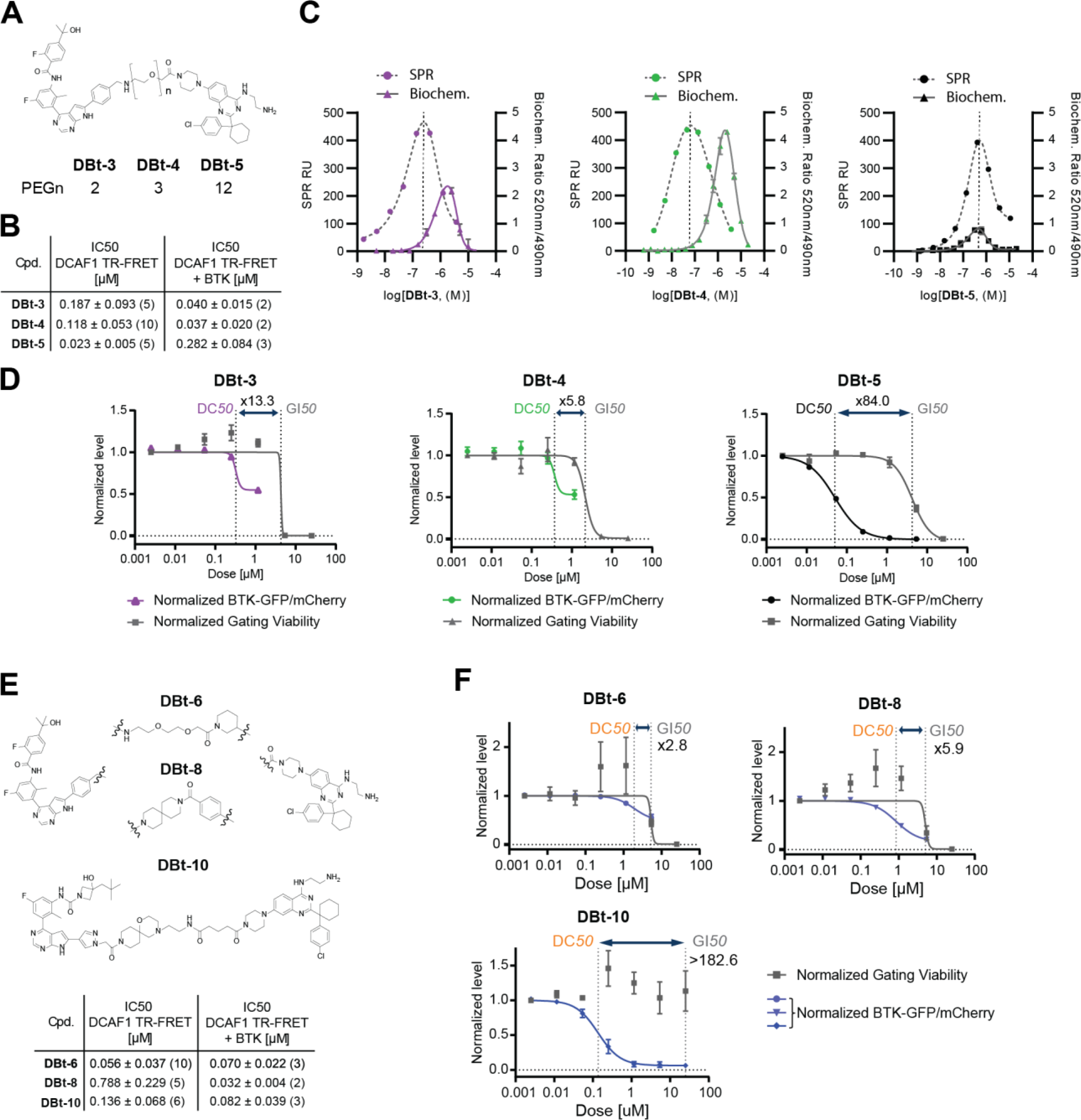
Characterization and discovery of DCAF1-BTK PROTACs. (A) Chemical structures of BTK- and DCAF1 binder with the respective linkers of **DBt-3-5**. (B) DCAF1 TR-FRET and DCAF1 TR-FRET + BTK assay (Supplementary Fig. 4A) results for **DBt-3-5**. Values displayed represent the average and standard deviation of multiple experiments as indicated by the number in brackets. (C) Overlay of SPR ternary complex formation assays (dashed lines, left y-axis) with biochemical Ubiquitination rates of DCAF1 measured in ubiquitin-transfer based TR-FRET assay (right y-axis). (D) BTK degradation and cellular viability (dashed lines) assessed in BTK-GFP/mCh TMD8 cells after 24 hours. The data points have been normalized to DMSO controls and each point represents the average and standard deviation of three independent experiments in triplicates. (E) Chemical structures of the linker region of the DCAF1-BTK PROTACs **DBt-6**, **DBt-8** and **DBt-10** with their respective number (n) of PEG units in their linker regions. Table displays DCAF1 TR-FRET and DCAF1 TR-FRET + BTK assay results for **DBt-6**,**DBt-8** and **DBt-10**. Values displayed represent the average and standard deviation of multiple experiments as indicated by the number in brackets. (F) BTK degradation assessed in GFP-BTK/mCherry TMD8 cells after 24 hours. The data points have been normalized to DMSO controls and each point represents the average and standard error of the mean three independent experiments in triplicates. The viability window as ratio between GI_50_/ DC_50_ is indicated with dotted vertical lines.

Nevertheless, we could confirm ternary complex formation between DCAF1 and BTK with all three compounds (**DBt-3-5)** using SPR (Fig. 4C, Supplementary Fig. 4B). The resulting data displayed a typical bell-shaped curve attributed to the “hook effect”, as expected for bidentate molecules (Casement et al., 2021), with all PROTACs displaying similar maximal signals, but for the shorter PEG linkers of **DBt-3/4** the maxima occurred already at 2-5 fold lower PROTAC concentrations. Despite this slightly improved ternary complex formation, the ubiquitination rate measured by time-resolved fluorescence energy transfer (TR-FRET) remained similar for all three compounds in an *in vitro* CRL4^DCAF1^ reconstituted BTK ubiquitination system (Fig. 4C, Supplementary Fig. 4B). Interestingly, **DBt-5** harboring a PEG-12 linker, showed the maximal ubiquitination rate, at approx. 5-fold lower concentrations, albeit at a slower rate compared to the shorter PEG linkers (Fig. 4C). This could indicate the importance of other factors for successful ubiquitination, such as accessibility of lysine residues in addition to ternary complex formation, which might be amenable through a longer more flexible linker enabling more poses between BTK and DCAF1.

The results of the *in vitro* assays were corroborated by BTK degradation in TMD8 BTK-GFP/mCh cells. In agreement with the ubiquitination results, the two shorter PEG linkers exhibited a similar degree of partial BTK degradation, while the PEG-12 analogue **DBt-5** displayed strong degradation with a DC_50_ value 0.055µM with a >80 fold viability window (Fig. 4D, Supplementary table 7). The compound induced cytotoxicity in the single-digit µM range after 24h, which was comparable between the three PEG analogues (Fig. 4D, Supplementary table 7). Intriguingly, **DBt-5** was able to potently bind to BTK in cells as measured by a cellular target engagement assay with an IC50 value of 0.39 µM and 0.22 µM in digitonin (DIG) permeabilized cells to overcome any permeability effects (Supplementary Fig. 4C) (Poller et al., 2022). The shorter linker containing **DBt-3** displayed lower cellular BTK affinity in both intact and DIG permeabilized cells (2.40 and 0.92µM, respectively). The difference in potency in permeabilized cells compared to biochemical enzymatic assays could potentially be attributed to the full length BTK, which could impair binding of PROTAC molecules bearing a shorter linker. The surprisingly good cell permeability of **DBt-5** might be positively influenced by active transport mechanisms of PEG-linked bifunctional molecules via IFITM genes (Lou et al. (2022), although further studies are needed to assess their impact in this case.

Next, we investigated if more complex linkers or change of the BTK binding moiety could provide alternative PROTAC molecules with similar or better degradation efficiency than **DBt-5**. Therefore, we designed 3 new molecules **DBt-6 DBt-8** and **DBt-10** (Fig. 4E), and thoroughly tested them with the previously described assays (Figure 4E, Supplementary Table 7,).

Compounds **DBt-6** and **DBt-10** incorporating slightly more complex linkers showed comparable affinity towards DCAF1 (Fig. 4E). However, the rigid linker of **DBt-8** reduced affinity for DCAF1 (>4 fold). Addition of BTK to the DCAF1 TR-FRET assay was tolerated without a significant change of DCAF1 affinity for **DBt-6** and **DBT-10 but** improved the measured affinity of **DBt-8** for DCAF1 by >20fold. Despite these differences, SPR ternary complex formation assays showed similar stable DCAF1-PROTAC-BTK complexes (Supplementary Fig. 4D). The rigid linker containing **DBt-8** as well as **DBt-10** with an alternative BTK binder displayed higher maximal RU values at similar concentrations compared to **DBt-5**, indicating that the altered linker design yielded in improved ternary complex stability. However, despite this improvement in ternary complex formation, the more complex linkers of **DBt-6-10** resulted in reduced cellular target engagement of BTK than **DBt-5**, which could be only improved for **DBt-10** by permeabilizing the cells (Supplementary Fig. 4C). While the latter suggested impaired cellular permeability of **DBt-10**, the weaker affinity for BTK of the **DBt-6** and **DBt-8** was additionally corroborated by enzymatic inhibition assays (supplementary Fig. 4E, Supplementary Table 7).

Interestingly, this lack of binary affinity with BTK or DCAF1 alone, but potent ternary complex formation in particular of **DBt-8** pointed towards a large degree of cooperativity of this interaction, which was supported by the TR-FRET assay ± BTK. The cooperativity factor αof the 6 PROTAC molecules correlated with a relative decrease in DCAF1 binary affinity compared to their parent compound **13** (Supplementary Fig. 4F). This suggests that compound activity itself was positively influenced by the initial binary complex formation, and the cooperativity might therefore not involve extensive protein-protein interaction. However, further structural studies both on the compound, as well as on the ternary complex would be necessary to clarify this.

Design of compounds with a high degree of cooperativity and prolonged ternary complex half-life has recently been suggested as a promising strategy to enhance PROTAC efficacy (Hu & Crews, 2022; Rosenberg et al., 2023; Roy et al., 2019; Yu et al., 2021). Intriguingly when testing for BTK degradation in the TMD8 BTK-GFP/mCherry cells, **DBt-10** outperformed **DBt-8** as well as **DBt-6** and showed comparable degradation efficacy to **DBt-5** (DC_50_ of 0.149 vs. 0.050µM) with an improved viability (>182 vs. 84×) window providing an DCAF1-BTK degrader with an optimized linker (Fig.4F).

However, the two most potent DCAF1-based BTK degraders **DBt-5** and **DBt-10** showed only modest cooperativity (<1 and 1.6) and while **DBt-10** induced a relatively stable DCAF1-BTK-ternary complex (t_1/2_ = 671s), the half-life of the **DBt-5** ternary complex was over 6 times shorter (Supplementary Fig. 4G). This less stable complex of **DBt-5** could be compensated by better cell permeability and better accessibility of lysine residues due to the longer and more flexible linker length compared to **DBt-10**.

Therefore, this small subset of 6 degraders highlights the multifactorial process of optimizing compounds, which must take into account not only the binary potencies and ternary complex formation, but also the accessibility of lysine residues and permeability of the PROTAC molecules.

### Potent DCAF1-BTK degrader enables BTK degradation in settings resistant to CRBN-BTK degraders

Since **DBt-10** showed superior degradation and the best degradation-viability score of the DCAF1-BTK PROTACs tested (V/D > 182.6 after 24h treatment), we wanted to assess if **DBt-10** is a DCAF1-specific BTK degrader suitable to further study in a CRBN-BTK PROTAC resistant setting by switching to an alternative ligase.

We confirmed the high viability score of **DBt-10** by measuring early and late apoptotic cells (Supplementary Fig. 5A), and after 24h we did not observe a strong induction of apoptosis as compared to BTKi (**17**) (Supplementary Fig. 3B), potentially underlying different cell permeability and kinetic properties of inhibition *versus* degradation. We observed efficient degradation of endogenous BTK in TMD8 cells at around 6 hours (Figure 5A) whereas control (**13**) treatment did not reduce BTK at any time points (Supplementary Fig. 5B). A dose response at the 6h time point showed that we drive significant reduction of BTK at a concentration of 2.5 µM (Figure 5B), a concentration and time point we used for a chemical rescue experiment using 26Si, NAE1i and UAE1i to validate proteasome, CRL and ubiquitination dependency of **DBt-10** mediated BTK degradation, respectively (Figure 5C).

**Figure 5.**
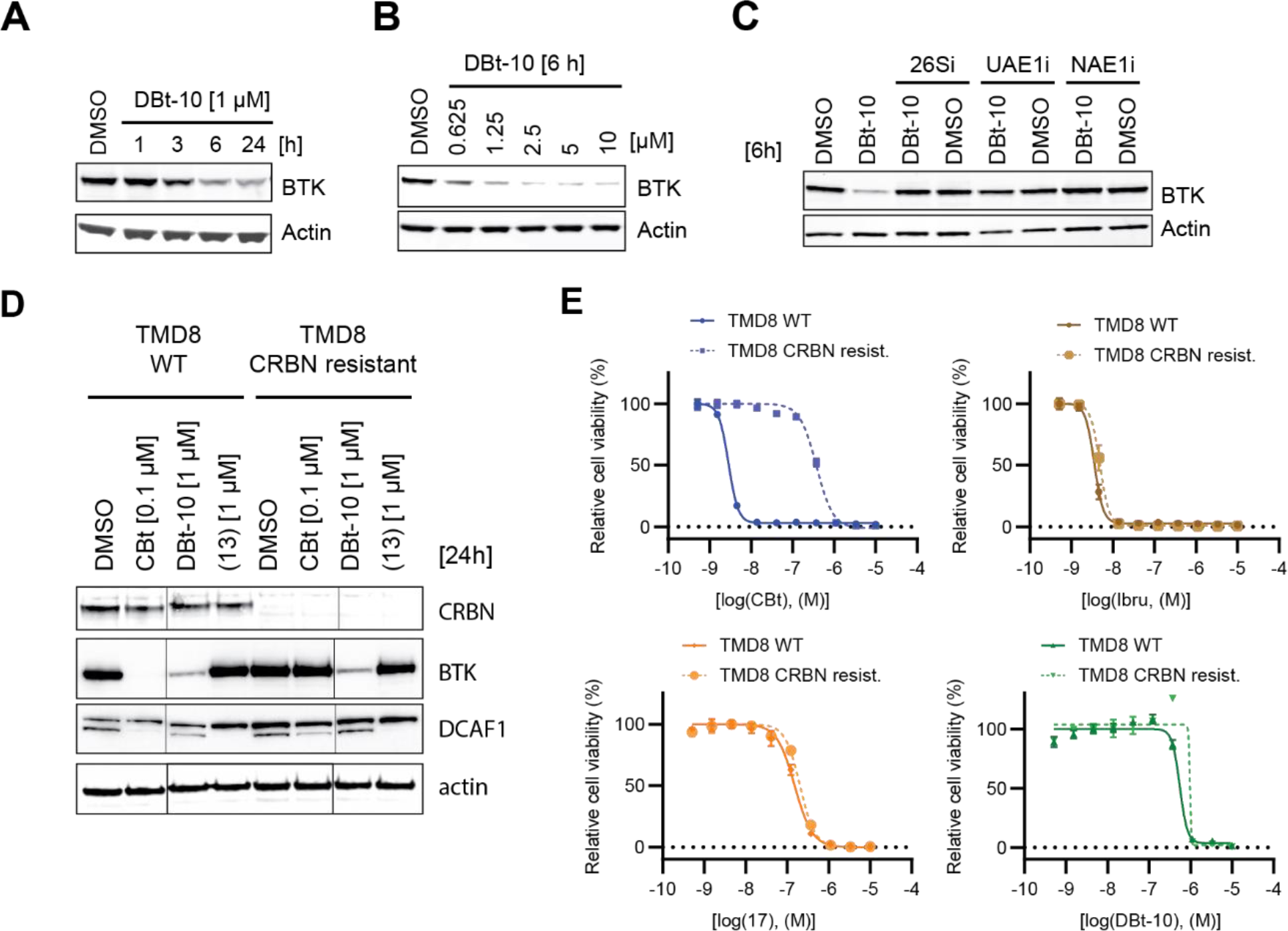
Characterization of a potent DCAF1-BTK degrader and rescue of BTK degradation in CBt resistant cells. (A) Immunoblot analysis of TMD8 cells treated with 1 µM (**DBt-10**) for 1, 3, 6 and 24 hours time points. (B) Immunoblot analysis of TMD8 cells treated with various doses of (**DBt-10**) for 6 hours. (C) Immunoblot analysis of TMD8 cells cells pretreated with proteasome inhibitor Bortezomib (26Si) [1000nM], Ubiquitin E1 inhibitor (UAE1i) [1000 nM] and NEDD8 E1 inhibitor (NAE1i) [1000 nM] for 30 min, followed by (**DBt-10**) treatment [2500 nM] for additional 6h. (D) Immunoblot analysis of TMD8 WT and TMD8 CRBN resistant cells after 0.1µM CBt and 1µM DBt-10 treatment for 24h. (E) Viability measured by celltiter-Glo for CBt, Ibrutinib (Ibru), BTKi (17) and DBt-10 in WT (solid) vs CRBN resist. (dotted) TMD8 cells.

To study **DBt-10** in a CRBN-BTK PROTAC resistant setting, we generated TMD8 cells resistant to CRBN-BTK mediated PROTACs by prolonged treatment with escalating suboptimal doses of a recently patented CRBN-BTK PROTAC (**CBt**) (WO2021/53495 A1). Probing both TMD8 cell settings (**CBt** sensitive *vs* resistant) with **CBt** enabled BTK degradation only in the WT but not in the **CBt** resistant TMD8 cells, in line with loss of CRBN expression in the latter (Supplementary Fig. 5C).

Probing these TMD8 **CBt** sensitive and resistant cells with the DCAF1-BTK PROTAC **DBt-10** enabled equipotent degradation of BTK, consistent with retention of DCAF1 protein levels independent of CRBN status (Fig. 5D). Next, we assessed if degradation or lack thereof translates into viability effects in the TMD8 **CBt** sensitive and resistant cells when incubating with **CBt** or **DBt-10**. As expected, treatment of these two TMD8 cell lines with **CBt** led to a 100-fold difference in proliferation inhibition, as BTK PROTACs bind to BTK via an inhibitor and BTK inhibition is not impaired by loss of CRBN. We confirmed a CRBN mediated resistance mechanism by probing both **CBt** sensitive and resistant TMD8 cells with BTK inhibitors ibrutinib (**Ibru**) or (**17**), and neither inhibitor showed any relative potency change between the two different cell settings, thus proving that BTK catalytic function is still intact and is not the cause of resistance. Intriguingly, treatment with **DBt-10** also did not show any relative potency differences in proliferation between **CBt** sensitive and resistant settings (Fig. 5E, Supplementary figures 5D/E). The DCAF1 binder (**13**) did not show any reduction of BTK as well, as we do observe a 5-fold window between DBt-10 *vs* (**13**) treatment, which makes us believe that the impact on TMD8 cell proliferation by **DBt-10** is mediated through BTK degradation.

## Discussion

In this study we describe the functionalization of a non-covalent DCAF1 WDR binder (Vulpetti et al. submitted) as a PROTAC. We selected DCAF1 as our ligase of choice due to its unique essentiality within the WDR containing E3 ligase receptor family, with the hypothesis that this might reduce the chance of ligase-specific resistance mechanisms through e.g., loss of ligase expression. DCAF1 as an E3 ligase has been targeted previously by covalent PROTACs (Tao et al., 2022), although in this study *de novo* substrate degradation was only possible when DCAF1 was overexpressed from a very strong CMV-promoter. Since the CRL4^DCAF1^ ligase complex has been structurally described to be in a tetrameric state that likely represents an inhibited state as well as a dimeric, active conformation favored through substrate binding (Mohamed et al., 2021), it is likely that DCAF1 overexpression changes this equilibrium between active and auto-inhibited CRL4^DCAF1^ ligase complex. Our study shows that PROTACs with a reversible non-covalent double digit nM binder to the DCAF1 WDR donut-hole can hi-jack endogenous DCAF1 and mediate degradation of target proteins, confirming accessibility of active CRL4^DCAF1^ ligase. Our degradation studies with BRD9 and tyrosine kinase (dasatinib and BTK) targeting PROTACs show that although DCAF1 is predominantly localized in the nucleus, it can degrade cytosolic and even membrane localized tyrosine kinases such as CSK or TEC, respectively. However, based on the potency of our BRD9-PROTAC **DBr-1** compared to the potency of the first tyrosine kinase PROTAC prototypes **DDa-1** and **DBt3-10** we speculate that DCAF1 might preferentially degrade nuclear targets such as e.g., transcription factors. However, further comparative studies are needed to better understand substrate preferences of DCAF1 compared to other E3 ligase receptors such as CRBN or VHL. After establishing that endogenous DCAF1 can degrade a variety of *de novo* substrates at sub-µM potencies, we wanted to expand our studies to BTK-specific degraders, a target that has been extensively studied and characterized in hematological cancers (Alu et al., 2022; Ran et al., 2022; Wen et al., 2021), with the aim to discover potent DCAF1-BTK PROTACs and test if these can overcome acquired resistance to CRBN-BTK PROTACs. One important confounding factor when we established a BTK degradation assay is that our DCAF1 binder (**13**) as well as some of the DCAF1 PROTACs lead to cellular toxicity at concentrations above 5-10 µM. Therefore, we established a viability score in our flow cytometry-based assays to enable confident degradation data reporting (Supplementary Fig. 3). Our systematic discovery approach revealed observations that led to the final potent DCAF1-BTK PROTAC DBt-10 with minimal off-target toxicities. Key observations we made were that individual potencies for DCAF1 and BTK translated into ternary complex formation using linear PEG linkers. However, shorter linkers did not translate into potent ubiquitination activities and only a very long PEG linker (PEG-12) allowed ubiquitination at 10-fold lower concentrations *in vitro*, probably due to enhanced flexibility to more optimally position various surface exposed lysine residues, an observation that translated into better degradation in cells. We note that the 10-fold boost of cellular degradation potency (DC_50_) and net degradation of app. 100% versus 50% of **DBt-5** compared to **DBt-3** does not fully reflect the *in vitro* observation with respect to ubiquitination activity and complex formation. However, we do note a 5-fold higher cellular BTK occupancy for **DBt-5** compared to **DBt-3** indicative of a better cell permeability of the former. We further extended our linkerology studies towards more rigid non-linear spiro-cyclic linkers as well, as we exchanged the BTK binder. Both of these changes were well tolerated in terms of complex formation as measured by SPR, but reduced the cellular potency of BTK binding and in the case of **DBt-10** the cellular permeability. Nevertheless, **DBt-10** proved to potently degrade endogenous BTK with sub-µM potencies and highlighted the successful change of a prototypic PEG linker-based PROTAC to a more complex drug-like molecule.

It is important to note that all the biophysical and biochemical studies were performed with truncated BTK (kinase domain) and DCAF1 (WDR domain) and the cellular experiments were all performed with full-length BTK and endogenous DCAF1 that might confound some of the observations. Further studies are necessary to understand the full potential of DCAF1 as an efficient ligase for PROTAC-mediated degradation, and our limited comparative studies between DCAF1 and CRBN PROTACs (dasatinib: **DDa-1** vs **CDa-1**, Fig. 3B; BTK: **DBt-10** vs **CBt**, Fig. 5D) revealed that in both cases CRBN-mediated degradation was more efficient, although this is beyond the scope of this work.

Our in-depth DCAF1-BTK degrader characterization further supports many previous observations in the TPD field that efficient PROTAC discovery cannot currently be predicted through simple rules and single parameters. Successful PROTAC discovery is a multi-parametric empiric optimization process, which in our case finally led to the discovery of the DCAF1-BTK PROTAC **DBt-10** with sufficient potency to enable studies in a CRBN-BTK PROTAC resistant setting. Indeed, with this optimized PROTAC **DBt-10** we could show that BTK is still degraded with comparable efficiency in CBRN-BTK PROTAC resistant cells that have lost CRBN expression, and this translates to equipotent inhibition of cell proliferation independent of CRBN status in these BTK dependent DLBCL lymphoma cell lines. These observations demonstrate that targeting an alternative ligase in CRBN-PROTAC resistant settings is a viable option to overcome resistance and might provide a strategy for combinations of degraders which engage different ligases.

## Acknowledgements

We thank our extended team members, the TPD initiative and in particular Marcel Re ck, Alexandre Luneau, Emine Sager, Corinne Marx, B. Forrester, L. Tordella and C. Wiesmann for support and scientific input and the NIBR Innovation Postdoctoral program for support of M. S. We wanted to thank Prof. S. Tohda for providing TMD8 cells.

## Author Contributions

Conceptualization: MS, MR, CRT, TR, PH, AV, NHT Writing – original draft: MS, MR, CRT

Writing – original, review and editing: TR, MR, MS, GH, CRT

Experimental contribution: MS, MR, XL, FM, SF, FG, XLi, M-TS, JT, DB, PL, RA-R, SC, BP, AH, MSci, DG, KC, BB-P, MM, MN, RM, MH, JA

Chemistry: TZ, NS, BYC, RM, PI

Data & computational analysis: FS, SG, EA

## METHODS

**KEY RESOURCES TABLE**

| REAGENT or RESOURCE | SOURCE | IDENTIFIER |
| --- | --- | --- |
| **Antibodies** |  |  |
| IRDye 680RD Goat anti-Mouse secondary antibody | LI-COR | Cat # 926-68070 |
| IRDye 800CW Goat anti-Rabbit secondary antibody | LI-COR | Cat # 926-32211 |
| HRP Goat anti-rabbit secondary antibody | Abcam | Cat # 7090 |
| HRP Sheep anti-mouse secondary antibody | GE Healthcare | Cat # NA931V |
| Rabbit polyclonal anti-BRD9 | Bethyl Laboratories | Cat # A303-781A |
| Mouse monoclonal anti-actin | Cell Signaling Technology | Cat # 3700 |
| Rabbit polyclonal anti-actin | Cell Signaling Technology | Cat # 4970 |
| Mouse monoclonal anti-CAS9 | Cell Signaling Technology | Cat # 14697 |
| Rabbit monoclonal anti-LYN | Cell Signaling Technology | Cat # 2796 |
| Rabbit monoclonal anti-LIMK2 | Cell Signaling Technology | Cat# 3845 |
| Rabbit polyclonal anti-c-ABL | Cell Signaling Technology | Cat # 2862 |
| Rabbit monoclonal anti-CSK | Cell Signaling Technology | Cat # 4980 |
| Mouse monoclonal anti-α-tubulin | Cell Signaling Technology | Cat # 3873 |
| Rabbit polyclonal anti-BTK | Cell Signaling Technology | Cat # 8547 |
| Rabbit polyclonal anti-CYLD | Cell Signaling Technology | Cat # 4495 |
| Rabbit monoclonal anti-α-tubulin | Cell Signaling Technology | Cat # 2125 |
| Rabbit polyclonal anti-DCAF1/VprBP | Novus | Cat # NBP1-05953 |
| **Bacterial and Virus Strains** |  |  |
| **Biological Samples** |  |  |
| **Chemicals, Peptides, and Recombinant Proteins** |  |  |
| Protease Inhibitor Tablet | Thermo Fisher Scientific | Cat # A32955 |
| Phosphatase Inhibitor Cocktail | Thermo Fisher Scientific | Cat # A32957 |
| UAE1i (TAK243) | Selleckchem | Cat # S8341 |
| MLN4924 (NAEi) | R&D Systems Inc. | Cat # I-502-01M |
| Dasatinib | Selleckchem | Cat # S1021 |
| ViraPower™ Lentiviral Packaging Mix | Thermo Fisher Scientific | Cat # K497500 |
| Bortezomib | Sigma Aldrich | Cat # 5043140001 |
| NHS-activated Sepharose™ 4 Fast Flow | Cytiva | Cat# 17090601 |
| 11-Azido-3,6,9-trioxaundecan-1-amine | Millipore Sigma | Cat # 17758 |
| 2-(2-aminoethoxy)ethanol | Millipore Sigma | Cat # A54059 |
| Trypsin protease, MS grade | Thermo Fisher Scientific | Cat # 90058 |
| Glu-C endoproteinase | Thermo Fisher Scientific | Cat # 90054 |
| 16plex TMTpro | Thermo Fisher Scientific | Cat # A44520 |
| Avi-His-PreSc-Btk(389-659) | In-house |  |
| Ubiquitin activating enzyme E1 (human), (recombinant) (Histag) | Enzo | Cat # BML-UW9410 |
| In-house NEDD8~CRL4DCAF1 Complex (E3) | In-house |  |
| Ubch5b (human), (recombinant) (His-tag) | Enzo | Cat # BML-UW9060 |
| Ubiquitin (human), (fluorescein labeled) | Enzo | Cat # BML-UW1240 |
| LanthaScreen® Tb-Streptavidin | Thermo Fisher Scientific | Cat # PV3965 |
| Annexin V-Pacific Blue | Thermo Fisher Scientific | Cat # R37177 |
| Live/Dead Far Red | Invitrogen | Cat # L34973 |
| **Critical Commercial Assays** |  |  |
| DC Protein Assay Reagent A | Bio-Rad | Cat # 5000113 |
| DC Protein Assay Reagent B | Bio-Rad | Cat # 5000114 |
| Cell Titer Glo | PROMEGA | G7573 |
| HiBIT | PROMEGA |  |
| FireFly | PROMEGA |  |
| **Deposited Data** |  |  |
| DCAF1-WD40 domain-Cpd. 13 X-ray structure | PDB | 8OO5 |
| DCAF1-WD40 domain-Cpd. 15 X-ray structure | PDB | 8OOD |
| **Experimental Models: Cell Lines** |  |  |
| HEK293T |  |  |
| HEK293 BRD9-HiBIT pool | PROMEGA | CS3023119 |
| HEK293 BRD9-HiBIT FF/CAS9 | this study |  |
| 293FT | Invitrogen | R70007 |
| TMD8 | PMID: 16780947 | Prof. Tohda |
| TMD8 BTK-GFP/mCh | this study |  |
| Experimental Models: Organisms/Strains |  |  |
| Oligonucleotides |  |  |
| sgRNAs from STAR methods |  |  |
| Recombinant DNA |  |  |
| Plasmid: UBC_hNOS2[NM_000625.4]/3xGS/HA | Vector Builder | N/A |
| Gateway™ pENTR™ 1A Dual Selection Vector | Invitrogen | Cat # A10462 |
| pLenti4/V5-DEST™ Gateway™ Vector | Invitrogen | Cat # V49810 |
| Gateway™ pDONR™ 221 Vector | Invitrogen | Cat # 12536017 |
| Software and Algorithms |  |  |
| PRISM | GraphPad | N/A |
| Adobe Illustrator | Adobe Creative Cloud | N/A |
| Fiji | ImageJ | N/A |
| Proteome Discoverer 2.4 | Thermo Scientific | N/A |
| Spotfire | Tibco | N/A |
| Python | Anaconda | v 3.6.10 |
| Numpy | Anaconda | v 1.17.2 |
| Other |  |  |

### LEAD CONTACT AND MATERIALS AVAILABILITY

Requests for further information, resources, and/or reagents should be directed to, and will be fulfilled by the Lead Contact, Martin Schroeder.

### EXPERIMENTAL MODEL AND SUBJECT DETAILS

Cells used: HEK293T cells (donor sex: female); BRD9-iBIT KI pool (PROMEGA #CS3023119 BRD9-HiBiT KI HEK293 cells (donor sex: female), were cultured in DMEM (Gibco, Thermo Fisher Scientific) supplemented with 10% fetal bovine serum (FBS) and 1% penicillin/streptomycin (Thermo Fisher Scientific). TMD8(Tohda et al., 2006) (donor sex: male), TMD8 BTK-GFP/mCh wer cultured in α-MEM with nucleosides (AMIMED I-23F01-I) supplemented with 10% fetal bovine serum (FBS), 1% L-Gln (25030-024;Invitrogen) and penicillin/streptomycin (Thermo Fisher Scientific).

## METHOD DETAILS

### Extraction of genetic dependencies of WDR E3 ligases

We manually annotated all E3 ligase related proteins and function from a published resource of 361 WDR containing proteins(Schapira et al., 2017) and further manually classified the E3 subfamilies (Supplementary Table 1). From this table we extracted Achilles Demeter2 scores from the DepMap 22Q2 release (D2_combined_gene_dep_scores.csv at https://depmap.org/portal/download/all/). Downloads, data handling, and plotting were performed using Python 3.9.5.

### Protein Production

His6-TEV-DCAF1(1039-1401)Q1250L, His-TEV-DCAF1(1079-1393)Q1250L, Avi_DCAF1(1073-1399)_E1398S_His, Strep-PreSc-DCAF1(987-1507), DDB1(1-1140), His-TEV-CUL4(38-759), His-TEV-Rbx1(12-108), His_PreSc_BTK (389-659), His_PreSc_Avi_BTK (389-659) were all subcloned into pIEx/Bac-3 derived vectors for expression in Sf9 cells using the flashBAC ™ system. Recombinant baculoviruses were generated and amplified followed by large scale expression in Sf9 cells as previously described(Rieffel et al., 2014). Transfected cells were typically grown at 27℃ for up to 62h before harvesting by centrifugation. For in-vivo biotinylation BirA was co-transfected.

Cell pellets were resuspended in lysis buffer (see Method table 1) supplemented with o’complete (EDTA free) protease inhibitor cocktail (Roche) and Benzonase and lysed by sonication or use of a microfluidizer. After removal of cell debris by centrifugation at 4℃ at 50,000xg for at least 30 minutes the cleared supernatants were purified using Ni-NTA columns (Cytivia) on AKTA™ systems (Cytivia). For proteins containing cleavable affinity tags, C3 protease (PreScission™) or TEV protease were used to cleave the tag. Proteins were further purified by Ion exchange chromatography (see Methode table 1) followed by a final Size Exclusion Chromatography step on SuperdexS200 or SuperdexS75 columns (Cytivita). Proteins were concentrated, aliquoted and flash-frozen in liquid nitrogen.

The DCAF1//DDB1//CUL4A//RBX1 complex was expressed, purified and neddylated as previously described(Mohamed et al., 2021). The enzymes PPBP1-UBA3 and UBC12 as well as the substrate NEDD8 were expressed and purified as reported before(Duda et al., 2008; Huang et al., 2007; Huang et al., 2008).

BRD9 (130-250) was subcloned into a pGEX6P1 vector and transformed into E.Coli Rosetta cells. Cells were grown at 37℃ to an OD600 of 0.7-1.0 in Terrific Broth (Teknova cat# G8110). Expression was induced with 0.2mM at 20℃ overnight. Cells were harvested by centrifugation. The cell pellet was resuspended in lysis buffer (see Method table 1) and cells were lysed by passing through a microfluidizer. Cell debris was removed by centrifugation at 4℃. Affinity chromatography was performed on Glutathione beads (GE Bioscience cat# 17-0756-05) and successively the affinity tag was cleaved by addition of C3 protease (PreScission™) while dialyzing against 1xPBS pH7.4, 400mM KCl, 5mM DTT. After a reverse affinity step with fresh GST beads the protein was applied to a Superdex S200 column equilibrated in the final buffer (see Method table 1). The protein was concentrated, aliquoted and flash-frozen in liquid nitrogen

| Protein | Expression system | Lysis buffer | Ion-exchange column | Final buffer |
| --- | --- | --- | --- | --- |
| His6-TEV-DCAF1(1039-1401)Q1250L | Sf9 insect cells | 50mM Tris pH8.0, 200mM NaCl, 0.1% Triton X-100, 4mM TCEP | ResourceQ | 20mM Hepes pH 7.5, 200 mM NaCl, 2mM TCEP |
| His-TEV-DCAF1(1079-1393)Q1250L | Sf9 insect cells | 50mM Tris pH8.0, 200mM NaCl, 0.1% Triton X-100, 4mM TCEP | ResourceQ | 20mM Hepes pH 7.5, 200 mM NaCl, 2mM TCEP |
| Avi_DCAF1(1073-1399)E1398S-His6 | Sf9 insect cells<br>Coexpressed with BirA | 50 mM Tris (pH 8.0), 300 mM NaCl, 10% Glycerol (w/v), 5 mM TCEP, 5 mM Imidazole | Q HP | 50 mM Tris, 300 mM NaCl, 10% Glycerol, 2 mM TCEP (pH 8.0) |
| DCAF1(1073-1399)E1398S-Avi-His6 | Sf9 insect cells<br>Coexpressed with BirA | 50 mM Tris (pH 8.0), 300 mM NaCl, 10% Glycerol (w/v), 5 mM TCEP, 5 mM Imidazole | Q HP | 50 mM Tris, 300 mM NaCl, 10% Glycerol, 2 mM TCEP (pH 8.0) |
| Strep-PreSc-DCAF1(987-1507)//DDB1(1-1140)//His-TEV-CUL4(38-759)//His-TEV-Rbx1(12-108) | Sf9 insect cells | 50mM Tris pH8.0, 200mM NaCl, 4mM TCEP | ResourceQ | 50 mM HEPES pH 7.4, 200 mM NaCl, and 0.25 mM TCEP |
| His_PreSc_BTK (389-659) | Sf9 insect cells | 50mM Tris pH7.6, 400mM NaCl, 10% Glycerol, 3mM TCEP | ResourceQ | 20mM TrisHCl, 100mM NaCl, 3mM TCEP, 1mM EDTA, pH8.0 |
| His_PreSc_Avi_Btk (389-659) | Sf9 insect cells<br>Coexpressed with BirA | 50mM Tris pH7.6, 400mM NaCl, 10% Glycerol, 3mM TCEP | ResourceQ | 50mM Tris pH8.0, 200mM NaCl, 5% Glycerol, 2mM TCEP |
| GST-PreSc-BRD9 (130-250) | E.Coli | 1x PBS pH7.4, 400mM KCl, 5mM DTT, 1mM MgCL<sub>2</sub>, 10% Glycerol, 1mM PMSF | n/a | 25mM HEPES pH7.5, 100mM KCl, 5mM DTT |

Method table 1: Purification details of recombinant proteins used.

### Surface Plasmon Resonance (SPR)

Surface plasmon resonance (SPR) experiments were performed on Biacore T200 and 8 K+ instruments (Cytiva). Biotinylated Avi_DCAF1(1073-1399)E1398S-His6 was immobilized (approx. 3000-4000 RUs) on a SA chip. Compounds were diluted in the SPR running buffer (20 mM HEPES, pH 8, 200 mM NaCl, 0.5 mM TCEP, 0.01% Tween-20, 5% Glycerol) for binary compound-DCAF1 complex assessments (typically in a range from 0.003 µM up to 100 µM). For the measurement of the VHF543-DCAF1 complex on the Biacore T200 samples were applied to the flow cell at a flow rate of 30µL/minute. The association was assessed for 60s and the dissociation was monitored for 410s. For measurements of all other compounds and ternary complexes on the Biacore 8 K+ the association was assessed for 150s and the dissociation was monitored for 1510s. For the ternary complexes the compounds were diluted in the SPR running buffer supplemented with 0.2µM BTK (389-659) or BRD9 (130-250). Binary complexes were analyzed with a kinetic 1:1 fit model to obtain Kd values. The fit of apparent steady state RU intensities of the ternary complexes was performed using a bell-shaped curve algorithm implemented in Graphpad Prism. For the off-rate determination of the ternary complexes only the datapoints of compound concentrations contributing to the transition at lower compound concentration were included in a 1:1 dissociation fit model. Residence times (t1/2) were calculated by the inversion of the off-rate.

### X-Ray Crystallography

The DCAF1 apo crystals were grown by mixing cleaved His-TEV-DCAF1(1079-1393)Q1250L at 9mg/mL with 1.91M LiSO4 and 0.1M Tris pH7.5 in a 1:1 ratio (200nL + 200nL) in sitting drop crystallization plates at 20℃. Crystals were soaked with 1mM (**13**) for 15 minutes and cryo-protected by the addition of 20% ethylene glycol and frozen in liquid N2.

The complexes DCAF1-GQN626 and DCAF1-IVH258 were obtained by incubation of cleaved His6-TEV-DCAF1(1039-1401)Q1250L at 1.6mg/mL with a 1:200 dilution of Trypsin Gold (Promega) for 2h on ice. The cleaved protein was further purified on HiLoad 16/60 Superdex 75 equilibrated with 50mM Hepes, 200mM NaCl, 0.5mM TCEP. Fractions containing a fragment of ∼27kDa were pooled and concentrated to 13mg/mL. Protein was then mixed with 2mM compound and incubated briefly on ice before mixing with reservoir solution 1:1 (100nL + 100nL) in sitting drop crystallization plates at 20℃. Crystals were obtained within 1-3 days with reservoir solutions containing 2.13M LiSO4 and 0.1M Tris pH7.5 and were cryo-protected by quickly transferring the crystals to reservoir solution containing additional 20%(V/V) ethylene glycol.

Datasets were collected at the PXII beamline at the Swiss Lightsource (SLS). Data was integrated by XDS (Kabsch, 2010) and successively merged and scaled by AIMLESS(Evans & Murshudov, 2013) in the CCP4I2 interface (Winn et al., 2011). Molecular replacement was performed using Phaser(McCoy et al., 2007) with the starting coordinates 4PXW. Refinement was performed in iterative cycles of modelbuilding in Coot (Emsley & Cowtan, 2004) and refinement in Refmac5 (Murshudov et al., 1997). Geometrical correctness of the model was validated by Molprobity (Williams et al., 2018). Coordinates were deposited in the PDB with accession codes: 8OO5 and 8OOD and data processing and refinement statistics can be found in Supplementary table 8.

#### Chemical Proteomics pull-down

Synthesis of azide-functionalized resin. Azide-modified sepharose fastflow4 resin was created by adding 2 umol of 11-azido-3,6,9-trioxaundecan-1-amine/mL bead volume and 30 uL triethylamine in anhydrous DMSO. This coupling reaction was allowed to proceed overnight at room temperature with end-over-end agitation. Next, 100 uL 2-(2-aminoethoxy)ethanol was added directly to the coupling reaction and allowed to procced overnight at room temperature with end-over-end agitation. The resulting azide-functionalized resin was then washed several times with anhydrous DMSO.

Synthesis of DCAF1 affinity matrix. 10 umol YYJ150 was added to 1 mL of the azide-functionalized resin in the presence of 1 mM CuSO4, 1 mM TBTA, and 50 mM TCEP in anhydrous DMSO. The YYJ150 coupling reaction was allowed proceed overnight at room temperature with end-over-end agitation, then washed extensively with anhydrous DMSO.

Lysate generation. HEK293T cell pellets were suspended in lysis buffer (50 mM HEPES pH 7.5, 150 mM NaCl, 1.5 mM MgCl2, 1 mM DTT, 0.8% NP-40, 5% glycerol, and 1× protease inhibitors (Thermo)) at 2× pellet volume. The samples were sonicated on ice for 30 seconds at 20% amplitude with a 2 second pulse ON and 3 second pulse OFF. Cell debris was removed by centrifugation at 20,000 × g for 20 minutes at 4°C. Protein concentration was determined by BCA (Pierce) and diluted down to 5 mg/mL with ChemProt buffer (50 mM HEPES pH 7.5, 150 mM NaCl, 1.5 mM MgCl2, 1 mM DTT, 0.4% NP-40, 1× protease inhibitors (Thermo).

Chemical proteomic experiments. 1 mL of 5 mg/mL HEK293T lysates was treated with 100 uM YYJ150 DMSO for 1 h at 4°C. Following incubation, the lysates were incubated with 35 μL YYJ150-functionalized resin 4°C for 4. Unbound proteins on affinity matrix were washed and bound proteins were eluted with 2× LDS sample buffer (NuPAGE, Invitrogen) containing 10 mM DTT, alkylated with 25 mM iodoacetamide, and processed through detergent removal columns (Thermo). In-solution digestion and isobaric labeling with TMT reagents was performed according to standard procedures. Quantitative proteomics by two-dimensional nano liquid chromatography-tandem mass spectrometry (nLC-MS/MS). Quantitative proteomics was performed as previously described.2 The TMTlabeled sample was reconstituted in 0.1% formic acid (FA) / 2% acetonitrile for offline high-pH reversed phase separation (RP10) using a Dionex UltiMate 3000 high-performance liquid chromatography (HPLC) system with fraction collection using Chromeleon v.6.8 (Thermo) software. Separation was achieved on an Xbridge C18 3.5 μm 2.1 × 150 mm HPLC column (Waters) using a flow rate of 250 μL min-1 and the following ternary gradient and conditions: mobile phase A = water (HPLC grade), mobile phase B = acetonitrile (HPLC grade), and mobile phase C = 200 mM ammonium acetate, pH 10. Mobile phase C is held at 10% throughout gradient. Starting conditions are 89% A and 1% mobile phase B, ramping mobile phase B to 80% over 65 min. Fractions were collected in 96-well plate at 100 sec per well. Fractions were reduced down to 12 fractions, dried, and reconstituted in 0.1% FA / 2% acetonitrile for nLC-MS/MS. All data were acquired using a Orbitrap Fusion Lumos coupled to an Easy-nLC 1200 nanoflow liquid chromatograph operating at 300 nL min-1. Peptides were cleaned up using a 1 cm online custom trap then eluted on a custom 20 cm laser pulled 75 um column with C18 resin. Peptides were quantified using the synchronous precursor selection (SPS-MS3) or higher-energy C-trap dissociation (HCD) method for TMT quantitation with 120k MS1 resolving power, 50 ms max injection time and 100% automatic gain control (AGC). MS2 spectra were selected using the top ten most abundant features with a charge state between 2-6 using collision induced dissociation (CID) in the ion trap. The AGC was set to 100% with an isolation width of 0.7. MS3 was performed on the top 10 most abundant MS2 features between 400-1600 amu with a collision energy of 55%, AGC of 250% and 50k resolution. For HCD TMT quantitation the MS2 isolation width was set to 1.2 with a collision energy of 55%, AGC of 250% and a resolution of 50K. Mass spectrometry data processing. Peptide and protein identification and quantification were performed using Proteome Discoverer version 2.2 (Thermo). MS data were searched using the Mascot (Matrix Science) search engine against the UniProt reference database (human proteins, 42,233 entries, downloaded 2017) containing common contaminants and reversed sequences. Carbamidomethylated cysteine, oxidation of methionine, and TMT modification on N-termini and lysine were set as dynamic modifications. Trypsin was specified as the proteolytic enzyme with up to one missed cleavage site allowed. Precursor and fragment ion tolerances were set to 10 ppm and 0.8 Da, respectively. Search results were filtered for a minimum of 2 unique peptides, 1% FDR peptide and protein identification and quantification, and common contaminants. Protein abundances were normalized and competition ratios were calculated using scripts provided by Proteome Discoverer. Dose-response curves were fitted to a three-parameter log logistic regression using GraphPad Prism.

#### Immunoblotting

Whole cell lysates for immunoblotting were prepared by pelleting cells at 4°C (300 g) for 3 minutes. Cell pellets were then washed 1× with PBS and resuspended in RIPA lysis buffer (VWR, cat # 97063-270) supplemented with protease (Thermo Scientific, cat # A32955) and phosphatase (Thermo Scientific, cat # A32957) inhibitor tablets. Lysates were clarified at 13,200 rpm for 15 min at 4°C prior to quantification by Lowry assay (Bio-Rad cat # 5000113 and cat # 5000114). Whole cell lysates were loaded into 4-20% Criterion TGX Precast 18 well gels (Bio-Rad, cat # 5671094) and separated by electrophoreses at 120 V for 1 hr. The gels were transferred to a nitrocellulose membrane using the Trans-Blot Turbo (Bio-Rad) for 7 minutes and then blocked for 1 hr at room temperature in Odyssey blocking buffer (LICOR Biosciences, cat # 927-50000). Membranes were probed with the appropriate primary antibodies (diluted 1:1000) overnight at 4°C in 20% Odyssey blocking buffer in 1× TBST. Membranes were washed three times with 1× TBST (5 minutes per wash), and then incubated with IRDye goat anti-mouse (LICOR, cat # 926-32210) or goat anti-rabbit (LICOR, cat # 926-32211) secondary antibody diluted 1:10,000 in 20% Odyssey blocking buffer for 1 hr at room temperature. After three 5-minute washes with 1× TBST, the immunoblots were visualized using the ODYSSEY Infrared Imaging System (LICOR). Alternatively, SDS-PAGE resolved proteins were transferred to a PVDF membrane using wet blotting system (Invitrogen). Membranes were blocked in 5% Milk-PBS-Tween 20 for 1 hour and incubated with primary antibody over night at 4°C. Secondary antibodies (secondary antibody: anti-rabbit, goat pAb to Rb-IgG (Abcam, #7090); secondary antibody: anti-mouse, sheep IgG HRP-linked (GE healthcare, #NA931V)).were incubated for 1 hour at room temperature (RT) before blots were developed on film using the Dection kit SuperSignal West Femto Maximum Sensitivity Substrate (Thermo Fisher, #34096).

### Engineering of BRD9-HiBiT/FF/CAS9 cells

We performed clonal selection from purchased 293 BRD9-HiBiT pools (PROMEGA # CS3023119) using limiting dilution and screened for HiBIT expression using Nano-Glo HiBiT Lytic Detection System (Promega # N3030). We further engineered a validated Hibit clone by lentiviral delivery of both Cas9 in pNGx-LV-c004 as described previously (PMID: 27351204), and FireFly Luciferase (in pNGx-LV-u003, constructed in-house, using an EF1alpha promoter, C-terminal PEST domain fusion, and a hygromycin resistance cassette). To produce lentiviral particles, each plasmid was co-transfected with ViraPower™ Lentiviral Packaging Mix into 293T cells. Supernatant from both plasmid transfections containing viral particles were spun down at 2000rpm for 10 minutes, aliquoted and frozen in at -80 °C. Thawed virus supernatant was used to co-infect 293 BRD9-HiBIT cells. And selected for Cas9 with 10 µg/ml Blasticidine, and for Firefly luciferase (FF) with 300 µg/ml Hygromycin for two weeks to obtain BRD9-HiBIT/CAS9/FF expressing cell lines. We performed clonal selection from this BRD9-HiBIT/CAS9/FF pool by limiting dilution. Multiple recovering clones were tested for HiBIT and FF signal using Nano-Glo HiBiT Dual-Luciferase Reporter System (Promega # CS1956A08) as well as CAS9 expression by western blotting using CAS9 antibody (Cell Signalling Technology cat# 14697). All subsequent experiments described in this publication were performed with this isolated clone referred to as BRD9-HiBiT/CAS9/FF cells.

### BRD9-HiBIT protein abundance luminescence assay

To measure BRD9-HiBiT abundance in BRD9-HiBiT/FF/CAS9 we pre-treated the cells with corresponding Ubiquitination pathway inhibitors (NAE1i, 26Si or UAE1i) for 2h before DBr-1 PROTAC treatment at various doses for 2h. HiBIT and FF signals were measured using using Nano-Glo HiBiT Dual-Luciferase Reporter System (Promega # CS1956A08) according to the manufacturer’s manual with a EnVision plate reader (Perkin Elmer).

### sgRNA-mediated control knockout studies

Individually designed sgRNAs targeting BRD9 (a mix of 3 guides: CGACTTTGATCCTGGGAAGA, GCCTCTAAAGCTAGTCCTGA, GTCACGGATGCAATTGCTCC), DCAF1 (mix A: TCACACATCTTGAACCTTCC and GAAGGAACCCTTACCCTGGA; mix B: CACGCGACACTGATCCAGAG and TCGCGTGGTGTTCAATCACA; mix C: GATGTTCCGTTTCACATCAA and AGTCTTTGGTGTCTGTACAC), PSMA1 (mix of 4 guides: ACAAGACGAGACACAGGCAG, ATGACAATGATGTCACTGTT, CCGCAATTGAGATACCAATA, and TTTTATGCGTCAGGAGTGTT), CUL4A (individual guides, guide A: CATAGGTGCGGTCCAAGAAC, guide B: CGCAGTTGCTTGTAGAGCAT, guide C: CTCCTCGAGGTTGTACCTGA, and guide D: TCTCGCGCTCGATCAGCAGT), CUL4B (individual guides, guide A: CATCAAACCCTACAAACTCC, guide B: GCCGAATCCCTGGGTTGTAA, guide C: GCTTCTTCTGTATCGGTACG, guide D: GTTTGATGCGAAGATGGCTG), DDB1 (individual guides, guide A: CAATAATGCCGGTCTCTGAG, guide B: GACATCATTACGCGAGCCCA, guide C: GATCTATGTGGTCACCGCCG, guide D: GTCGTACAACTACGTGGTAA), RBX1(individual guides, guide A: ATGGATGTGGATACCCCGAG, guide B: GAAGAGTGTACTGTCGCATG, guide C: GTCCATTGGACAACAGAGAG, guide D: TCGCTGGCTCAAAACACGAC) and sgAAV1 as ctrl (GGGGCCACTAGGGACAGGAT) were cloned into pNGx-LV-g003 lentiviral plasmid as previously described (DeJesus et al., 2016) for lentiviral infection of BRD9-HiBiT/CAS9/FF cells. Lentiviral particles were produced as described above and cells were infected either with sgRNA-expressing particles against an individual target gene or co-infected with a combination of two particles against *CUL4A* and *CUL4B* (volume ratio 1:1) and selected 24 h post infection for 4 days with 1 µg/ml Puromycin. To determine genetic rescue from BRD9 PROTAC mediated degradation we performed a BRD9-HiBIT protein abundance luminescence assay as described.

### Arrayed CRISPR screen with a ubiquitin pathway sgRNA sublibrary

The arrayed ubiquitin pathway rescue screen is outlined in Supplementary Fig. 2A. On day 1 we seeded 500 BRD9-HiBiT FF/CAS9 293 cells per well of a 384 well plates (Corning Cat # 3570) and stamped on day 2 each library plate in triplicate assay plates. Per assay plate we used 1 µl of ready-to-use lentiviral particles of the ThermoFisher Ubiquitin CRISPR LentiArray Library (A42270) targeting 943 genes with 3722 sgRNAs, that are pooled by 4 guides per gene in average. On day 3, infected cells were selected with 1 µg/ml Puromycin. On day 7, after 4 consecutive days of Puromycin selection, we incubated cells for 2h with BRD9 PROTAC DBr-1 before performing BRD9-HiBIT and FireFly luminescence assay using the Nano-Glo HiBit dual-Luciferase Reporter System (Promega # CS1956A08) according to the manufacturer’s manual.

### Analysis of arrayed CRISPR screen

For hit calling we set a threshold based on a minimal Firefly signal > log10(1.25), this threshold was based on signal measurements of media containing as well as non-infected, puromycin selected control wells. The Firefly signal served as control for well-based variation of cell number and was used to normalize the BRD9-HiBit signals on a plate basis using a non-linear correlation (Supplementary Fig. 2D and E) from this normalized data we calculated a robust z-score per gene (Supplementary Figure F) to feed into a gene-level RSA statistical model (Zeng et al., 2019) to calculate gene effect scores as mx. rel. change and significance (Figure 2H).

#### Proteomics profiling of DCAF1 PROTACs

293T were cultured according to protocol and 24h before compound incubation, 3×106 293T cells were distributed in T25 flasks in 5ml and subsequently incubated with indicated compounds at corresponding concentrations for 6h. Cells were then harvested and washed 2× with PBS before removing all liquid and snap freezing in liquid nitrogen before sample preparation for proteomics.

TMT-labeled peptides were generated with the iST-NHS kit (PreOmics, #P.O.00030) and TMT18plex reagent (Thermo Fisher Scientific, #A52047). Equal amounts of labeled peptides were pooled and separated on a high pH fractionation system with a water-acetonitrile gradient containing 20 mM ammonium formate, pH 10(Wang et al., 2011). Alternating rows of the resulting 72 fractions were pooled into 24 samples, dried and resuspended in water containing 0.1% formic acid.

The LC-MS analysis was carried out on an EASY-nLC 1200 system coupled to an Orbitrap Fusion Lumos Tribrid mass spectrometer (Thermo Fisher Scientific). Peptides were separated over 180 min with a water-acetonitrile gradient containing 0.1% formic acid on a 25 cm long Aurora Series UHPLC column (Ion Opticks, #AUR2-25075C18A) with 75 µm inner diameter. MS1 spectra were acquired at 120k resolution in the Orbitrap, MS2 spectra were acquired after CID activation in the ion trap and MS3 spectra were acquired after HCD activation with a synchronous precursor selection approach(McAlister et al., 2014) using 8 notches and 50k resolution in the Orbitrap.

LC-MS raw files were analyzed with Proteome Discoverer 2.4 (Thermo Fisher Scientific). Briefly, spectra were searched with Sequest HT against the Homo sapiens UniProt protein database and common contaminants (Sep 2019, 21494 entries). The database search criteria included: 10 ppm precursor mass tolerance, 0.6 Da fragment mass tolerance, maximum three missed cleavage sites, dynamic modification of 15.995 Da for methionines, static modifications of 113.084 Da for cysteines and 304.207 Da for peptide N-termini and lysines. The Percolator algorithm was applied to the Sequest HT results. The peptide false discovery rate was set to 1% and the protein false discovery rate was set to 1%. TMT reporter ions of the MS3 spectra were integrated with a 20 ppm tolerance and the reporter ion intensities were used for quantification. All LC-MS raw files were deposited to the ProteomeXchange Consortium(Vizcaíno et al., 2014) via the PRIDE partner repository with the dataset identifier PXD00000. Protein relative quantification was performed using an in-house developed R (v.3.6) script. This analysis included multiple steps; global data normalization by equalizing the total reporter ion intensities across all channels, summation of reporter ion intensities per protein and channel, calculation of protein abundance log2 fold changes (L2FC) and testing for differential abundance using moderated t-statistics(Ritchie et al., 2015) where the resulting false rate discovery (FDR) p values (or q values) reflect the probability of detecting a given L2FC across sample conditions by chance alone.

### Engineering of stably expressing TMD8 BTK-GFP-chysel-mCHERRY sensor cells and flow cytometry to monitor BTK-GFP degradation

The bicistronic pLenti6 BTK-GFP-chysel-mCHERRY expression vector was cloned as described (Schukur et al., 2020) using Gateway cloning with a pENTR-BTK plasmid lacking a STOP codon. TMD8 cells were then infected with lentiviral particles from this vector as described above and stably transfected cells were selected with blasticidin at a concentration of 10μg/mL referred to as TMD8 BTK-GFP/mCh cells.

For flow cytometry measurements, 25’000 exponentially growing TMD8 BTK-GFP/mCh cells were plated in 100uL of each well of a 96-well plate and grown for 24 hours. Cells were then dosed with corresponding compounds and doses from 10mM DMSO stock for 24 hours, after which GFP and mCherry fluorescence was read with a Beckmann Coulter Cytoflex. Cells were gated based on Forward & Side Scatter and selection for Single cells using SSC-A vs. SSC-H. Further, the population of viable cells was defined by the forward & side-scatter characteristics of DMSO-treated cells. To confirm cells outside of this population as non-viable, cells were plated and treated for 24 hours, washed with PBS and stained for 30 mins with two markers of early (Annexin-V) and late (LIVE/DEAD) apoptosis. Effects on viability was measured by the reduction of the percentage of cells included in this viable-population gate with increasing doses of compound tested. Degradation of BTK was calculated based on subtracting background (BG) fluorescence of parental TMD8 signal via (GFP-BG)/(mCherry-BG) of viable cells. Median ratios of 3 technical (different wells) replicates were used to calculate daily averages. Each compound dose-response was generated via 3 biological replicates (different days of experiment). Degradation curve parameters (DC_50_ & Abs. DC_50_ & max. Deg) were extracted using GraphPad Prism’s 4-parameter logistic curve fitting.

The viability-window was determined by measuring the GI_50_ of viability based on the percentage of cells included in the viable gate and comparing it to the DC_50_ of degradation. The fold-change difference describing this window is [GI_50_ Viability / DC_50_ Degradation], where values above 1 indicate a positive (lower DC_50_ degradation than AC50 viability) window. If a compound did not have an DC_50_ or GI_50_, the top dose assessed was used for calculations.

### DCAF1 TR-FRET assay in standard binary format and cooperative format with BTK

Competition assay conditions consist of 20 µL total volume in white 384-well plates (Greiner), in 30 mM phosphate buffer pH7.4 containing 140 mM NaCl, 0.01% Tween20, 1 mM TCEP and 1% final DMSO. Tested compounds at 14 different concentrations were pre-incubated at 22◦C for 30 min with a large excess of 10 µM human BTK (G389-S659) in the cooperative assay or with buffer in standard binary assay. The following reagent were then added: 2 nM human DCAF1(1073-1399)_E1398S-Avi-His (biotinylated at the Cter), 0.75 nM cryptate terbium streptavidin (Perkin Elmer) and 10 nM bodipy labelled ligand (SMILES: CC1=CC(C)=C2C=C3C=CC(CCC(N4CCN(C5=CC6=NC(C(C)(C7=CC=C(C=C7)Cl)C)=NC(NCCN)=C6C =C5)CC4)=O)=N3=B(F)(N12)F) (internal preparation, KD = 69 ± 6 nM towards DCAF1 measured by fluorescence polarization). Samples were then incubated at 22◦C for 30 min before reading using a Tecan infinite M-1000 (excitation 340 nm, emission 490 and 520 nm). DCAF1 protein was omitted in the negative control. IC50 fitting was carried out using an in-house developed software (Novartis Helios software application, unpublished) using the methods described by Fomenko et al., 2006 (regression algorithms for nonlinear dose-response curve fitting). Following normalization of activity values for the wells to % inhibition (% inhibition= [(high control-sample)/ (high control-low control)] × 100). Data analysis could also be carried out using GraphPad Prism. The reported IC50 values are the average of at least 2 independent experiments.

### DCAF1 BTK Ubiquitination assay

To measure DCAF1 and PROTACs dependent BTK ubiquitination activity in vitro, we reconstituted the ubiquitination system using in-house produced and commercial proteins. The assay buffer consisted of 50mM Tris-HCl pH 7.4, 30mM NaCl, 10mM MgCl2, 0.2mM CaCl2, 0.01% TritonX, 1mM DTT, 20U/mL IPP and 0.05mM ATP. The biochemical assay was performed in 10µl volume in the following order of addition into the MTP: (1) 50nl of test compound in 100% DMSO. (2) 950nl of assay buffer. (3) 3µl of BTK-Tb-SA pre-mixture. (4) 3µl of E1-E2-Fluo-Ub pre-mixture. (5) 3µl of CRL4-DCAF1. The final concentrations of components in assay were: 100nM BTK, 4nM Tb-SA, 100nM E1, 2.5µM E2, 2.5µM Fluro-Ub, and 50nM CRL4-DCAF1. After addition of all components (in the order listed above), the mixture was read in kinetic mode with a PheraStar fluorescence reader. every 2-minutes on a using a standard TR-FRET protocol (LanthaScreen Optic module, Flash lamp Excitation source, 30 flashes per well).

#### BTK EPK assay

BTK kinase domain was mixed in assay buffer containing 50mM HEPES pH7.5, 0.02% Tween20, 0.02% Bovine Serum albumin, 1mM DTT, 0.01mM Na3VO4, 10mM b-Glycerolphosphat,18mM MgCl2, 1mM MnCl2 to a 2× concentration of 8nM. This solution was added to pre-dispensed compound dilution series in black, 384 well -low dead volume plates (Greiner) and the reaction was initiated by the addition of ATP and a BTK consensus peptide (Sequence: 5-Fluo-Ahx-TSELKKVVALYDY-Nle-P-Nle-NAND-NH2) to a final assay concentrations of 0.047mM and 0.002mM, respectively. of in assay buffer. The reaction was stopped after 60 minutes at 32℃ by addition of the same volume of spot solution containing 100mM HEPES pH7.5, 5% DMSO, 10mM EDTA, 0.015% Brij 35. Stopped reactions were analyzed on a Caliper LC3000, which was treated with the commercially available coating reagents CR-3 and CR-8 as instructed by the manufacturer. The relative intensities of phosphorylated peptide were calculated. Data was normalized to control wells containing 100mM EDTA solution instead of the 2× kinase solutions (0%) and DMSO-only control wells (100%). Normalized data was fitted using a sigmoidal fit.

### Intracellular BTK engagement NanoBRET

Apparent binding affinity of test compounds for BTK in intact and permeabilized cells was assessed under steady-state (end point) competition conditions by measuring their concentration-dependent reduction of the bioluminescence resonance energy transfer signal between BTK-NanoLuc (light donor) and Tracer-05 (Promega), a fluorescent BTK ATP-site binder (light acceptor), after 2 hours concurrent incubation, basically as described (Vasta et al. 2018) with minor adaptations. Thus, HEK293-A cells were batch-transfected by adding 1 μg pLenti6 BTK-Nanoluc vector and 9 μg Transfection Carrier DNA (pcDNA3.1 Hygro+) to 1 mL of Opti-MEM without phenol red (Thermo Fisher). After addition of 40 μL ViaFect (Promega) and incubation for 20 min, the DNA:lipid mixture was added to trypsinized HEK293-A cells suspended at 2 × 105/mL in 10 ml growth medium. 100 μL of the resulting cell:DNA:lipid suspension was then seeded at 2 × 104 cells/well into white, clear-bottom 96-well plates (Costar) that were pre-coated with poly-D-Lysine by incubation for 60 min with 30 μL of a sterile-filtered solution of 0.1 mg PDL (Mpbio) dissolved in phosphate-buffered saline, followed by a wash with 200 μL sterile phosphate-buffered and airdrying. After a 20 hr incubation in a humidified, 37°C/5% CO2 incubator, the cell supernatant was replaced with 95 μL serum- and phenol red-free Opti-MEM. Cells were then incubated for 2 hrs with serially diluted compounds added using an HP300 non-contact dispenser (Tecan), and 5 uL Tracer-05 (Vasta et al. 2018) pre-diluted in NanoBRET Dilution Buffer (12.5 mM HEPES, 31.25% PEG-300, pH 7.5) to achieve a final concentration of 0.5 μM. To measure BRET, NanoBRET NanoGlo substrate (Promega, final dilution of 1:1,000) and Extracellular NanoLuc Inhibitor (Promega, final concentration 20 μM) were added, followed by quantification of donor and acceptor luminescence emission on a Pherastar FX (BMG Labtech) multimode reader using 460 nm band-pass and 610 nm long-pass filters, respectively. To calculate raw BRET ratio values, the acceptor emission value was divided by the donor emission value for each sample. For background correction, the BRET ratio generated in the absence of tracer was subtracted from the BRET ratio of each sample. Raw BRET units were then converted to milliBRET units (mBU) by multiplying each raw BRET value by 1,000.

The mBU values for each treated sample was then divided by the average mBU value determined based on four DMSO-treated vehicle control samples and multiplied by 100 to arrive at % of vehicle-treated control. Four-parametric dose-response curve fits (model 203, non-fixed minima and maxima) were performed using the XLfit add-in for Excel (IDBS) to determine the compound concentration achieving half-maximal tracer displacement (IC50).

To enable parallel assessment of cellular and cell-free target engagement, all compounds were tested in the absence and presence of 50 μg/mL digitonin (Sigma), respectively.

### Generation of CRBN-BTK PROTAC resistant TMD8 cells for DCAF1-BTK PROTAC rescue studies

TMD8 cells were grown in the presence of a CRBN-BTK PROTAC CBt using a gradual dose increase. Start treatment was 0.1 nM for 6 weeks, followed by 3 weeks at 0.4 nM and another 4 weeks at 10 nM to finally obtain CBt resistant TMD8 cells (TMD8 CRBN resist.).

## QUANTIFICATION AND STATISTICAL ANALYSIS

For all experiments, the number of replicates, error bars, and statistical significance are defined in the relevant figure legends

## DATA AND CODE AVAILABILITY

The full list of E3 ligase with WDR motifs is attached as Supplementary Table 1. The raw data for the proteomics study is attached as Supplementary Table 2. The raw data for the arrayed CRISPR rescue screen is attached as Supplementary Table 4.

