## Supplementary Information for "Reinstating targeted protein degradation with DCAF1 PROTACs in CRBN PROTAC resistant settings"

### Table of Contents

**SUPPLEMENTARY FIGURES** .... S3

**Suppl. Figure 1**. Characterization of exit vector and chemical proteomics of compound 16 .... S3

**Suppl. Figure 2**. Genetic BRD9 degradation rescue screen .... S4

**Suppl. Figure 3**. Establishing a ratio-metric BTK-GFP assay in TMD8 cells .... S6

**Suppl. Figure 4**. Biochemical/biophysical characterization of DCAF1-based BTK degrader .... S8

**Suppl. Figure 5**. Validation of DBt-10 in TMD8 WT and TMD8 cells with acquired resistance to CRBN-based BTK degraders .... S10

**SUPPLEMENTARY TABLES** .... S12

**SUPPLEMENTARY METHODS: Chemistry** .... S14

**Synthesis of DBr-1** .... S14

**Synthesis of DDa-1** .... S19

**Synthesis of DBt-3** .... S23

**Synthesis of DBt-4** .... S27

**Synthesis of DBt-5** .... S30

**Synthesis of DBt-6** .... S34

**Synthesis of DBt-8** .... S40

**Synthesis of DBt-10** .... S45

**NMR spectra** .... S55

**High resolution MS spectra** .... S71

#### SUPPLEMENTARY FIGURES

**A**

**15**

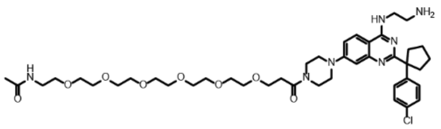

**B**

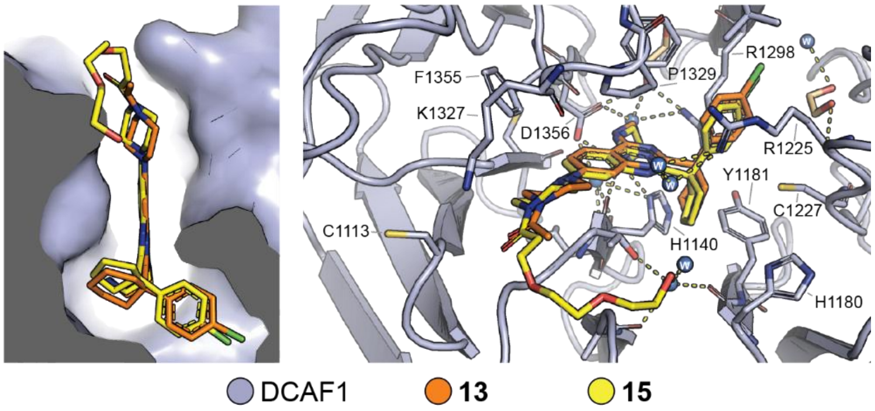


**C**

**DMSO vs.**

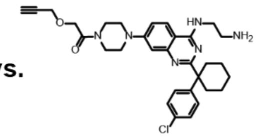

**16**

**HEK293T lysate on bead competition**

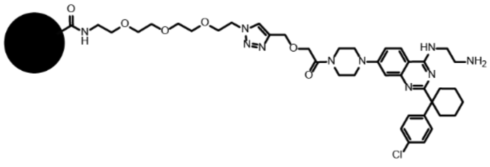

**Supplementary Figure 1.**

(A) Structure of PEGylated DCAF1 binders **15**.
(B) Left panel: Surface representation of the complex of DCAF1 with **13** (orange, pdb ID 8OO5), and **15** (yellow, pdb ID: 8OOD). Right panel: Detailed representation of the binding mode of **15** (yellow, pdb ID: 8OO5) overlayed with **13** (orange, pdb ID 8OOD). Hydrogen bonds are indicated as dashes.
(C) Structure and schematic of on-bead coupling of **16** for chemical proteomics.

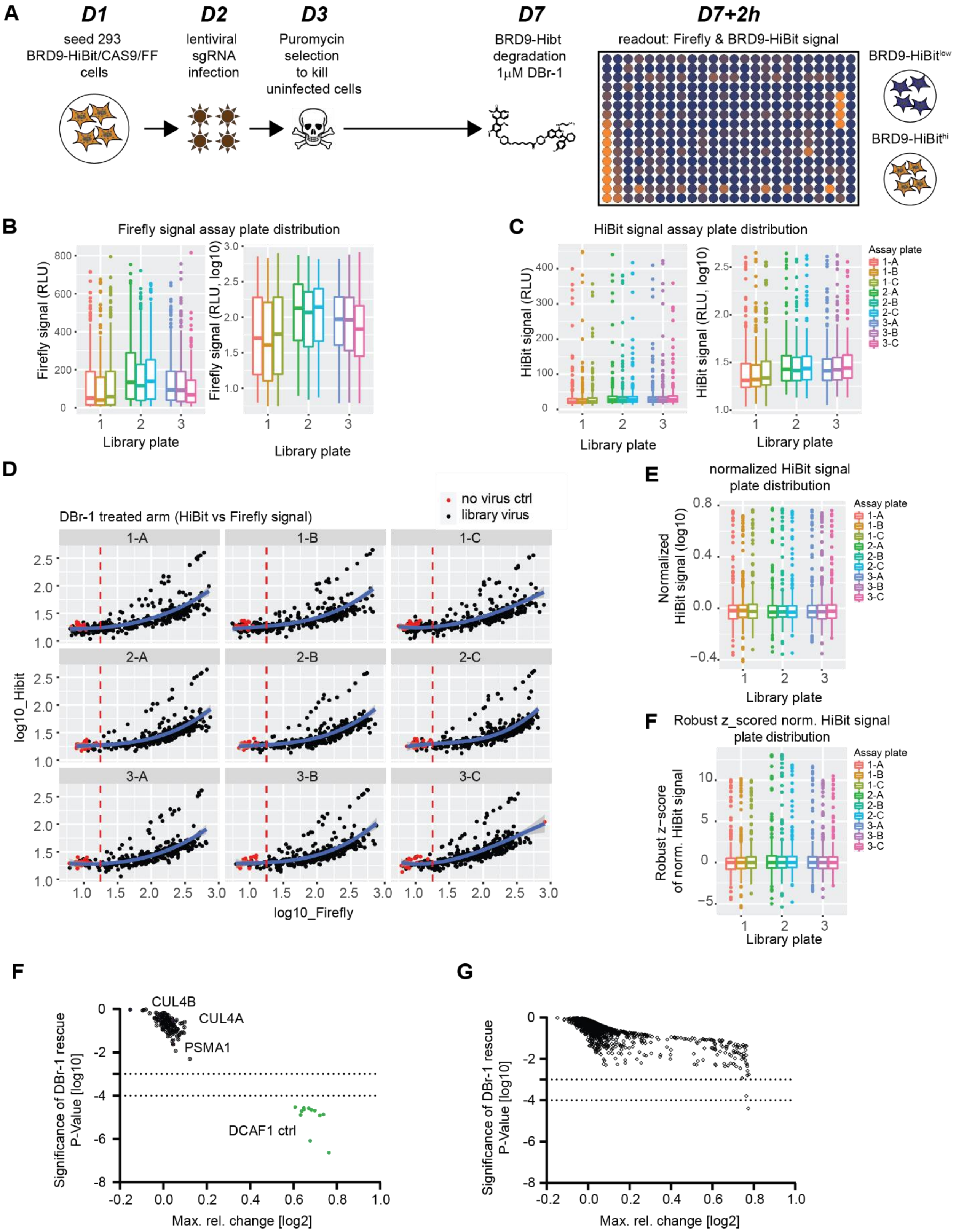
A
D1
seed 293 BRD9-HiBit/CAS9/FF cells
D2
lentiviral sgRNA infection
D3
Puromycin selection to kill uninfected cells
D7
BRD9-Hibt degradation 1μM DBr-1
D7+2h
readout: Firefly & BRD9-HiBit signal
BRD9-HiBit^low
BRD9-HiBit^hi
B
Firefly signal assay plate distribution
Firefly signal (RLU)
Firefly signal (RLU, log10)
Library plate
C
HiBit signal assay plate distribution
HiBit signal (RLU)
HiBit signal (RLU, log10)
Library plate
Assay plate
1-A
1-B
1-C
2-A
2-B
2-C
3-A
3-B
3-C
D
DBr-1 treated arm (HiBit vs Firefly signal)
no virus ctrl
library virus
log10_Hibit
log10_Firefly
E
normalized HiBit signal plate distribution
Normalized HiBit signal (log10)
Library plate
F
Robust z_scored norm. HiBit signal plate distribution
Robust z−score of norm. HiBit signal
Library plate
F
Significance of DBr-1 rescue P-Value [log10]
Max. rel. change [log2]
CUL4B
CUL4A
PSMA1
DCAF1 ctrl
G
Significance of DBr-1 rescue P-Value [log10]
Max. rel. change [log2]

**Supplementary Figure 2.**

(A) Schematic representation and timeline for Ubiquitin-sublibrary sgRNA rescue screen for DCAF1-BRD9 PROTAC (**DBr-1**).

(B) Relative Firefly luciferase signal per assay plate depicted as box plots. From each sgRNA library plate 3 assay plates were stamped.

(C) Relative HiBit luciferase signal per assay plate depicted as box plots. From each sgRNA library plate 3 assay plates were stamped.

(D) Dot plot depicting individual Firefly (x-axis) and HiBit (y-axis) signals per assay plate (A-C) stamped in triplicate from library plates (1-3), each dot represent an individual well. Red dots depict no virus ctrl wells and black dots represent a library or ctrl virus treated well of the assay plate. The red line at Firefly log10 1.25 represents the threshold for hit-calling. The blue line represents the assay plate based normalization curve (non-linear polynomial degree 3).

(E) Box plots representing normalized HiBit signal to Firefly signal based on a non-linear polynomial degree 3 relationship (see (D)). From each sgRNA library plate 3 assay plates were stamped.

(F) Box plots representing robust z-score from norm. HiBit signal (see (E)). From each sgRNA library plate 3 assay plates were stamped.

(G) Ubiquitin sgRNA sublibrary rescue scores from (**DBr-1**) treatment plotted as significance of rescue P-value (y-axis) vs max. rel. change. Dotted lines at -4 and -3 P-value indicate strong and weaker hits with a false-discovery rate of 7%. Black dots represent sgRNA treatments of wells that did not pass the Firefly signal threshold of > log10 1.25, CUL4A, CUL4B and PSMA1 are indicated. Green dots represent DCAF1 ctrl sgRNAs distributes on each assay plate.

(H) Randomized run of Ubiquitin sgRNA sublibrary screen results plotted as significance of rescue P-value (y-axis) vs max. rel. change. Dotted lines at -4 and -3 P-value indicate strong and weaker hits.

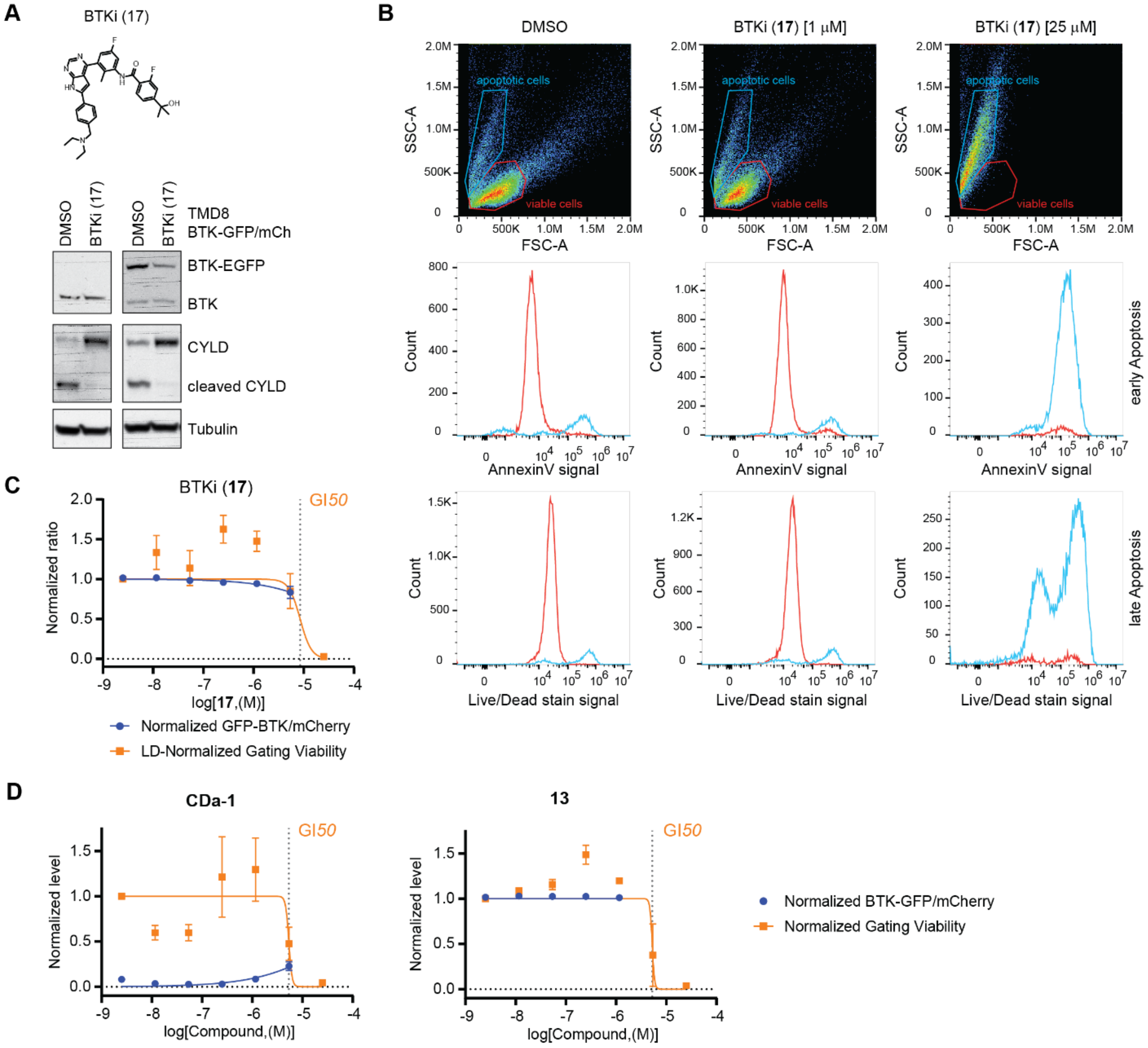


| Compound ID | Degradation BTK-GFP/mCherry max. Deg | Abs. DC50 [µM] | DC50 [µM] | Viable Gate Inclusion Abs. GI50 [µM] | GI50 [µM] | Max concentration [µM] Gating Viability >25% | Avg #single cells Max concentraion | V/D (GI50 Viability / DC50 Degradation) |
|---|---|---|---|---|---|---|---|---|
| (**17**) | 17% | | >25 | 6.817 | 6.748 | 5.39 | 9197 | 0.3 |
| (**13**) | 0% | | >25 | 5.22 | 5.044 | 1.16 | 15731 | 0.2 |
| (**CDa-1**) | 97% | <0.0025 | <0.0025 | 9.882 | 5.312 | 5.39 | 6225 | 2124.8 |

**Supplementary Figure 3.**

(A) Structure of BTKi (17) and characterization of TMD8 and TMD8 BTK-GFP/mCh cell lines after (17) treatment for 24 hours.

(B) Dot plot representing Forward (FSC-A, x-axis) versus Sideward (SSC-A, y-axis) scattering of cells measured by flow cytometry treated with (**17**) at concentrations of 1 and 25 µM or DMSO as control. Each dot represents an individual cell, color code represents cell density in plot where single dot resolution was not possible (the warmer the color the more cells per plot region). Selection gates for viable (red) and apoptotic (blue) cells are indicated (first row). Histogram plots based on above gating strategy depicting individual cell count versus signal of an early and a late apoptotic marker (Annexin

V, middle row; Live/Dead stain, bottom row, respectively). Viable populations are plotted as red curves and apoptotic populations as blue curves.

(C) Degradation of BTK-GFP in TMD8 BTK-GFP/mCh cells after 24h (**17**) treatment displayed as normalized rel. change of the ratio between BTK-GFP and mCherry (mCh) signals (blue curve), Viability after 24h (**DDa-1**) treatment is displayed as rel. change in cellular distribution between viable and apoptotic FSC/SSC gate (see (B)). DC50, GI50 and V/D is shown in (D).

(D) Degradation and viability of BTK-GFP in TMD8 BTK-GFP/mCh cells after 24h (**CDa-1**) or (**13**) treatment as in (C). Table depicting parameters extracted from flowcytometry measurements of TMD8 BTK-GFP/mCh cells treated with BTKi (**17**), DCAF1 binder (**13**) and CRBN-Das PROTAC (CDa-1).

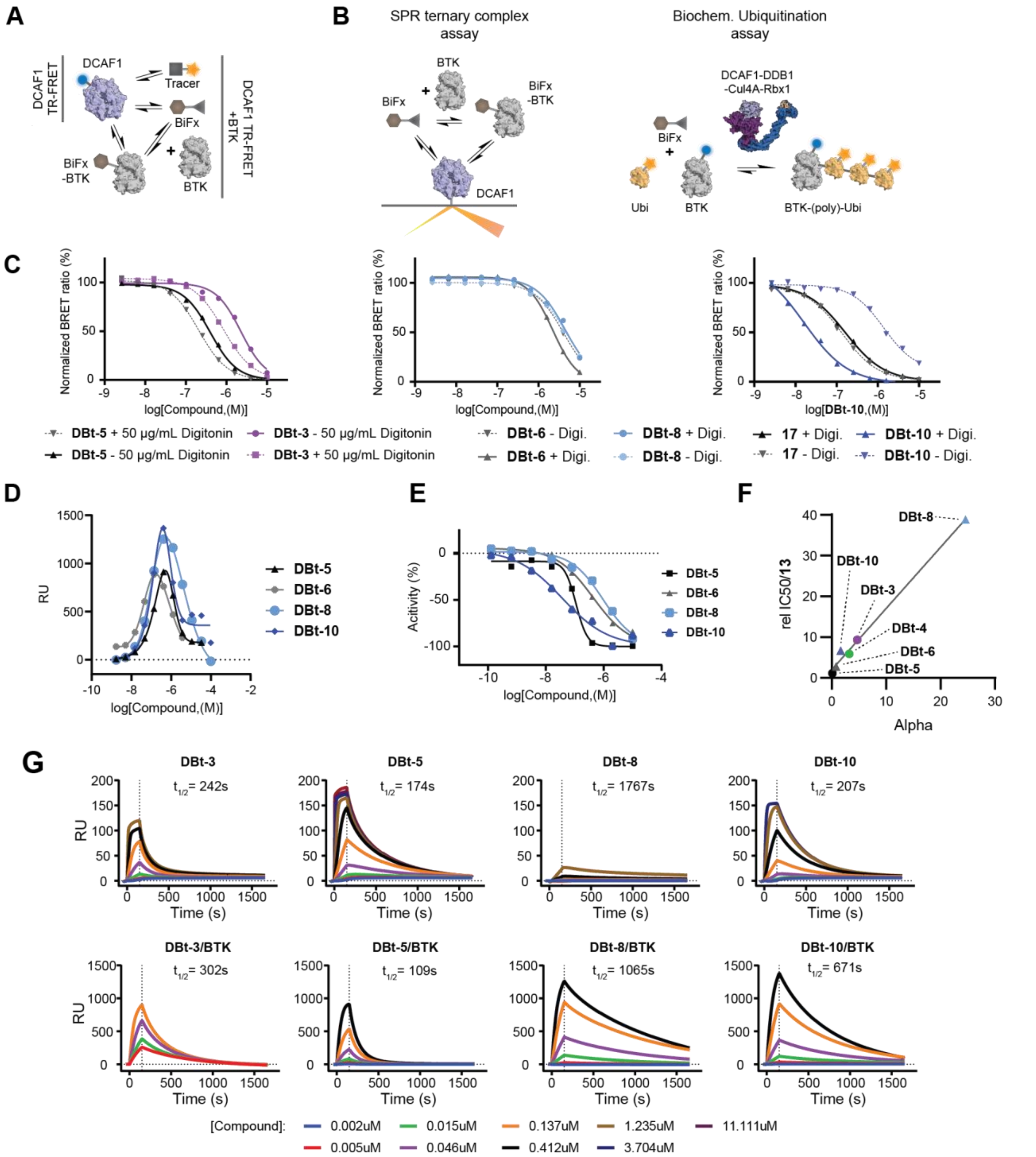


**Supplementary figure 4.**

(A) Schematic depiction of the DCAF1 TR-FRET and Alpha DCAF1 TR-FRET assay. Table with results of assays.

(B) Schematic illustration of the SPR ternary complex assay and the biochemical ubiquitination assay based on a Ubiquitin-BTK TR-FRET pair.

(C) Cellular BTK target engagement assays in the absence or presence of the cell permeabilizing agent Digitonin (IC50 values are displayed in Supplementary table 7).

(D) Overlay of SPR ternary complex formation assays (.

(E) Characterization of PROTACs in an biochemical BTK enzymatic assay (IC50 values are displayed in Supplementary table 7).

(F) Correlation of the relative DCAF1 TR-FRET $IC_{50}$ values of the PROTAC molecules DBt-1 - 5 normalized to the $IC_{50}$ value of the parent DCAF1 binder (**13**) with the calculated alpha factor of the Alpha DCAF1 TR-FRET assay (additional presence of BTK). Circles indicate PEG-based linkers while triangles mark other linkers or a different BTK ligand for DBt-10.

(G) SPR sensograms of either compounds measured alone or in presence of 0.2uM BTK against immobilized DCAF1 (WD40). Additionally, the residence times of the binary or ternary complexes (t1/2) are shown.

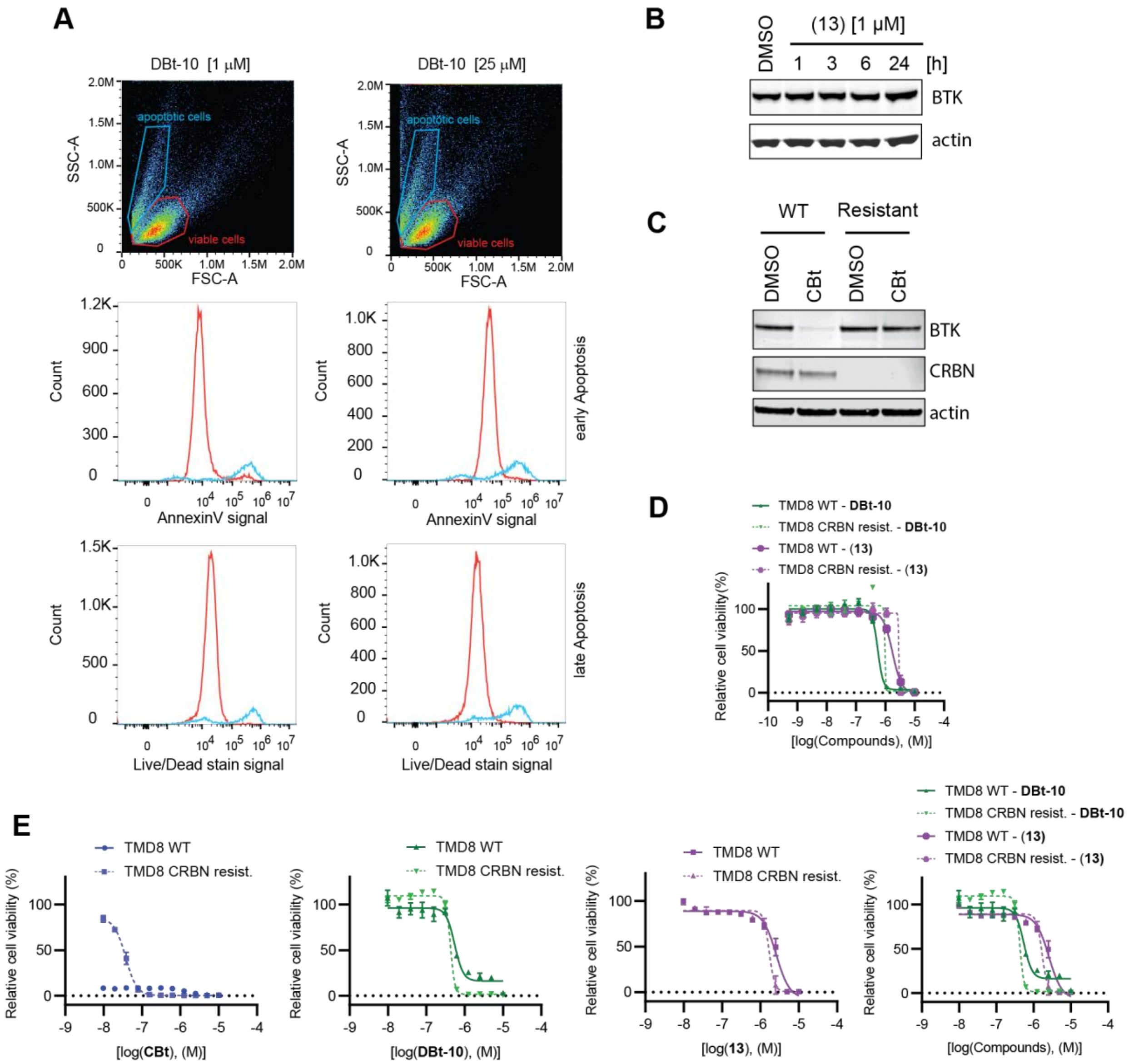


**Supplementary figure 5.**

(A) Dot plot representing Forward (FSC-A, x-axis) versus Sideward (SSC-A, y-axis) scattering of cells measured by flow cytometry treated with DBt-10 at concentrations of 1 and 25 µM. Each dot represents an individual cell, color code represents cell density in plot where single dot resolution was not possible (the warmer the color the more cells per plot region). Selection gates for viable (red) and apoptotic (blue) cells are indicated (first row). Histogram plots based on above gating strategy depicting individual cell count versus signal of an early and a late apoptotic marker (Annexin V, middle row; Live/Dead stain, bottom row, respectively). Viable cells are plotted as red curves and apoptotic cells as blue curves.

(B) Immunoblot analysis of TMD8 cells treated with 1 µM (13) for 1, 3, 6 and 24 h time points.

(C) Characterization of parental and CRBN resistant TMD8 cells probing for CRBN and BTK after 24h CBt treatment. Actin was used as loading control.

(D) Viability measured by cell titer-Glo for DBt-10 (green) vs (13) (purple) in WT (solid) vs CRBN resist. (dotted) TMD8 cells.

(E) Repeat of the viability measured by cell titer-Glo for DBt-10 (green) vs (13) (purple) in WT (solid) vs CRBN resist. (dotted) TMD8 cells with adjusted dose response range.

**SUPPLEMENTARY TABLES**

Suppl. Tables 1-7 can be found in additional Supplementary file

Supplementary table 1: Genes included in the DepMap analysis

Supplementary table 2: Raw data of chemical proteomics experiment in HEK293T cells with compound **16**

Supplementary table 3: Details of the sgRNA library used in BRD9 degradation rescue experiment

Supplementary table 4: Raw data of the BRD9 degradation rescue experiment

Supplementary table 5: Genes classified as hits in the BRD9 degradation rescue experiment

Supplementary table 6: Raw data of the proteomics experiment comparing DDa-1 and compound 13

Supplementary table 7: Characterization data of the DCAF1 based BTK degrader and control compounds

Supplementary table 8: Data collection and refinement statistics of the DCAF1-ligand complex X-ray structures

**Supplementary table 8.** Data collection and refinement statistics.

| Complex | DCAF1-VHF543 | DCAF1-IVH258 |
|---|---|---|
| **PDB accession code** | **8OO5** | **8OOD** |
| **Beamline** | **SLS PXII** | **SLS PXII** |
| ***Data Collection*** | | |
| Spacegroup | $P6_122$ | $P6_122$ |
| Cell dimensions | $a = b = 81.6$ Å, $c = 234.1$ | $a$ = b = 81.5 Å, $c$ = 232.7 Å |
| | $\alpha = \gamma = 90.0^{\circ}$; $\beta = 120^{\circ}$ | $\alpha = \gamma = 90.0^{\circ}$; $\beta = 120^{\circ}$ |
| Resolution (Å) [a] | 52.38-2.25 (2.32-2.25) | 52.20-1.50 (1.52-1.50) |
| No. unique reflections [a] | 22,902 (2,066) | 74,179 (3,405) |
| Completeness[a] (%) | 100.0 (100.0) | 99.6 (94.0) |
| I/σI[a] | 11.4 (2.2) | 11.1 (1.1) |
| $R_{merge}$[a] | 0.280 (2.821) | 0.091 (1.726) |
| CC (1/2) [a] | 0.996 (0.771) | 0.997 (0.452) |
| Redundancy[a] | 21.8 (22.9) | 8.7 (7.9) |
| ***Refinement*** | | |
| No. atoms in refinement (P/L/W)[b] | 2,383/88/77 | 2,570/136/316 |
| B factor (P/L/W)[b] (Å$^2$) | 43/58/45 | 21/32/33 |
| $R_{fact}$ (%) | 18.1 | 17.0 |
| $R_{free}$ (%) | 22.0 | 18.6 |
| Rmsd [c] (Å) | 0.017 | 0.021 |
| Rmsd [c] angle (°) | 2.08 | 2.5 |
| ***Molprobity Ramachandran*** | | |
| Favoured (%) | 95.95 | 97.30 |
| Outliers (%) | 0.34 | 0 |

[a] Values in brackets show the statistics for the highest resolution shell.

[b] P/L/W indicate protein, ligand molecules, and water molecules, respectively.

[c] Rmsd indicates root-mean-square deviation

**SUPPLEMENTARY METHODS: Chemistry Experimental**

**Compound DDa-1 Synthetic Methods:**

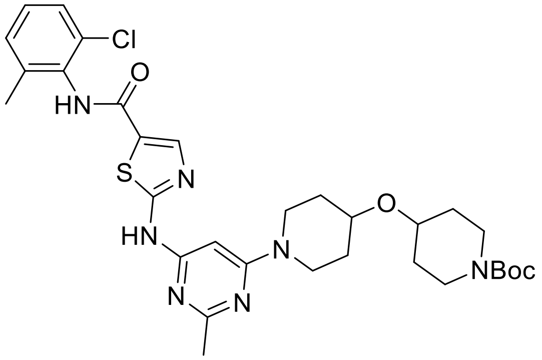

**tert-butyl 4-((1-(6-((5-((2-chloro-6-methylphenyl)carbamoyl)thiazol-2-yl)amino)-2-methylpyrimidin-4-yl)piperidin-4-yl)oxy)piperidine-1-carboxylate**

Tert-butyl 4-(piperidin-4-yloxy)piperidine-1-carboxylate (0.11 g, 3 Eq, 0.38 mmol) was added to a suspension of 2-((6-chloro-2-methylpyrimidin-4-yl)amino)-N-(2-chloro-6-methylphenyl)thiazole-5-carboxamide (50 mg, 1 Eq, 0.13 mmol) and N,N-Diisopropylethylamine (33 mg, 44 µL, 2 Eq, 0.25 mmol) dissolved in 1,4-Dioxane (1.2 mL) at room temperature. The reaction was heated on a hot plate to 90 °C and stirred. After 24 hours, the material was diluted with water and ethyl acetate, and the layers were separated. The aqueous layer was washed with 3 x ethyl acetate and the organic layers were combined. The combined organics were dried with sodium sulfate, filtered, and concentrated under reduced pressure. Purification by column chromatography (ISCO, 12 g silica column, 0-15 % MeOH/DCM) gave the desired product (57.1 mg, 88 mmol, 69 % yield) as an off-white solid.

**$^1$H NMR** (400 MHz, DMSO) δ 11.42 (s, 1H), 9.87 (s, 1H), 8.22 (s, 1H), 7.40 (dd, *J* = 7.5, 2.0 Hz, 1H), 7.34 – 7.22 (m, 2H), 6.08 (s, 1H), 3.90 (s, 2H), 3.72 (s, 1H), 3.69 – 3.58 (m, 4H), 3.28 – 3.19 (m, 3H), 3.02 (s, 2H), 2.40 (s, 4H), 2.33 (s, 1H), 2.24 (s, 4H), 1.84 (s, 2H), 1.77 (d, *J* = 12.9 Hz, 3H), 1.40 (s, 12H).

**LCMS:** 642.5 [M+H$^+$].

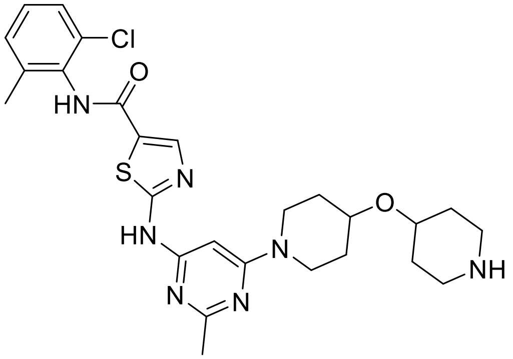

**N-(2-chloro-6-methylphenyl)-2-((2-methyl-6-(4-(piperidin-4-yloxy)piperidin-1-yl)pyrimidin-4-yl)amino)thiazole-5-carboxamide**

tert-butyl 4-((1-(6-((5-((2-chloro-6-methylphenyl)carbamoyl)thiazol-2-yl)amino)-2-methylpyrimidin-4-yl)piperidin-4-yl)oxy)piperidine-1-carboxylate (111.2 mg, 1 Eq, 173.1 µmol) was suspended in dichloromethane (1.5 mL) at room temperature. HCl (4M in Dioxane) (44.19 mg, 303.0 µL, 4 molar, 7 Eq, 1.212 mmol) was subsequently added, and the reaction was stirred at room temperature while monitored by LCMS. After 4.5 hours of stirring, LCMS revealed full conversion to the desired product. The material was concentrated under reduced pressure and further dried under high vacuum overnight. The desired product was isolated (100.2 mg, 0.16 mmol, quantitative yield) as a light orange oil and used without further purification.

**1H NMR** (400 MHz, DMSO) δ 10.10 (s, 1H), 8.96 (s, 2H), 8.35 (d, J = 7.5 Hz, 1H), 7.40 (dd, J = 7.4, 2.0 Hz, 1H), 7.32 – 7.21 (m, 2H), 3.91 (s, 2H), 3.81 – 3.66 (m, 3H), 3.49 – 3.36 (m, 2H), 3.14 (s, 3H), 2.94 (s, 3H), 2.54 (s, 3H), 2.24 (s, 3H), 2.01 – 1.84 (m, 5H), 1.75 – 1.61 (m, 3H), 1.56 (d, J = 30.8 Hz, 2H).

**LCMS:** 542.3 [M+H$^+$].

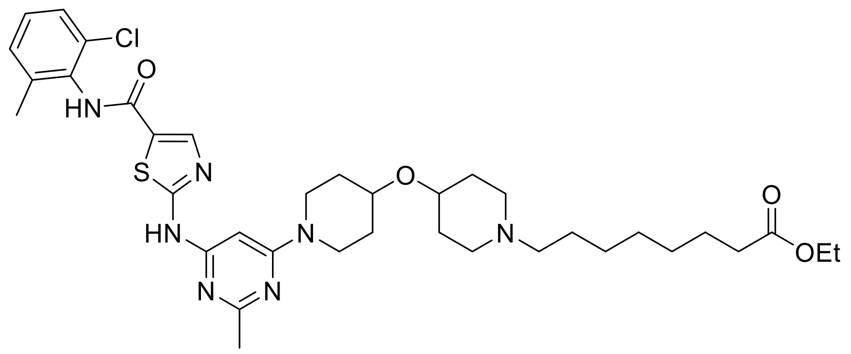


**ethyl 8-(4-((1-(6-((5-((2-chloro-6-methylphenyl)carbamoyl)thiazol-2-yl)amino)-2-methylpyrimidin-4-yl)piperidin-4-yl)oxy)piperidin-1-yl)octanoate**

Ethyl 8-bromooctanoate (139.3 mg, 3 Eq, 554.5 µmol) was added to a solution of N-(2-chloro-6-methylphenyl)-2-((2-methyl-6-(4-(piperidin-4-yloxy)piperidin-1-yl)pyrimidin-4-yl)amino)thiazole-5-carboxamide (100.2 mg, 1 Eq, 184.8 µmol) and N,N-Diisopropylethylamine (119.5 mg, 161 µL, 5 Eq, 924.2 µmol) dissolved in DMF (1 mL). The reaction was heated to 80°C on a hot plate and monitored by LCMS. After 24 hours of stirring, the crude material was adsorbed onto a bed of celite and purified by column chromatography (ISCO, 12 g silica column, 0-20 % MeOH/DCM). The desired product was isolated as a dark orange solid (100 mg, 0.15 mmol, 80 % yield)

$^1$H NMR (400 MHz, $CDCl_3$) δ 11.21 (s, 2H), 7.29 (s, 1H), 7.20 – 7.12 (m, 1H), 4.10 (q, *J* = 7.1 Hz, 1H), 3.65 (pd, *J* = 6.7, 4.0 Hz, 6H), 3.31 (d, *J* = 11.5 Hz, 1H), 3.08 (dt, *J* = 7.4, 3.7 Hz, 5H), 2.88 (q, *J* = 5.6 Hz, 1H), 2.51 (s, 3H), 2.35 (s, 2H), 2.27 (t, *J* = 7.5 Hz, 1H), 1.87 (s, 4H), 1.59 (d, *J* = 6.4 Hz, 2H), 1.56 (s, 2H), 1.54 (s, 3H), 1.51 (s, 4H), 1.48 (d, *J* = 6.5 Hz, 1H), 1.44 (s, 4H), 1.32 (d, *J* = 4.8 Hz, 4H), 1.24 (t, *J* = 7.1 Hz, 2H).

LCMS: 712.7 [M+H+].

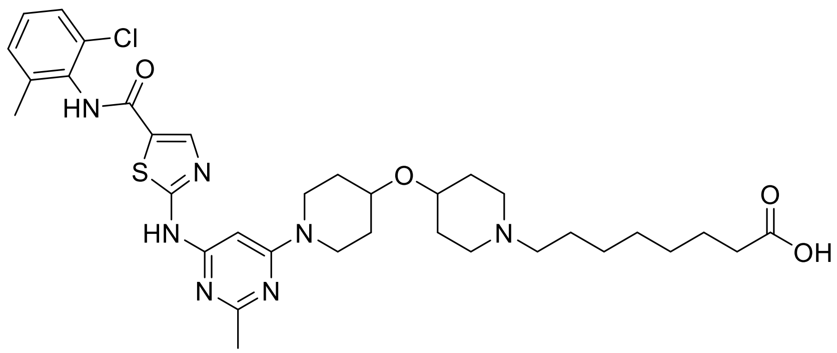

**8-(4-((1-(6-((5-((2-chloro-6-methylphenyl)carbamoyl)thiazol-2-yl)amino)-2-methylpyrimidin-4-yl)piperidin-4-yl)oxy)piperidin-1-yl)octanoic acid**

Ethyl 8-(4-((1-(6-((5-((2-chloro-6-methylphenyl)carbamoyl)thiazol-2-yl)amino)-2-methylpyrimidin-4-yl)piperidin-4-yl)oxy)piperidin-1-yl)octanoate (100.0 mg, 1 Eq, 140.4 µmol) was dissolved in THF (800.0 µL) and Water (533.3 µL). Lithium hydroxide (6.724 mg, 2 Eq, 280.8 µmol) was added in one portion and the reaction was stirred at room temperature. After 3 hours of stirring, LCMS revealed full conversion to the desired carboxylic acid. The crude material was concentrated under vacuum and further dried by high vac overnight. The product was used in the subsequent reaction without further purification.

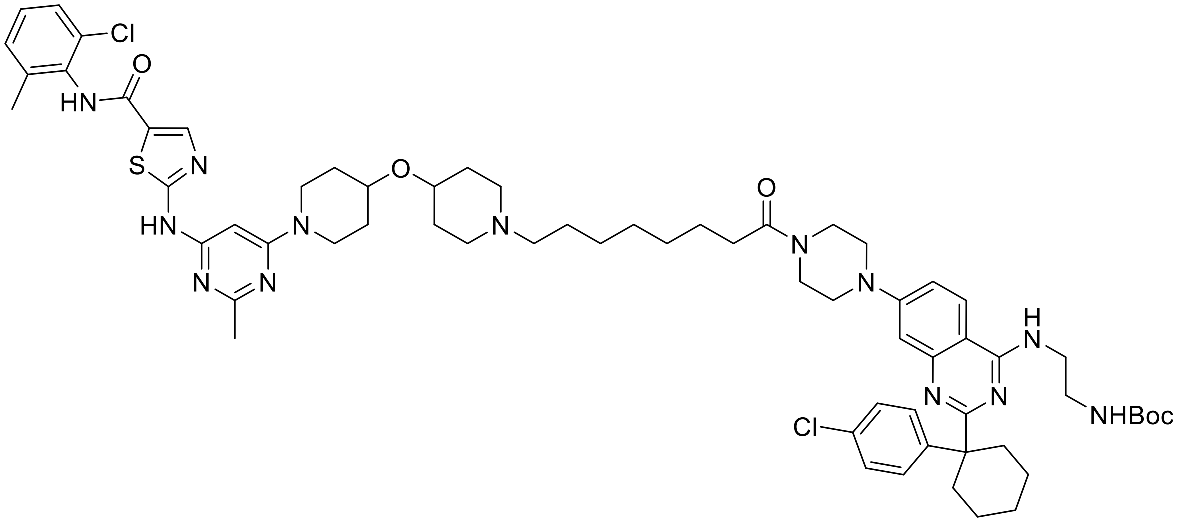

**tert-butyl (2-((7-(4-(8-(4-((1-(6-((5-((2-chloro-6-methylphenyl)carbamoyl)thiazol-2-yl)amino)-2-methylpyrimidin-4-yl)piperidin-4-yl)oxy)piperidin-1-yl)octanoyl)piperazin-1-yl)-2-(1-(4-chlorophenyl)cyclohexyl)quinazolin-4-yl)amino)ethyl)carbamate**

In a pressure-release vial, dissolved 8-(4-((1-(6-((5-((2-chloro-6-methylphenyl)carbamoyl)thiazol-2-yl)amino)-2-methylpyrimidin-4-yl)piperidin-4-yl)oxy)piperidin-1-yl)octanoic acid (44.38 mg, 1 Eq, 64.85

µmol), HATU (49.32 mg, 2 Eq, 129.7 µmol) and N,N-Diisopropylethylamine (33.53 mg, 45.2 µL, 4 Eq, 259.4 µmol) in DMF (500.0 µL). The reaction was stirred at room temperature for 10 minutes before tert-butyl (2-((2-(1-(4-chlorophenyl)cyclohexyl)-7-(piperazin-1-yl)quinazolin-4-yl)amino)ethyl)carbamate (36.65 mg, 1 Eq, 64.85 µmol) was added in one portion. After five hours of stirring, the reaction was purified by reverse-phase flash chromatography (Waters XBridge C18 OBD 30 x 50 mm column, 65% - 95% ACN/water with 5 mM $NH_4OH$ modifier at 75 mL/min, 1.5mL injection). The fractions were lyophilized to afford the desired product (16.70 mg, 13 µmol, 21 % yield) as a white fluffy solid.

$^1$H NMR (400 MHz, DMSO) δ 11.41 (s, 1H), 9.86 (s, 1H), 8.21 (s, 1H), 8.02 – 7.82 (m, 2H), 7.47 (d, *J* = 8.4 Hz, 2H), 7.40 (dd, *J* = 7.6, 2.0 Hz, 1H), 7.33 – 7.17 (m, 5H), 6.91 (t, *J* = 5.7 Hz, 1H), 6.85 (d, *J* = 2.4 Hz, 1H), 6.07 (s, 1H), 3.90 (d, *J* = 12.9 Hz, 2H), 3.65 (s, 1H), 3.59 (d, *J* = 5.5 Hz, 4H), 3.46 (d, *J* = 6.2 Hz, 2H), 3.41 (d, *J* = 9.2 Hz, 1H), 3.29 (s, 4H), 3.26 – 3.12 (m, 5H), 2.80 (d, *J* = 12.8 Hz, 2H), 2.66 (d, *J* = 10.2 Hz, 2H), 2.40 (s, 3H), 2.34 (t, *J* = 7.4 Hz, 3H), 2.22 (d, *J* = 13.5 Hz, 5H), 2.03 – 1.88 (m, 4H), 1.77 (d, *J* = 15.6 Hz, 5H), 1.50 (dd, *J* = 15.2, 8.5 Hz, 7H), 1.37 (s, 11H), 1.25 (d, *J* = 16.1 Hz, 7H).

**LCMS**: 1231.2 [M+H$^+$].

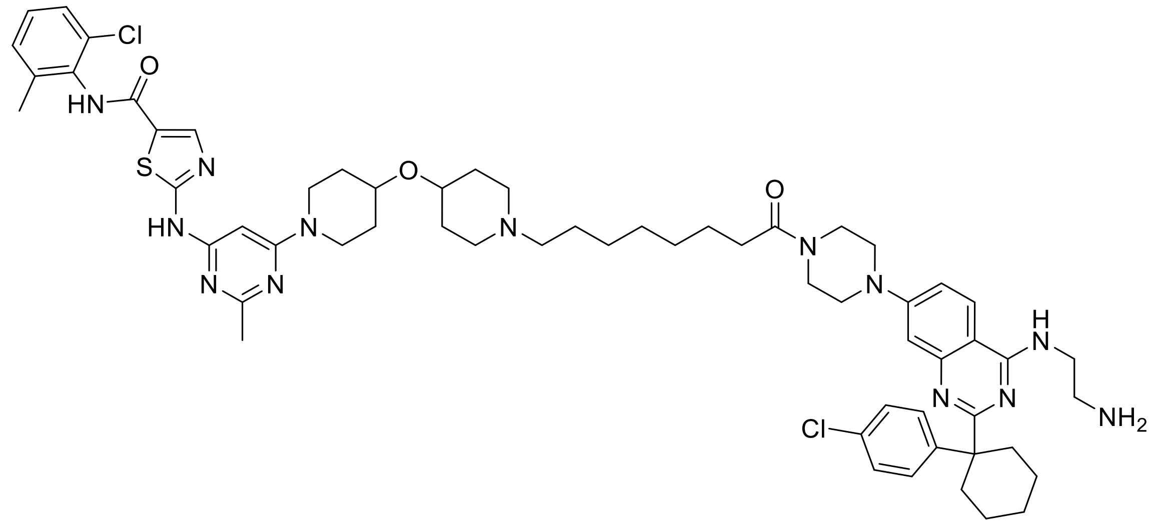

**2-((6-(4-((1-(8-(4-(4-((2-aminoethyl)amino)-2-(1-(4-chlorophenyl)cyclohexyl)quinazolin-7-yl)piperazin-1-yl)-8-oxooctyl)piperidin-4-yl)oxy)piperidin-1-yl)-2-methylpyrimidin-4-yl)amino)-N-(2-chloro-6-methylphenyl)thiazole-5-carboxamide**

In a vial, tert-butyl (2-((7-(4-(8-(4-((1-(6-((5-((2-chloro-6-methylphenyl)carbamoyl)thiazol-2-yl)amino)-2-methylpyrimidin-4-yl)piperidin-4-yl)oxy)piperidin-1-yl)octanoyl)piperazin-1-yl)-2-(1-(4-chlorophenyl)cyclohexyl)quinazolin-4-yl)amino)ethyl)carbamate (14.80 mg, 1 Eq, 12.02 µmol) was dissolved in dichloromethane (100 µL) at room temperature. HCl (4M in Dioxane) (2.191 mg, 15.02 µL, 4 molar, 5 Eq, 60.09 µmol) was added dropwise to the vial, and the reaction continued to be stirred at room temperature. After 4 hours, the material was concentrated under reduced pressure and further dried overnight under high vacuum overnight to afford the final product (14.3 mg, 11 µmol, quantitative yield) as a white solid.

$^{1}H$ NMR (400 MHz, DMSO) δ 13.39 (d, J = 5.2 Hz, 1H), 10.75 (s, 1H), 10.26 (s, 1H), 10.13 (s, 1H), 8.47 (d, J = 9.4 Hz, 1H), 8.37 (s, 3H), 7.67 (d, J = 4.0 Hz, 1H), 7.57 (dd, J = 8.7, 1.6 Hz, 2H), 7.41 – 7.36 (m, 4H), 7.31 – 7.22 (m, 2H), 3.94 (dd, J = 37.3, 11.0 Hz, 4H), 3.76 – 3.60 (m, 5H), 3.50 (ddd, J = 4.9, 4.1, 1.4 Hz, 3H), 3.48 – 3.36 (m, 6H), 3.26 (d, J = 11.7 Hz, 1H), 3.16 (d, J = 6.1 Hz, 2H), 2.96 (q, J = 12.9 Hz, 5H), 2.54 (s, 3H), 2.35 (t, J = 7.4 Hz, 2H), 2.24 (s, 3H), 2.12 (d, J = 6.0 Hz, 2H), 2.03 (s, 2H), 1.89 (s, 3H), 1.74 – 1.58 (m, 4H), 1.46 (d, J = 40.0 Hz, 8H), 1.32 (d, J = 17.9 Hz, 8H), 0.92 – 0.82 (m, 1H).

$^{13}C$ NMR (151 MHz, DMSO) δ 170.97, 165.56, 162.50, 159.10, 156.34, 154.24, 154.00, 142.80, 141.20, 138.76, 137.29, 133.34, 132.41, 131.77, 131.72, 131.62, 129.06, 128.88, 128.66, 128.53, 128.29, 128.23, 127.93, 127.04, 125.99, 125.01, 116.05, 102.11, 98.65, 72.17, 70.53, 70.17, 69.61, 60.18, 55.57, 55.08, 49.94, 48.96, 47.04, 46.21, 43.96, 43.64, 37.87, 34.13, 32.18, 30.71, 29.79, 28.88, 28.53, 28.36, 27.08, 26.08, 24.93, 24.55, 23.25, 23.01, 22.66, 18.34, 13.91, 10.82.

LCMS: 1130.8 [M+H$^{+}$].

**Compound DBr-1 Synthetic Methods:**

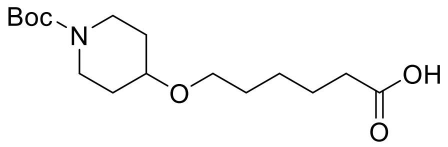


**6-((1-(tert-butoxycarbonyl)piperidin-4-yl)oxy)hexanoic acid**

To a solution of tert-butyl 4-((6-methoxy-6-oxohexyl)oxy)piperidine-1-carboxylate* (15.0 g, 1.0 Eq, 45.5 mmol) in a mixture of MeOH (100 mL), THF (100 mL) and water (100 mL) at 20 °C was added lithium hydroxide hydrate (19.0 g, 10.0 Eq, 455 mmol) and the reaction mixture was heated at 80 °C for 2 hours. The mixture was concentrated, water (100 mL) was added and the pH was adjusted to 3 by the addition of a concentrated aqueous solution of citric acid. The mixture was extracted with ethyl acetate (2 x 200 mL), the combined organic phases were washed with brine (200 mL), dried over sodium sulfate and concentrated. The residue was purified by silica gel column chromatography, eluted with ethyl acetate (from 10 % to 15 %) in petrol ethers, yielding the title compound 6-((1-(tert-butoxycarbonyl)piperidin-4-yl)oxy)hexanoic acid as an yellow oil (12.4 g, 39.3 mmol, 86 %).

**$^{1}$H NMR** (400 MHz, DMSO-$D_6$) δ 12.05 – 11.84 (m, 1H), 3.66 – 3.53 (m, 2H), 3.10 – 2.89 (m, 2H), 2.24 – 2.14 (m, 2H), 1.80 – 1.69 (m, 2H), 1.55 – 1.43 (m, 4H), 1.43 – 1.35 (m, 9H), 1.35 – 1.25 (m, 4H).

**LCMS:** 216.2 [M+$H^+$-Boc].

* synthesis described in patent WO2020/181050, page 145.

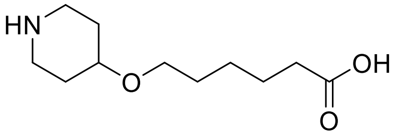


**6-(piperidin-4-yloxy)hexanoic acid**

To a solution of 6-((1-(tert-butoxycarbonyl)piperidin-4-yl)oxy)hexanoic acid (1.0 g, 1.0 Eq, 3.17 mmol) in dichloromethane (10 mL) was added a solution of HCl in 1,4-dioxane (10 mL, 4 M), the reaction mixture was stirred at 30 °C for 1 hour and concentrated under reduced pressure, yielding the title compound 6-(piperidin-4-yloxy)hexanoic acid hydrochloride salt as a yellow solid (780 mg, 3.1 mmol, 98 %), which was used for the next step without further purification.

**Mp:** 104 – 107 °C

**$^{1}$H NMR** (400 MHz, DMSO-$D_6$) δ 11.99 (br s, 1H), 8.98 – 8.72 (m, 2H), 3.60 – 3.46 (m, 1H), 3.41 – 3.35 (m, 2H), 3.14 – 3.03 (m, 2H), 2.93 (br s, 2H), 2.20 (br t, J = 7.3 Hz, 2H), 2.04 – 1.81 (m, 2H), 1.65 (br d, J = 8.9 Hz, 2H), 1.55 – 1.43 (m, 4H), 1.39 – 1.23 (m, 2H).

**$^{13}$C NMR** (100 MHz, DMSO-D$_6$) δ 175.0, 71.0, 67.7, 41.0, 34.2, 29.8, 27.9, 25.9, 24.9.

**LCMS:** 216 [M+H$^+$].

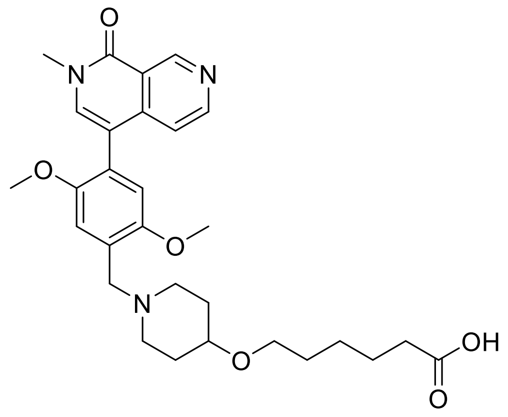

**6-((1-(2,5-dimethoxy-4-(2-methyl-1-oxo-1,2-dihydro-2,7-naphthyridin-4-yl)benzyl)piperidin-4-yl)oxy)hexanoic acid**

To a solution of 6-(piperidin-4-yloxy)hexanoic acid (1.0 g, 1.0 Eq, 4.64 mmol) and 2,5-dimethoxy-4-(2-methyl-1-oxo-1,2-dihydro-2,7-naphthyridin-4-yl)benzaldehyde* (1.51 g, 1.0 Eq, 4.64 mmol) in ethanol (20 mL) was added diisopropyl ethylamine (1.8 g, 3.0 Eq, 13.9 mmol) and a solution of zinc chloride in THF (4.6 mL, 1 M, 1.0 Eq, 4.64 mmol) and the reaction mixture was stirred at 30 °C for 1 hour. Solid sodium cyanoborohydride (877 mg, 3.0 Eq, 13.9 mmol) was added and stirring was continued at 30 °C for 15 hours. Water (20 mL) was added, the mixture was extracted with dichloromethane (3 x 30 mL), the combined organic phases were washed with brine (2 x 10 mL) and dried over sodium sulfate. The residue was purified by preparative HPLC on a Phenomenex Luna C18 column (150 x 40 mm, 15 μm) eluting with acetonitrile (from 10 % to 40 %) in an aqueous solution of $NH_4HCO_3$ (0.1 %). Fractions containing the title compound were combined and freeze dried yielding the title compound 6-((1-(2,5-dimethoxy-4-(2-methyl-1-oxo-1,2-dihydro-2,7-naphthyridin-4-yl)benzyl)piperidin-4-yl)oxy)hexanoic acid as a white solid (600 mg, 1.14 mmol, 25 %).

**$^1$H NMR** (400 MHz, DMSO-D$_6$) δ 9.40 (s, 1H), 8.63 (d, J = 5.6 Hz, 1H), 7.73 (s, 1H), 7.13 (s, 1H), 7.03 (d, J = 5.6 Hz, 1H), 6.91 (s, 1H), 3.74 (s, 3H), 3.63 (s, 3H), 3.56 (s, 3H), 3.52 – 3.46 (m, 2H), 3.37 (t, J = 6.4 Hz, 2H), 2.73 (br s, 2H), 2.24 – 2.11 (m, 4H), 1.84 (br d, J = 9.9 Hz, 2H), 1.54 – 1.41 (m, 6H), 1.35 – 1.25 (m, 2H), 1.01 – 0.94 (m, 2H).

**$^{13}$C NMR** (100 MHz, DMSO-D$_6$) δ 175.0, 161.1, 150.8, 150.6, 142.0, 138.6, 128.1, 122.1, 120.0, 118.7, 115.1, 113.6, 113.3, 74.8, 67.3, 55.8, 51.4, 36.8, 34.2, 31.7, 29.9, 25.9, 24.9.

**LCMS:** 524.5 [M+H$^+$].

* synthesis described in patent WO2021/55295, page 195.

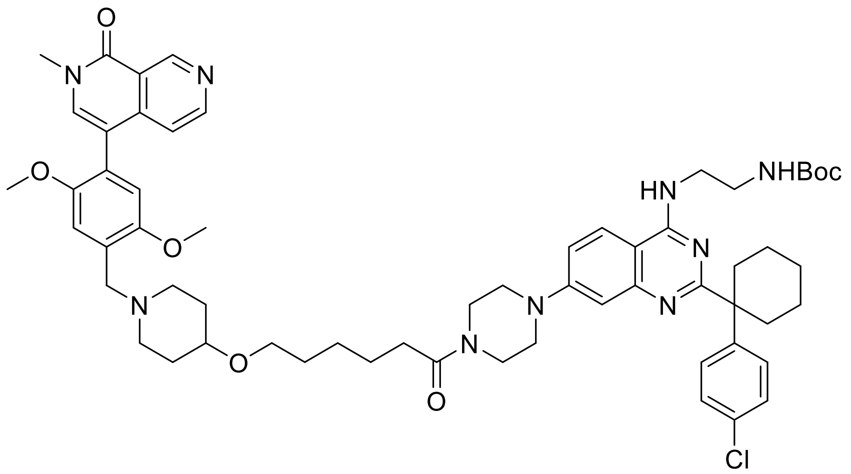


**tert-butyl (2-((2-(1-(4-chlorophenyl)cyclohexyl)-7-(4-(6-((1-(2,5-dimethoxy-4-(2-methyl-1-oxo-1,2-dihydro-2,7-naphthyridin-4-yl)benzyl)piperidin-4-yl)oxy)hexanoyl)piperazin-1-yl)quinazolin-4-yl)amino)ethyl)carbamate**

To a solution of 6-((1-(2,5-dimethoxy-4-(2-methyl-1-oxo-1,2-dihydro-2,7-naphthyridin-4-yl)benzyl)piperidin-4-yl)oxy)hexanoic acid (310 mg, 1.0 Eq, 0.59 mmol) and tert-butyl (2-((2-(1-(4-chlorophenyl)cyclohexyl)-7-(piperazin-1-yl)quinazolin-4-yl)amino)ethyl)carbamate (334 mg, 1.0 Eq, 0.59 mmol) in DMF (5 mL) was added EDCI (227 mg, 2.0 Eq, 1.18 mmol), HOBt (160 mg, 2.0 Eq, 1.18 mmol) and diisopropyl ethylamine (153 mg, 2.0 Eq, 1.18 mmol) and the reaction mixture was stirred at 25 °C for 2 hours. The mixture was diluted with water (20 mL), extracted with ethyl acetate (3 x 20 mL), the combined organic phases were washed with brine (2 x 20 mL) and dried over sodium sulfate. The residue was purified by preparative HPLC on a Phenomenex Luna C18 column (150 x 40 mm, 15 μm) eluting with acetonitrile (from 26 % to 56 %) in an aqueous solution of TFA (0.1 %). Fractions containing the title compound were combined and freeze dried yielding the title compound tert-butyl (2-((2-(1-(4-chlorophenyl)cyclohexyl)-7-(4-(6-((1-(2,5-dimethoxy-4-(2-methyl-1-oxo-1,2-dihydro-2,7-naphthyridin-4-yl)benzyl)piperidin-4-yl)oxy)hexanoyl)piperazin-1-yl)quinazolin-4-yl)amino)ethyl)carbamate as a white solid (400 mg, 0.37 mmol, 63 %).

**$^{1}$H NMR** (400 MHz, DMSO-D$_6$) δ 12.62 (br s, 1H), 9.76 (br s, 2H), 9.45 (d, J = 4.5 Hz, 1H), 8.69 – 8.65 (m, 1H), 8.15 (br d, J = 9.5 Hz, 1H), 7.83 (s, 1H), 7.48 (br d, J = 8.5 Hz, 2H), 7.42 – 7.39 (m, 2H), 7.34 (d, J = 12.5 Hz, 1H), 7.12 (s, 2H), 7.10 – 7.02 (m, 2H), 4.34 (br s, 2H), 3.83 (d, J = 2.1 Hz, 3H), 3.73 – 3.63 (m, 6H), 3.59 (br d, J = 1.1 Hz, 7H), 3.49 (br s, 3H), 3.46 – 3.38 (m, 5H), 3.20 – 2.97 (m, 3H), 2.80 – 2.69 (m, 2H), 2.43 – 2.28 (m, 3H), 2.18 – 2.03 (m, 3H), 2.03 – 1.77 (m, 3H), 1.63 (br s, 3H), 1.52 (td, J = 6.6, 13.3 Hz, 7H), 1.31 (s, 9H), 1.25 – 1.05 (m, 2H).

**$^{13}$C NMR** (100 MHz, DMSO-D$_6$) δ 160.9, 152.6, 151.2, 149.9, 149.2, 143.5, 142.6, 139.9, 132.1, 129.1, 128.8, 125.8, 120.1, 118.9, 116.7, 115.8, 112.8, 98.5, 78.3, 67.8, 56.8, 56.3, 53.8, 49.4, 37.0, 28.6.

**HR MS:** obs. m/z [M+H$^+$]: 1070.5660; calculated formular: $C_{60}H_{76}ClN_9O_7$; target mass error: -2.9 ppm.

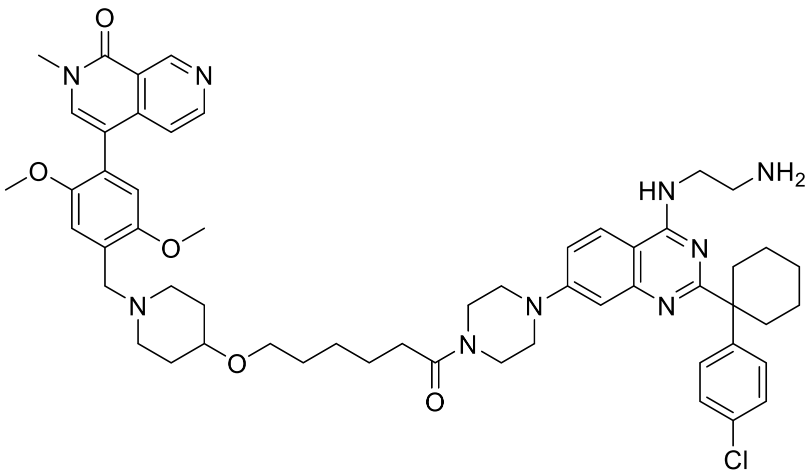


**4-(4-((4-((6-(4-(4-((2-aminoethyl)amino)-2-(1-(4-chlorophenyl)cyclohexyl)quinazolin-7-yl)piperazin-1-yl)-6-oxohexyl)oxy)piperidin-1-yl)methyl)-2,5-dimethoxyphenyl)-2-methyl-2,7-naphthyridin-1(2H)-one**

To a solution of tert-butyl (2-((2-(1-(4-chlorophenyl)cyclohexyl)-7-(4-(6-((1-(2,5-dimethoxy-4-(2-methyl-1-oxo-1,2-dihydro-2,7-naphthyridin-4-yl)benzyl)piperidin-4-yl)oxy)hexanoyl)piperazin-1-yl)quinazolin-4-yl)amino)ethyl)carbamate (300 mg, 1.0 Eq, 0.28 mmol) in 1,4-dioxane (1 mL) was added a solution of HCl in 1,4-dioxane (1.0 mL, 4 M) at 25 °C and the reaction mixture was stirred for 2 hours. A saturated solution of $NaHCO_3$ (10 mL) was added and the mixture was extracted with DCM (3 x 10 mL), the combined organic phases were washed with brine (10 mL) and dried over sodium sulfate, yielding the title compound 4-(4-((4-((6-(4-(4-((2-aminoethyl)amino)-2-(1-(4-chlorophenyl)cyclohexyl)quinazolin-7-yl)piperazin-1-yl)-6-oxohexyl)oxy)piperidin-1-yl)methyl)-2,5-dimethoxyphenyl)-2-methyl-2,7-naphthyridin-1(2H)-one as a white solid (207 mg, 0.21 mmol, 76 %).

**$^1$H NMR** (400 MHz, DMSO-$D_6$) δ 9.40 (s, 1H), 8.64 (d, J = 5.6 Hz, 1H), 8.02 – 7.92 (m, 1H), 7.86 – 7.77 (m, 1H), 7.73 (s, 1H), 7.49 – 7.36 (m, 2H), 7.28 – 7.22 (m, 2H), 7.20 (dd, J = 2.3, 9.0 Hz, 1H), 7.13 (s, 1H), 7.03 (d, J = 5.8 Hz, 1H), 6.91 (s, 1H), 6.88 – 6.82 (m, 1H), 3.74 (s, 4H), 3.65 – 3.62 (m, 3H), 3.60 (br s, 4H), 3.56 (s, 3H), 3.50 (br s, 2H), 3.44 – 3.36 (m, 4H), 3.29 (br s, 4H), 2.83 (br s, 2H), 2.78 – 2.65 (m, 4H), 2.36 (br t, J = 7.4 Hz, 2H), 2.15 (br t, J = 9.9 Hz, 2H), 1.99 – 1.76 (m, 5H), 1.59 – 1.41 (m, 12H), 1.39 – 1.23 (m, 4H).

**$^{13}$C NMR** (100 MHz, DMSO-$D_6$) δ 171.2, 161.1, 152.8, 152.1, 151.4, 150.8, 150.6, 148.0, 142.0, 138.6, 130.5, 128.7, 128.2, 128.1, 123.9, 122.1, 120.0, 118.7, 116.1, 115.2, 113.6, 113.3, 109.2, 106.1, 74.9, 67.4, 56.5, 56.2, 55.8, 51.4, 44.7, 41.1, 36.9, 32.7, 25.1.

**HR MS:** obs. m/z [M+H$^+$]: 970.5134; calculated formular: $C_{55}H_{68}ClN_9O_5$; target mass error: -2.9 ppm.

**Compound DBt-3 Synthetic Methods:**

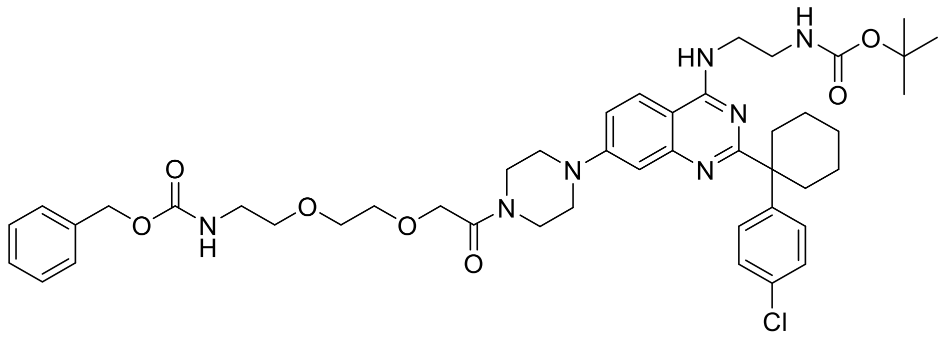

**benzyl (2-(2-(2-(4-(4-((2-((tert-butoxycarbonyl)amino)ethyl)amino)-2-(1-(4-chlorophenyl)cyclohexyl)quinazolin-7-yl)piperazin-1-yl)-2-oxoethoxy)ethoxy)ethyl)carbamate**

600mg 8-((Benzyloxycarbonyl-amino)-3,6-dioxaoctanoic acid dicyclohexylamine (Sigma Aldrich) was taken up in HCl 4N aqueous solution and extracted 4x with CH2Cl2. The organic layers were combined and dried over $MgSO_4$, filtered and concentrated in under reduced pressure yielding the free acid (0.42g, 1.353 mmol, 1.3 Eq.). The 8-((Benzyloxycarbonyl-amino)-3,6-dioxaoctanoic frees acid , HATU (530 mg, 1.353 mmol, 1.3 Eq.) and DIPEA (0.727 ml, 4.16 mmol, 4.0 Eq.) were solved 5mL DMA in $N_2$ atmosphere. The solution was stirred for 15min and then tert-butyl (2-((2-(1-(4-chlorophenyl)cyclohexyl)-7-(piperazin-1-yl)quinazolin-4-yl)amino)ethyl)carbamate (600 mg, 1.040 mmol, 1.0 Eq.) was added and stirred at room temperature for 2h yielding in a yellow to red solution. This reaction mix was diluted with ethyl acetate. The organic phase was washed with 30mL water/aq NaHCO3, 30mL water and 30mL brine. The organic layers were combined, dried with $Na_2SO_4$, filtered and volatiles were removed in vacuo to dryness. The resulting product was dissolved in DCM and adsorbed onto 4.7g SiO2 and was purified by Combi Flash chromatography (detection at 254/280 nM on a Combi Flash Rf 200 system with silica) with a gradient of DCM to DCM-MeOH 9-1. Fractions containing product were evaporated to produce a pink solid (0.699g, 0.828 mmol, 80%).

LC-MS: 844.2/846.4=M+H(+);

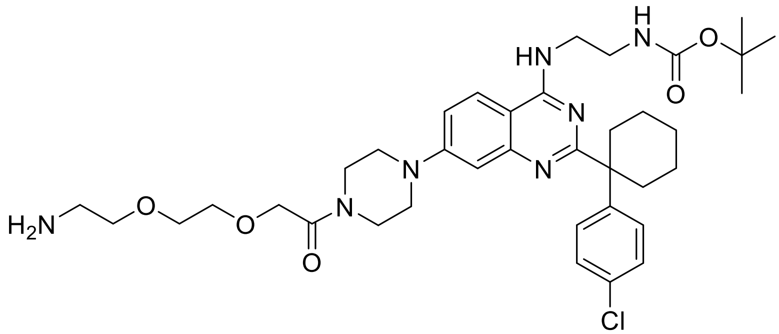

**tert-butyl (2-((7-(4-(2-(2-(2-aminoethoxy)ethoxy)acetyl)piperazin-1-yl)-2-(1-(4-chlorophenyl)cyclohexyl)quinazolin-4-yl)amino)ethyl)carbamate**

A solution of benzyl (2-(2-(2-(4-(4-((2-((tert-butoxycarbonyl)amino)ethyl)amino)-2-(1-(4-chlorophenyl)cyclohexyl)quinazolin-7-yl)piperazin-1-yl)-2-oxoethoxy)ethoxy)ethyl)carbamate (150 mg, 0.178 mmol) in Ethanol (Volume: 6 mL) was degassed. Afterwards Chlorobenzene (0.040 mL, 0.391 mmol, 1.0 Eq.), HCl 1.25M in EtOH (0.156 mL, 0.195 mmol, 1.1 Eq.) and Pd-C 10% (37.8 mg, 0.036 mmol, 0.2 Eq.) were added. Following a balloon of hydrogen gas was fitted. The reaction mixture was stirred at room temperature for 1.5h. The catalyst was filtered off and the filtrate was evaporated to dryness. Crude was purified by using preparative HPLC (Gylson GX-281 with a water-0.1%TFA / Acetonitrile gradient). To the fractions containing the product 86mg $NaHCO_3$ was added, the acetonitrile partially evaporated, and the product was extracted with ethyl acetate. The water-layer was re-extracted with 1x EtOAc-MeOH 9-1 and the combined-organic phases were dried over $Na_2SO_4$. After filtration the filtrate was evaporated under reduced pressure to yield white solid of tert-butyl (2-((7-(4-(2-(2-(2-aminoethoxy)ethoxy)acetyl)piperazin-1-yl)-2-(1-(4-chlorophenyl)cyclohexyl)quinazolin-4-yl)amino)ethyl)carbamate (96mg, 0.134 mmol, 75%).

LC-MS: 710/711=M+H(+);

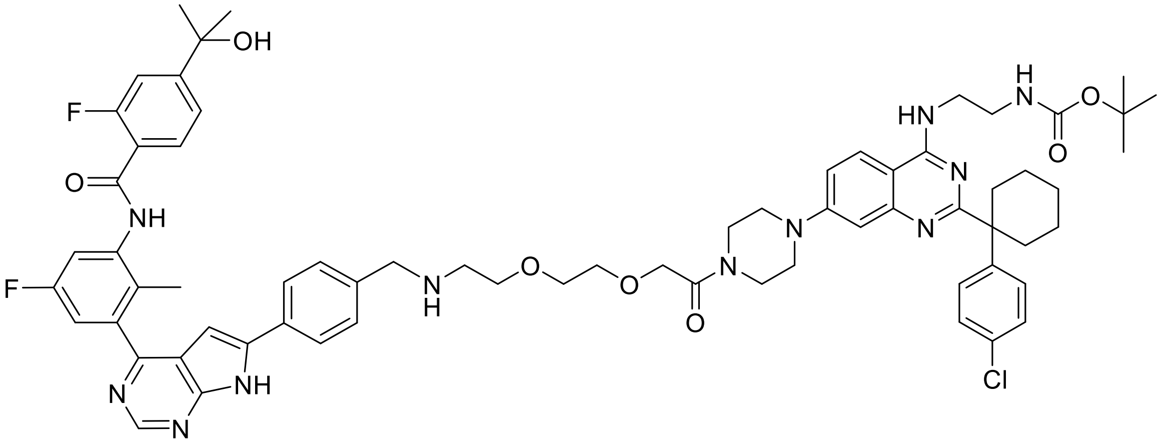


**tert-butyl (2-((2-(1-(4-chlorophenyl)cyclohexyl)-7-(4-(2-(2-(2-((4-(4-(5-fluoro-3-(2-fluoro-4-(2-hydroxypropan-2-yl)benzamido)-2-methylphenyl)-7H-pyrrolo[2,3-d]pyrimidin-6-yl)benzyl)amino)ethoxy)ethoxy)acetyl)piperazin-1-yl)quinazolin-4-yl)amino)ethyl)carbamate**

tert-butyl (2-((7-(4-(2-(2-(2-aminoethoxy)ethoxy)acetyl)piperazin-1-yl)-2-(1-(4-chlorophenyl)cyclohexyl)quinazolin-4-yl)amino)ethyl)carbamate (95 mg, 0.132 mmol, 1.0 Eq.) was suspended in 3mL MeOH and 3 mL THF, 2-fluoro-N-(5-fluoro-3-(6-(4-formylphenyl)-7H-pyrrolo[2,3-d]pyrimidin-4-yl)-2-methylphenyl)-4-(2-hydroxypropan-2-yl)benzamide* (69.7 mg, 0.132 mmol, 1.0 Eq.) and Sodium sulfate (94 mg, 0.662 mmol, 5.0 Eq.), acetic acid (7.58 µl, 0.132 mmol, 1.0 Eq.) were added at room temperature. The suspension was stirred at 23°C for 10 minutes. Sodium cyanoborohydride (42.5 mg, 0.662 mmol, 5.0 eq.) was added and the reaction mixture was stirred at room temperature overnight. To the resulting yellow-brown cloudy solution 1mL of water was added and left stirring for 10 minutes. The reaction mixture was partitioned between water and EtOAc, the layers were separated and extracted with 1x15ml EtOAc, washed with 3x10ml water. The organic layers were combined, dried with $Na_2SO_4$, filtered, and concentrated under reduced pressure. The crude product was dissolved in 3mL methanol and purified by using preparative HPLC (Gylson GX-281 with a water-0.1%TFA / Acetonitrile gradient). To the fractions containing the product 95mg $NaHCO_3$ was added, the acetonitrile was evaporated, and the product was extracted with ethyl acetate. The combined-organic phases were dried over $Na_2SO_4$, filtered and concentrated to dryness to produce the slightly yellow product (80mg, 0.066 mmol, 49.5 %).

* synthesis described in patent WO2021/55295, page 195.

LC-MS: 1218/1219 (M+H(-)

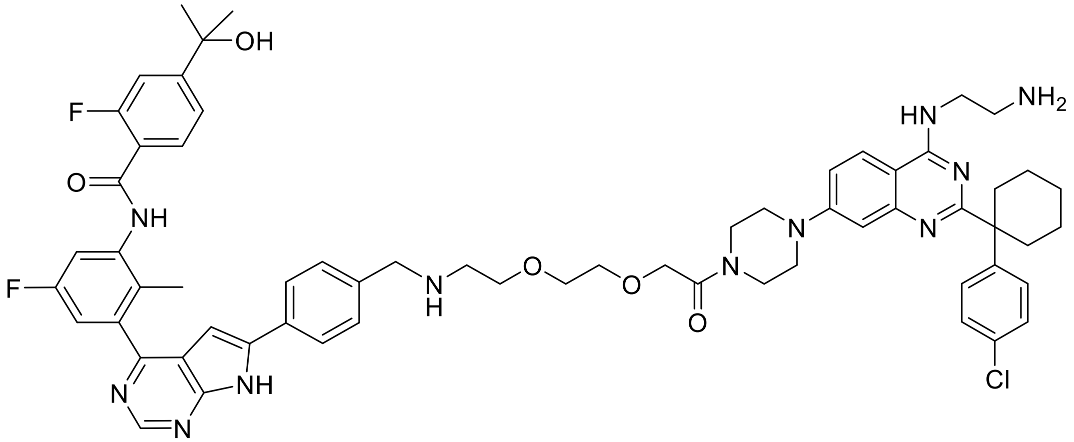


**DBt-3**

**N-(3-(6-(4-(((2-(2-(2-(4-(4-((2-aminoethyl)amino)-2-(1-(4-chlorophenyl)cyclohexyl)quinazolin-7-yl)piperazin-1-yl)-2-oxoethoxy)ethoxy)ethyl)amino)methyl)phenyl)-7H-pyrrolo[2,3-d]pyrimidin-4-yl)-5-fluoro-2-methylphenyl)-2-fluoro-4-(2-hydroxypropan-2-yl)benzamide**

A solution of tert-butyl (2-((2-(1-(4-chlorophenyl)cyclohexyl)-7-(4-(2-(2-(2-((4-(4-(5-fluoro-3-(2-fluoro-4-(2-hydroxypropan-2-yl)benzamido)-2-methylphenyl)-7H-pyrrolo[2,3-d]pyrimidin-6-yl)benzyl)amino)ethoxy)ethoxy)acetyl)piperazin-1-yl)quinazolin-4-yl)amino)ethyl)carbamate (78 mg, 0.064 mmol, 1.0 Eq.) in 4mL Dichloromethane was cooled in an ice bath before addition of TFA (0.148 mL, 1.917 mmol, 30.0 Eq.). The reaction mixture was stirred at room temperature for 2.5h resulting in a light yellow solution. Dichloromethane was evaporated while cooling the reaction mix in an ice bath followed by the addition of 3mL water/Acetonitrile (1:1). The reaction mix was lyophilized resulting in a slightly yellow powder of DBt-4 (98mg, 0.053 mmol, 84 %). The identity and purity of the final compound were confirmed both LC-High-resolution-MS and by 1H and 13C NMR.

**$^{1}$H NMR** (600 MHz, DMSO-D$_6$) δ ppm: 1.44 – 1.33 (m, 2H), 1.45 (s, 6H), 1.57 – 1.46 (m, 2H), 1.66 – 1.57 (m, 2H), 2.16 – 2.06 (m, 2H), 2.17 (s, 3H), 2.81 – 2.70 (m, 2H), 3.20 – 3.11 (m, 4H), 3.60 – 3.42 (m, 8H), 3.63 (s, 4H), 3.73 (t, J = 5.1 Hz, 2H), 3.97 – 3.88 (m, 2H), 4.27 – 4.20 (m, 4H), 6.95 (d, J = 2.0 Hz, 1H), 7.16 (s br, 1H), 7.23 (dd, J = 8.8, 2.8 Hz, 1H), 7.46 – 7.36 (m, 5H), 7.51 – 7.46 (m, 2H), 7.60 (d, J = 8.1 Hz, 2H),

7.67 (d, J = 10.2 Hz, 1H), 7.73 (t, J = 7.8 Hz, 1H), 8.15 – 8.02 (m, 6H), 8.86 (s, 1H), 9.08 (s br, 2H), 9.84 (s br, 1H), 9.96 (d, J = 2.5 Hz, 1H), 12.87 (s br, 2H).

**$^{13}$C NMR** (150 MHz, DMSO-D$_6$) δ ppm: 167.82, 165.74, 162.87, 159.42 (d, J = 242 Hz), 159.28, 159.11 (d, J = 248 Hz), 158.62, 156.82 (d, J = 7.2 Hz), 156.69 (d, J = 2.1 Hz), 154.29, 153.30, 150.85, 139.31 (d, J = 8.4 Hz), 139.23, 138.45 (d, J = 10.7 Hz), 132.29, 131.82, 131.10, 130.65, 129.89 (d, J = 2.9 Hz), 128.54, 128.41, 126.54, 126.52, 125.96, 125.37, 121.32 (d, J = 14.6 Hz), 120.77 (d, J = 2.9 Hz), 117.92, 116.24, 113.44 (d, J = 21.9 Hz), 112.61 (d, J = 23.2 Hz), 112.36 (d, J = 23.4 Hz), 97.13, 70.62 (d, J = 1.4 Hz), 69.78, 69.49, 68.76, 65.56, 49.76, 48.91, 46.02, 45.89, 45.72, 43.08, 40.42, 37.90, 34.03, 31.57, 24.86, 22.56, 14.58.

**HR MS:** obs. m/z [M+H$^+$]: calculated for $C_{62}H_{69}N_{11}O_5ClF_2$, 1120.51342; found, 1120.51247; deviation: 0.8 ppm.

**Compound DBt-4 Synthetic Methods:**

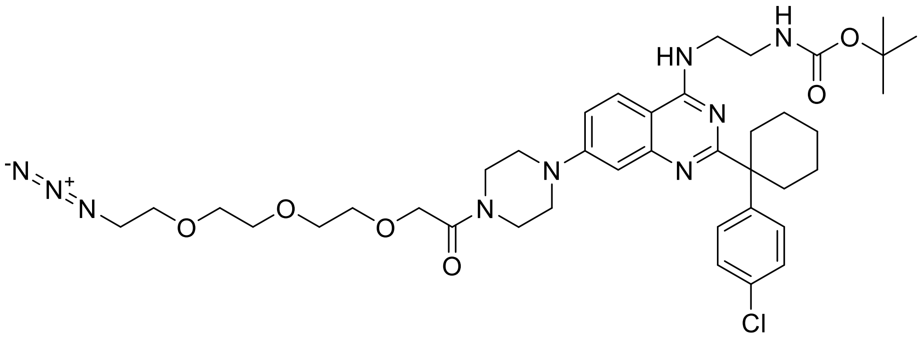

**tert-butyl (2-((7-(4-(2-(2-(2-(2-azidoethoxy)ethoxy)ethoxy)acetyl)piperazin-1-yl)-2-(1-(4-chlorophenyl)cyclohexyl)quinazolin-4-yl)amino)ethyl)carbamate**

To a solution of 11-Azido-3,6,9-trioxaundecanoic acid (842 mg, 1.0 Eq., 3.61 mmol) in DMA (Volume: 10 mL) DIPEA (1.576 mL, 2.0 Eq., 9.02 mmol) and HATU (1487 mg, 1.1 Eq., 3.91 mmol) were added. After stirring for 10 minutes at RT, tert-butyl (2-((2-(1-(4-chlorophenyl)cyclohexyl)-7-(piperazin-1-yl)quinazolin-4-yl)amino)ethyl)carbamate (1700 mg, 0.8 Eq., 3.01 mmol) was added and the suspension further stirred at RT. After 30min the suspension had almost dissolved. After 1.5h, reaction mixture turned into a clear solution; LC-MS analysis confirmed the completion of the reaction. Water:Brine (1:1 20 mL) was added, the mixture was extracted with ethyl acetate (2 x 20 mL), the combined organic phases were washed with brine (3 x 30mL) and evaporated under reduced pressure. The crude residue was adsorbed on Isolute and eluted of 24g silica gel column by a gradient of A= DCM and B= DCM:MeOH:NH4OH 90:9:1 with a gradient of 10% to 100% B in 20 min. Fractions containing product were evaporated to produce a colorless oil (2.54g, quant.).

LC-MS: m/z 780 [M+H]+

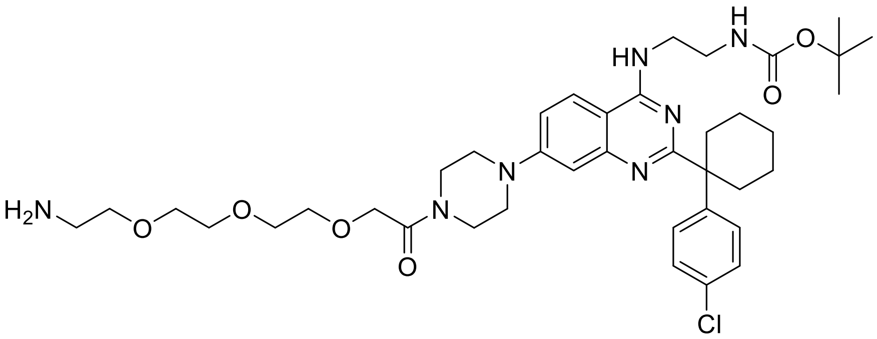

**tert-butyl (2-((7-(4-(2-(2-(2-(2-aminoethoxy)ethoxy)ethoxy)acetyl)piperazin-1-yl)-2-(1-(4-chlorophenyl)cyclohexyl)quinazolin-4-yl)amino)ethyl)carbamate**

To a solution of tert-butyl (2-((7-(4-(2-(2-(2-(2-azidoethoxy)ethoxy)ethoxy)acetyl)piperazin-1-yl)-2-(1-(4-chlorophenyl)cyclohexyl)quinazolin-4-yl)amino)ethyl)carbamate (1.42 g, 1.0 Eq., 1.820 mmol) in THF (Volume: 20 mL) Triphenylphosphine (0.525 g, 1.1 Eq., 2.002 mmol) and Water (0.049 mL, 1.5 Eq., 2.73

mmol) were added. The round bottom flask was put under Argon and the colorless solution was stirred for 3 days at RT. The reaction mixture was evaporated to dryness (colorless oil) and dissolved into DCM, adsorbed on Isolute and eluted of a 40g silica gel column with gradient of A=DCM and B=DCM:MeOH:NH4OH 80:18:2. Fractions containing pure product were pooled and evaporated to dryness to give a white solid foam as the desired product (778mg, 1.032 mmol, 57%).

LC-MS: m/z 754.5 [M+H]+

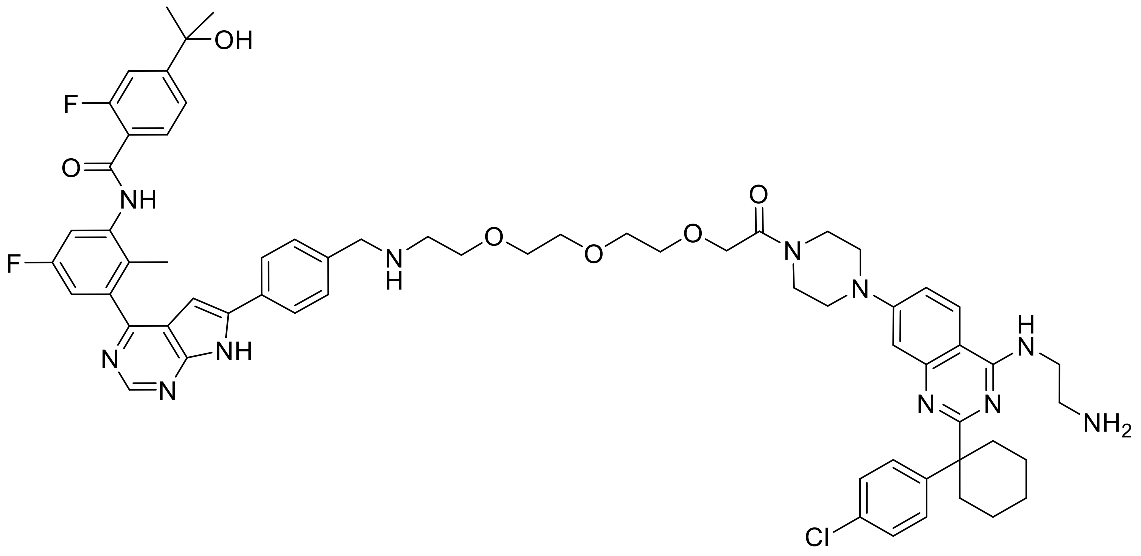


**DBt-4**

**N-(3-(6-(4-(13-(4-(4-((2-aminoethyl)amino)-2-(1-(4-chlorophenyl)cyclohexyl)quinazolin-7-yl)piperazin-1-yl)-13-oxo-5,8,11-trioxa-2-azatridecyl)phenyl)-7H-pyrrolo[2,3-d]pyrimidin-4-yl)-5-fluoro-2-methylphenyl)-2-fluoro-4-(2-hydroxypropan-2-yl)benzamide**

To a solution of tert-butyl (2-((7-(4-(2-(2-(2-(2-aminoethoxy)ethoxy)ethoxy)acetyl)piperazin-1-yl)-2-(1-(4-chlorophenyl)cyclohexyl)quinazolin-4-yl)amino)ethyl)carbamate (375 mg, 1.0 Eq., 0.498 mmol) in MeOH (Volume: 12 mL, Ratio: 1.000) and THF (Volume: 12 mL, Ratio: 1.000) were added 2-fluoro-N-(5-fluoro-3-(6-(4-formylphenyl)-7H-pyrrolo[2,3-d]pyrimidin-4-yl)-2-methylphenyl)-4-(2-hydroxypropan-2-yl)benzamide* (262 mg, 1.0 Eq., 0.498 mmol), Sodium sulfate (707 mg, 10 Eq., 4.98 mmol) and acetic acid (0.028 mL, 1.0 Eq., 0.498 mmol). The suspension was stirred under Argon for 10min at RT, then $NaCNBH_3$ (329 mg, 10 Eq., 4.98 mmol) was added and the mixture was stirred overnight (for 16h) at RT. The reaction mixture was poured into 0.1 M HCl (100mL, 20.1 Eq., 10.00 mmol) and stirred at RT for 45min. It was then neutralized with 10mL saturated, aqueous $NaHCO_3$ solution and extracted twice with DCM, dried on phase separator and evaporated to yield a yellow solid (770 mg). This residue was dissolved in a DCM/methanol mixture (1:1) and eluted of a 40g silica gel column with gradient of A=DCM, B=DCM/MeOH/NH4OH 80:18:2 (10% to 100% B). Fractions containing the product were pooled and the solvent was evaporated. This residue was used directly for BOC deprotection where the resulting tert-butyl (2-((2-(1-(4-chlorophenyl)cyclohexyl)-7-(4-(1-(4-(4-(5-fluoro-3-(2-fluoro-4-(2-hydroxypropan-2-yl)benzamido)-2-

methylphenyl)-7H-pyrrolo[2,3-d]pyrimidin-6-yl)phenyl)-5,8,11-trioxa-2-azatridecan-13-oyl)piperazin-1-yl)quinazolin-4-yl)amino)ethyl)carbamate (250mg, 1.0 Eq., 0.150 mmol) was dissolved in dichloromethane (2mL) and TFA (0.579 mL, 50 Eq., 7.51 mmol) and stirred at RT for 25 minutes. The reaction was diluted with dichloromethane and washed with a aqueous, saturated solution of $NaHCO_3$. The resulting aqueous phase was extracted with dichloromethane with additional % methanol and the combined organic phases were washed with brine (2 x 10 mL) and dried over sodium sulfate. The organic phase was further dried on phase separator and evaporated to yield a yellow solid (285 mg). This solid was dissolved in methanol (3mL) with approx. 10% NMP and was purified by preparative HPLC on a Waters SUNFIRE preparative C18 column (OBD 5µm. 30 X 100mm) eluting with acetonitrile (from 5 % to 70 %) in an aqueous solution of TFA (0.1 %). Fractions containing the title compound were combined and freeze dried yielding the title compound N-(3-(6-(4-(13-(4-(4-((2-aminoethyl)amino)-2-(1-(4-chlorophenyl)cyclohexyl)quinazolin-7-yl)piperazin-1-yl)-13-oxo-5,8,11-trioxa-2-azatridecyl)phenyl)-7H-pyrrolo[2,3-d]pyrimidin-4-yl)-5-fluoro-2-methylphenyl)-2-fluoro-4-(2-hydroxypropan-2-yl)benzamide as light yellow solid (84mg, 0.072 mmol, 37%). The identity and purity of the final compound were confirmed both by high resolution MS as well as 1H and 13C NMR.

* synthesis described in patent WO2021/55295, page 195.

**$^1$H NMR** (600 MHz, DMSO-$D_6$) δ ppm: 1.44 – 1.35 (m, 2H), 1.45 (s, 6H), 1.57 – 1.46 (m, 2H), 1.66 – 1.58 (m, 2H), 2.16 – 2.07 (m, 2H), 2.18 (s, 3H), 2.84 – 2.72 (m, 2H), 3.22 – 3.08 (m, 4H), 3.63 – 3.40 (m, 16H), 3.71 (t, J = 5.2 Hz, 2H), 3.98 – 3.89 (m, 2H), 4.19 (s, 2H), 4.25 – 4.21 (m, 2H), 6.96 (d, J = 2.1 Hz, 1H), 7.27 – 7.18 (m, 2H), 7.46 – 7.37 (m, 5H), 7.53 – 7.47 (m, 2H), 7.60 (d, J = 8.1 Hz, 2H), 7.67 (d, J = 10.2 Hz, 1H), 7.73 (t, J = 7.8 Hz, 1H), 8.07 (d, J = 8.3 Hz, 2H), 8.12 (s br, 2H), 8.18 (d, J = 9.4 Hz, 1H), 8.88 (s, 1H), 9.10 (s br, 2H), 9.90 (s br, 1H), 9.96 (d, J = 2.5 Hz, 1H), 12.88 (s, 1H), 12.91 (s br, 1H).

**$^{13}$C NMR** (150 MHz, DMSO-$D_6$) δ ppm: 171.61, 167.68, 165.70, 162.87, 159.42 (d, J = 241.7 Hz), 159.26, 159.11 (d, J = 248.2 Hz), 158.28, 156.81 (d, J = 6.9 Hz), 156.68, 154.31, 153.31, 150.86, 142.84, 140.90,f 139.32, 139.26, 138.44 (d, J = 10.8 Hz), 132.30, 131.81, 131.09, 130.69, 129.90 (d, J = 2.9 Hz), 128.54, 128.42, 126.52 (d, J = 3.2 Hz), 125.96, 125.47, 121.32 (d, J = 14.6 Hz), 120.77 (d, J = 2.8 Hz), 117.94, 116.26, 113.44 (d, J = 21.9 Hz), 112.60 (d, J = 24.1 Hz), 112.35 (d, J = 23.3 Hz), 97.15, 70.62, 69.78 (d, J = 10.3 Hz), 69.51, 69.05, 65.64, 49.73, 48.92, 46.06, 45.81, 43.19, 40.39, 37.90, 34.06, 31.56, 24.86, 22.57, 14.59.

**HR MS:** obs. m/z [$M+H^+$]: calculated for $C_{64}H_{73}N_{11}O_6ClF_2$, 1164.53964; found, 1164.53857; deviation: 0.9 ppm.

**Compound DBt-5 Synthetic Methods:**

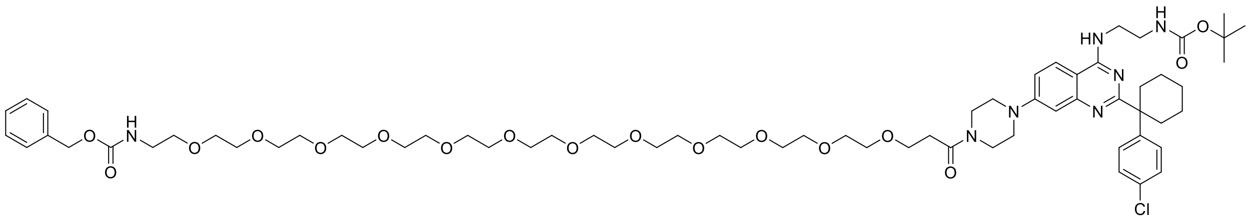


**tert-butyl (2-((2-(1-(4-chlorophenyl)cyclohexyl)-7-(4-(3-oxo-1-phenyl-2,7,10,13,16,19,22,25,28,31,34,37,40-tridecaoxa-4-azatritetracontan-43-oyl)piperazin-1-yl)quinazolin-4-yl)amino)ethyl)carbamate**

To a solution of CBZ-NH-PEG(12)-CH2-CH2-COOH (, 80 mg, 1.0 Eq., 0.105 mmol, CAS: 1334177-88-6, Sigma Aldrich) in DMA (Volume: 1 ml) DIPEA (0.055 ml, 3.0 Eq., 0.315 mmol) and HATU (52.5 mg, 1.3 Eq., 0.137 mmol) was added at room temperature (RT). The yellow suspension was stirred at RT and it was monitored by LC-MS. After 10min activation tert-butyl (2-((2-(1-(4-chlorophenyl)cyclohexyl)-7-(piperazin-1-yl)quinazolin-4-yl)amino)ethyl)carbamate (60 mg, 1.0 Eq., 0.105 mmol) was added at RT and the yellow solution was stirred at RT and it was monitored by LC-MS. After 1h The reaction mixture was partitioned between water and EtOAc, the layers were separated and extracted with 1x15ml EtOAc, washed with 3x10ml water The organic layers were combined, dried with $Na_2SO_4$, filtered and volatiles were removed in vacuo to dryness resulting in brown oil. This crude product was purified by using NP flash chromatography with a Isco Combiflash: 4g Silicagel NP Redisep column, eluent EtOAc/ EtOAc-MeOH/NH3 (9:1) 90-10, detection at 280 nm. monitoring 254 nm, Flow: 20ml/min. Fractions containing the product were combined and evaporated resulting in a pale-yellow oil (130mg, 0.100 mmol, 95%)

HPLC-MS: [MH]+ 1298.8+1300.8 + [M+Na/2]+

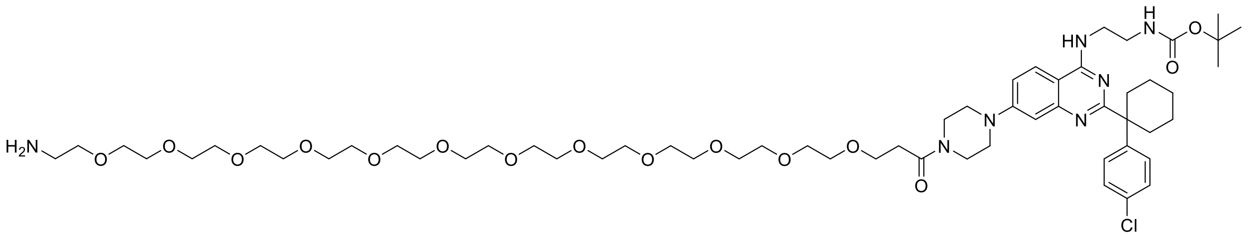


**tert-butyl (2-((7-(4-(1-amino-3,6,9,12,15,18,21,24,27,30,33,36-dodecaoxanonatriacontan-39-oyl)piperazin-1-yl)-2-(1-(4-chlorophenyl)cyclohexyl)quinazolin-4-yl)amino)ethyl)carbamate**

A solution of tert-butyl (2-((2-(1-(4-chlorophenyl)cyclohexyl)-7-(4-(3-oxo-1-phenyl-2,7,10,13,16,19,22,25,28,31,34,37,40-tridecaoxa-4-azatritetracontan-43-oyl)piperazin-1-yl)quinazolin-4-yl)amino)ethyl)carbamate (110 mg, 1.0 Eq., 0.080 mmol) in Ethanol (Volume: 5 mL) was degassed 3x with argon. Afterwards Chlorobenzene (0.016 mL, 2.0 Eq., 0.161 mmol), HCl 1.25M in EtOH (0.071 mL, 1.1 Eq.,

0.088 mmol) and Pd-C 10% (17.12 mg, 0.2 Eq., 0.016 mmol) were added. Following a ballon of hydrogen was fitted. The reaction mixture was stirred at room temperature and reaction was monitored by LC-MS. After 1.5h the catalyst was removed by filtration and the filtrate was evaporated to dryness. The residue was diluted with ethyl acetate and this the organic phase was washed with brine (50 mL), dried over sodium sulfate and concentrated under vacuum.

The residue was purified by preparative HPLC on a Waters SUNFIRE preparative C18 column (OBD 5µm. 30 X 100mm) eluting with acetonitrile TFA (from 5 % to 70 %) in an aqueous solution of TFA (0.1 %). Fractions containing the title compound were combined and the pH was adjusted to 7 using $NaHCO_3$ . After the evaporation of acetonitrile the water phase was partitioned between water and ethyl acetate. The layers were separated and extracted with 2x15ml ethyl acetate, washed with 1x10ml water. The organic layers were combined, dried with $Na_2SO_4$, filtered and volatiles were removed in vacuo to dryness resulting in a colorless resin (55mg, 0.047 mmol, 59%).

* synthesis described in patent WO2021/55295, page 195.

HPLC-MS: [MH]+ 1164.6 m/z

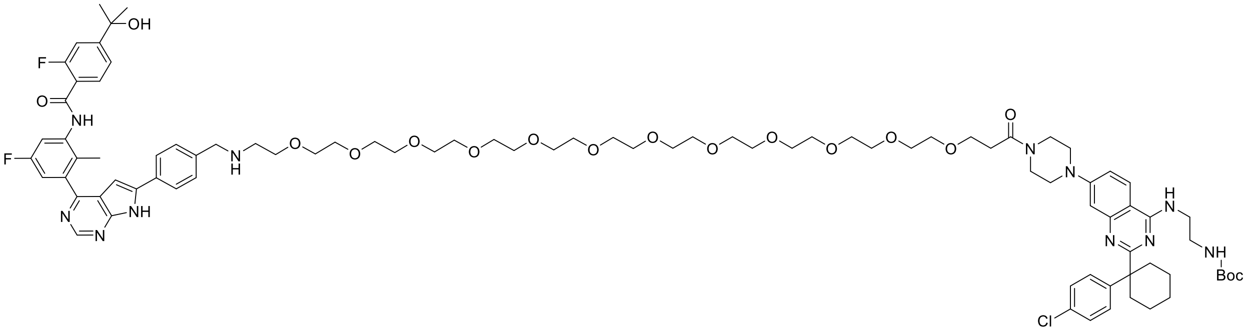


**tert-butyl (2-((2-(1-(4-chlorophenyl)cyclohexyl)-7-(4-(1-(4-(4-(5-fluoro-3-(2-fluoro-4-(2-hydroxypropan-2-yl)benzamido)-2-methylphenyl)-7H-pyrrolo[2,3-d]pyrimidin-6-yl)phenyl)-5,8,11,14,17,20,23,26,29,32,35,38-dodecaoxa-2-azahentetracontan-41-oyl)piperazin-1-yl)quinazolin-4-yl)amino)ethyl)carbamate**

2-fluoro-N-(5-fluoro-3-(6-(4-formylphenyl)-7H-pyrrolo[2,3-d]pyrimidin-4-yl)-2-methylphenyl)-4-(2-hydroxypropan-2-yl)benzamide* (11.63 mg, 1.0 Eq., 0.022 mmol) was suspended in CH3OH-THF 1:1 (Volume: 1 ml, Ratio: 2.500), tert-butyl (2-((7-(4-(1-amino-3,6,9,12,15,18,21,24,27,30,33,36-dodecaoxanonatriacontan-39-oyl)piperazin-1-yl)-2-(1-(4-chlorophenyl)cyclohexyl)quinazolin-4-yl)amino)ethyl)carbamate (26 mg, 1.0 Eq., 0.022 mmol) and 34mg $Na_2SO_4$ , acetic acid (1.265 µl, 1.0 Eq., 0.022 mmol) were added at RT. The yellow suspension was stirred at RT for 5min. $NaCNBH_4$ (7.3mg, 2.5 Eq., 0.055 mmol) was rapidly added at RT and the mixture was stirred at RT resulting in a yellow suspension. After 60 minutes additional NaCNBH4 (7.5 mg, 0.101 mmol) was added and under the addition of 0.5 mL TFA the suspension turned into a yellow solution. After 20h stirring the reaction was stopped by adding 50µl water, stirred for further 10min at RT, filtered, washed with MeOH and evaporated in vacuo to dryness resulting in yellow oil (70mg). This crude residue was purified by preparative HPLC on a Waters SUNFIRE preparative C18 column (OBD 5µm. 30 X 100mm) eluting with acetonitrile TFA (from 5 % to 70 %) in an aqueous solution of TFA (0.1 %). The tert-butyl (2-((2-(1-(4-chlorophenyl)cyclohexyl)-7-(4-(1-(4-(4-(5-fluoro-3-(2-fluoro-4-(2-hydroxypropan-2-yl)benzamido)-2-methylphenyl)-7H-pyrrolo[2,3-d]pyrimidin-6-yl)phenyl)-5,8,11,14,17,20,23,26,29,32,35,38-dodecaoxa-2-azahentetracontan-41-oyl)piperazin-1-yl)quinazolin-4-yl)amino)ethyl)carbamate as a white solid (12 mg, 0.007 mmol, 33 %).

HPLC-MS: [(M+2H)/2]+ 838.2 m/z

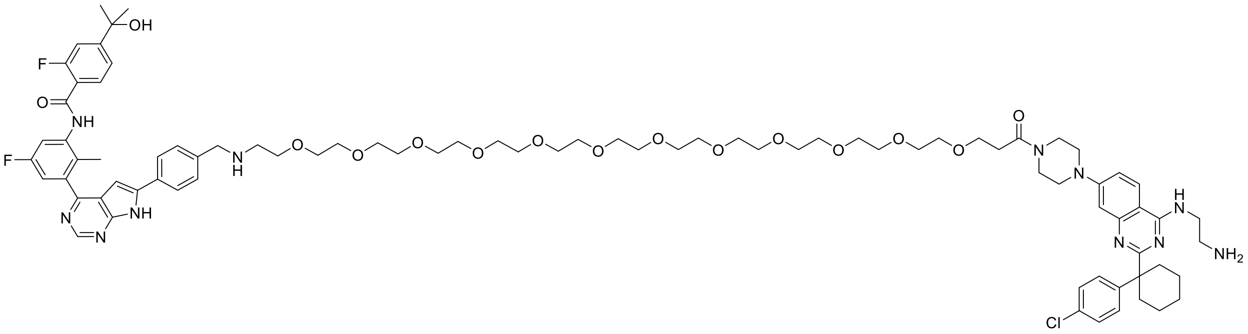


**DBt-5**

**N-(3-(6-(4-(41-(4-(4-((2-aminoethyl)amino)-2-(1-(4-chlorophenyl)cyclohexyl)quinazolin-7-yl)piperazin-1-yl)-41-oxo-5,8,11,14,17,20,23,26,29,32,35,38-dodecaoxa-2-azahentetracontyl)phenyl)-7H-pyrrolo[2,3-d]pyrimidin-4-yl)-5-fluoro-2-methylphenyl)-2-fluoro-4-(2-hydroxypropan-2-yl)benzamide**

To a solution of tert-butyl (2-((2-(1-(4-chlorophenyl)cyclohexyl)-7-(4-(1-(4-(4-(5-fluoro-3-(2-fluoro-4-(2-hydroxypropan-2-yl)benzamido)-2-methylphenyl)-7H-pyrrolo[2,3-d]pyrimidin-6-yl)phenyl)-5,8,11,14,17,20,23,26,29,32,35,38-dodecaoxa-2-azahentetracontan-41-oyl)piperazin-1-yl)quinazolin-4-yl)amino)ethyl)carbamate (11 mg, 1.0 Eq., 5.11 µmol) in $CH_2Cl_2$ (Volume: 50 µL) TFA (0.075 ml, 190 Eq., 0.968 mmol) was added at 0-5°C. The reaction was stirred at further at RT. Reaction monitoring by LC-MS and after 90 minutes was worked up by addition of Water 10 mL), the mixture was extracted with ethyl acetate (3 x 10 mL), the combined organic phases were washed with brine (2 x 10 mL) and dried over sodium sulfate. The yellow reaction mixture was evaporated at 0-5°C and the crude product was dissolved in 0.5ml methanol. This crude residue was purified by preparative HPLC on a Waters SUNFIRE preparative C18 column (OBD 5µm. 30 X 100mm) eluting with acetonitrile TFA (from 5 % to 70 %) in an aqueous solution of TFA (0.1 %). Fractions containing the final compound were combined and freeze-dried resulting in yellow powder (6mg, 3.81 µmol, 75%). The identity and purity of the final compound were confirmed both by high resolution MS as well as 1H and 13C NMR:

**$^1$H NMR** (600 MHz, DMSO-$D_6$) δ ppm: 1.43 – 1.32 (m, 1H), 1.45 (s, 6H), 1.45-1.48 (m, 2H), 1.57 – 1.50 (m, 1H), 1.67 – 1.58 (m, 2H), 2.16 – 2.04 (m, 2H), 2.18 (s, 3H), 2.63 (t, J = 6.7 Hz, 2H), 2.81 – 2.71 (m, 2H), 3.20 – 3.07 (m, 4H), 3.55 – 3.41 (m, 44H), 3.60 – 3.55 (m, 4H), 3.62 – 3.60 (m, 2H), 3.67 – 3.62 (m, 4H), 3.70 (t, J = 5.2 Hz, 2H), 4.00 – 3.85 (m, 2H), 4.22 (t, J = 5.6 Hz, 2H), 6.96 (d, J = 2.0 Hz, 1H), 7.13 (s, 1H), 7.24 (dd, J = 8.8, 2.8 Hz, 1H), 7.46 – 7.37 (m, 5H), 7.48 (d, J = 8.8 Hz, 2H), 7.60 (d, J = 8.4 Hz, 2H), 7.67 (d, J

= 10.1 Hz, 1H), 7.73 (t, J = 7.8 Hz, 1H), 8.10 – 7.99 (m, 5H), 8.13 (d, J = 9.1 Hz, 1H), 8.88 (s, 1H), 9.10 – 8.92 (m, 2H), 9.82 (s br, 1H), 9.96 (d, J = 2.5 Hz, 1H), 12.78 (s br, 1H), 12.87 (s, 1H).

**$^{13}$C NMR** (150 MHz, DMSO-D$_6$) δ ppm: 169.15, 162.86, 159.42 (d, J = 250 Hz), 159.28, 159.10 (d, J = 248 Hz), 158.24, 158.03, 156.83, 156.78, 154.36, 153.31, 150.97, 142.84, 139.42 (d, J = 8.6 Hz), 139.17, 138.43 (d, J = 10.5 Hz), 132.24, 131.14, 130.68, 129.89 (d, J = 2.9 Hz), 128.55, 128.37, 126.48 (d, J = 3.1 Hz), 125.96, 125.39, 121.33 (d, J = 14.6 Hz), 120.79 (d, J = 2.9 Hz), 117.94, 116.29, 113.41 (d, J = 22.1 Hz), 112.54 (d, J = 24.5 Hz), 112.35 (d, J = 23.3 Hz), 97.92, 97.12, 70.60 (d, J = 1.4 Hz), 69.73, 69.67, 69.56, 66.70, 65.62, 49.74, 48.93, 46.12, 45.83, 44.00, 37.92, 34.08, 32.80, 31.56, 24.86, 22.57, 14.57.

**HR MS:** obs. m/z [M+H$^+$]: calculated for $C_{83}H_{111}N_{11}O_{15}ClF_2$, 1574.79112; found, 1574.79100; deviation: 0.1 ppm.

**Compound DBt-6 Synthetic Methods:**

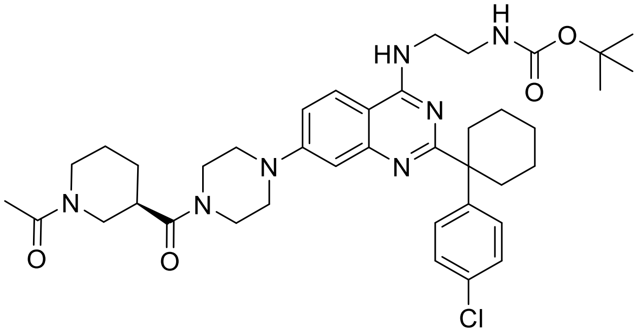

**tert-butyl (R)-(2-((7-(4-(1-acetylpiperidine-3-carbonyl)piperazin-1-yl)-2-(1-(4-chlorophenyl)cyclohexyl)quinazolin-4-yl)amino)ethyl)carbamate**

Under nitrogen atmosphere tert-butyl (2-((2-(1-(4-chlorophenyl)cyclohexyl)-7-(piperazin-1-yl)quinazolin-4-yl)amino)ethyl)carbamate (300 mg, 0.504 mmol, 1.0 Eq.), HATU (257 mg, 0.656 mmol, 1.3 Eq.) (3R)1-Acetylpiperidine-3-carboxylic acid (107 mg, 0.605 mmol, 1.2 Eq., CAS: 712270-39-8, PharmaBlockScience) , and DIPEA (0.264 ml, 1.513 mmol, 3.0 Eq.) were suspended in 2.3mL DMA. The suspension was stirred at room temperature for 2.5h resulting in a yellow-brown solution. The reaction mixture was partitioned between water and 100ml ethyl acetate. The phases were separated, and the organic phases were washed with 1x30ml water and 1x30ml brine. The organic layers were combined, dried with $Na_2SO_4$, filtered and volatiles were removed in vacuo to dryness yielding a yellow resin of the crude product. This crude product was dissolved in Dichloromethane and adsorbed onto 1.6g Isolute and was purified by Combi Flash chromatography (detection at 254/280nm on a Combi Flash Rf 200 system with silica) with a gradient of DCM to DCM-MeOH 9-1. Fractions containing product were evaporated to produce a white solid product (324 mg, 0.442 mmol, 88 %).

LC-MS: 718.4/720.4=M+H(+)

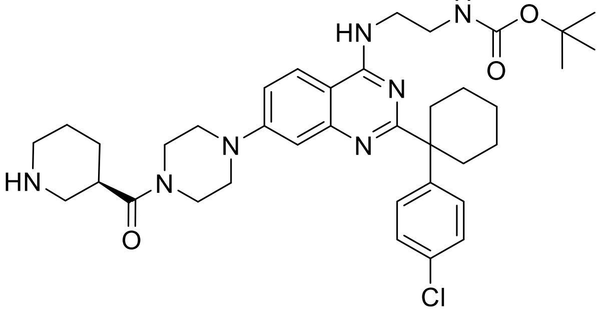

**tert-butyl (R)-(2-((2-(1-(4-chlorophenyl)cyclohexyl)-7-(4-(piperidine-3-carbonyl)piperazin-1-yl)quinazolin-4-yl)amino)ethyl)carbamate**

To a solution of tert-butyl (R)-(2-((7-(4-(1-acetylpiperidine-3-carbonyl)piperazin-1-yl)-2-(1-(4-chlorophenyl)cyclohexyl)quinazolin-4-yl)amino)ethyl)carbamate (324 mg, 0.451 mmol, 1.0 Eq.) in EtOH (Volume: 7 ml) was added KOH 2M (2.255 ml, 4.51 mmol, 10.0 Eq.). The reaction mixture was stirred at 80°C for 23h and then left at room temperature for 60h. The reaction mixture was evaporated and the residue was partitioned between water and ethyl acetate. The phases were separated, and the organic phases were washed with 1x30ml water and 1x30ml brine. The organic phase was dried with $Na_2SO_4$, filtered and filtrate was evaporated in vacuo to dryness producing a light beige foam of crude product. HPLC-MS analysis indicated that next to the desired tert-butyl (R)-(2-((2-(1-(4-chlorophenyl)cyclohexyl)-7-(4-(piperidine-3-carbonyl)piperazin-1-yl)quinazolin-4-yl)amino)ethyl)carbamate also tert-butyl (2-((2-(1-(4-chlorophenyl)cyclohexyl)-7-(piperazin-1-yl)quinazolin-4-yl)amino)ethyl)carbamate was obtained as side product.

LCMS : M+H(+) =676.4/678.4 m/z (66%) for tert-butyl (R)-(2-((2-(1-(4-chlorophenyl)cyclohexyl)-7-(4-(piperidine-3-carbonyl)piperazin-1-yl)quinazolin-4-yl)amino)ethyl)carbamate

M+H(+) = 565.3/567.3 m/z (32%) for tert-butyl (2-((2-(1-(4-chlorophenyl)cyclohexyl)-7-(piperazin-1-yl)quinazolin-4-yl)amino)ethyl)carbamate

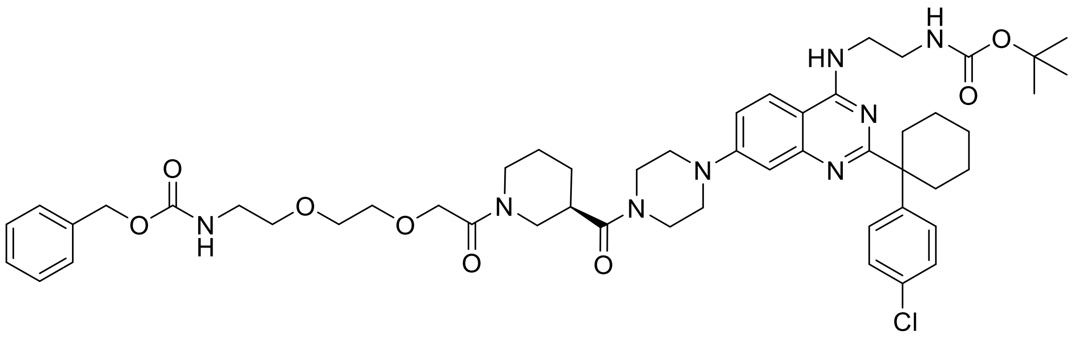


**tert-butyl (R)-(2-((2-(1-(4-chlorophenyl)cyclohexyl)-7-(4-(1-(3-oxo-1-phenyl-2,7,10-trioxa-4-azadodecan-12-oyl)piperidine-3-carbonyl)piperazin-1-yl)quinazolin-4-yl)amino)ethyl)carbamate**

8-((Benzyloxycarbonyl-amino)-3,6-dioxaoctanoic acid dicyclohexylamine (343mg, 0.673 mmol, 1.3Eq.) was taken up in 20mL HCl 4N aqueous solution and extracted three times with dichloromethane. The organic layers were combined, dried over $MgSO_4$, filtered and concentrated under reduced pressure resulting in white, free 8-((Benzyloxycarbonyl-amino)-3,6-dioxaoctanoic acid. Under nitrogen atmosphere free 8-((Benzyloxycarbonyl-amino)-3,6-dioxaoctanoic acid , HATU (264 mg, 0.673 mmol, 1.3 Eq.) and DIPEA (0.362 ml, 2.070 mmol, 4 Eq.) were solved 1.5mL DMA. The solution was stirred for 10min at room temperature and then the crude tert-butyl (R)-(2-((2-(1-(4-chlorophenyl)cyclohexyl)-7-(4-(piperidine-3-carbonyl)piperazin-1-yl)quinazolin-4-yl)amino)ethyl)carbamate product (350 mg, 0.518 mmol, 1.0 Eq.) (in 2ml DMA dissolved ) were added and stirred at room temperature for 30 minutes yielding a yellow-suspension. The reaction mixture was diluted with ethyl acetate and the organic phase was washed with a aqueous $NaHCO_3$ solution, water, and brine. The organic phases were combined, dried with $Na_2SO_4$, filtered and filtrate was removed in vacuo to dryness to produce a yellow-brown solid. This crude product was dissolved in Dichloromethane and adsorbed onto 1.8g Isolute and was purified by Combi Flash chromatography (detection at 254/280nm on a Combi Flash Rf 200 system with silica) with a gradient of ethyl acetate to methanol (100-0/90-10). In this step the side product of benzyl (2-(2-(2-(4-(4-((2-((tert-butoxycarbonyl)amino)ethyl)amino)-2-(1-(4-chlorophenyl)cyclohexyl)quinazolin-7-yl)piperazin-1-yl)-2-oxoethoxy)ethoxy)ethyl)carbamate was successfully separated. Fractions containing product were evaporated to produce a white solid product (245 mg, 0.247 mmol, 47.6 %).

LC-MS: M+H(+)=955.5/956.5 m/z

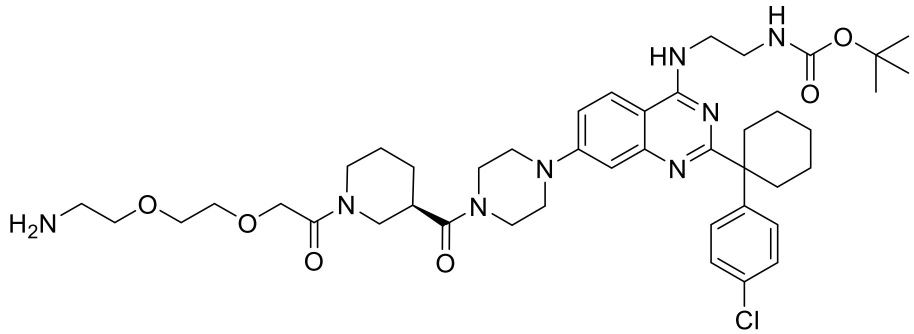

**tert-butyl (R)-(2-((7-(4-(1-(2-(2-(2-aminoethoxy)ethoxy)acetyl)piperidine-3-carbonyl)piperazin-1-yl)-2-(1-(4-chlorophenyl)cyclohexyl)quinazolin-4-yl)amino)ethyl)carbamate**

A solution of tert-butyl (R)-(2-((2-(1-(4-chlorophenyl)cyclohexyl)-7-(4-(1-(3-oxo-1-phenyl-2,7,10-trioxa-4-azadodecan-12-oyl)piperidine-3-carbonyl)piperazin-1-yl)quinazolin-4-yl)amino)ethyl)carbamate (245 mg, 0.256 mmol, 1.0 Eq.) in Ethanol (Volume: 8 mL) was degassed. Afterwards Chlorobenzene (0.057 mL, 0.564 mmol, 2.2 Eq.), HCl 1.25M in EtOH (0.226 mL, 0.282 mmol, 1.1 Eq.) and Pd-C 10% (54.6 mg, 0.051 mmol, 0.2 Eq.) were added. Following a balloon of hydrogen was fitted and the reaction mixture was stirred at room temperature for 3h. The catalyst was filtered off and the filtrate was evaporated to dryness. Ethyl acetate was added to the residue and washed with saturated $NaHCO_3$ aqueous solution, water and brine. The water-phase was re-extracted with a mixture of ethyl acetate-MeOH 9-1. The combined organic phase was dried over $Na_2SO_4$, filtered and filtrate was removed and dried under reduced pressure to produce a white crude product. This crude product was purified by using preparative HPLC (Gylson GX-281 with a water-0.1%TFA / Acetonitrile gradient). To the fractions containing the product 200mg $NaHCO_3$ was added, the acetonitrile partially evaporated, and the product was extracted with ethyl acetate. The water-layer was re-extracted with 1x EtOAc-MeOH 9-1 and the combined-organic phases were dried over $Na_2SO_4$. The solvent was evaporated yielding the white product (138 mg, 0.166 mmol, 64.9 %)

LC-MS: M+H(+)=821.5 m/z

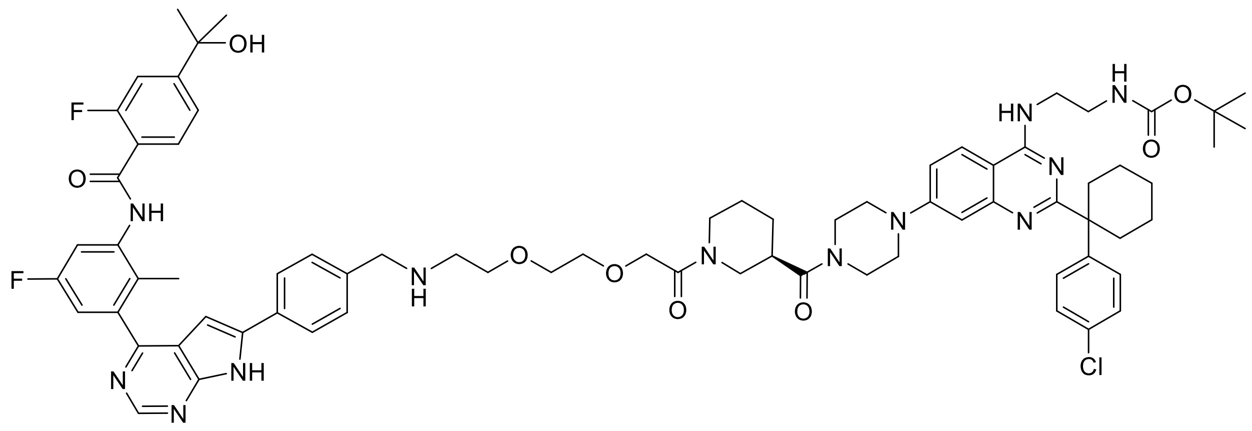

**tert-butyl (R)-(2-((2-(1-(4-chlorophenyl)cyclohexyl)-7-(4-(1-(2-(2-(2-((4-(4-(5-fluoro-3-(2-fluoro-4-(2-hydroxypropan-2-yl)benzamido)-2-methylphenyl)-7H-pyrrolo[2,3-d]pyrimidin-6-yl)benzyl)amino)ethoxy)ethoxy)acetyl)piperidine-3-carbonyl)piperazin-1-yl)quinazolin-4-yl)amino)ethyl)carbamate**

2-fluoro-N-(5-fluoro-3-(6-(4-formylphenyl)-7H-pyrrolo[2,3-d]pyrimidin-4-yl)-2-methylphenyl)-4-(2-hydroxypropan-2-yl)benzamide* (43 mg, 0.082 mmol, 1.0 Eq.) was suspended in 2mL MeOH and 2mL THF, tert-butyl (R)-(2-((7-(4-(1-(2-(2-(2-aminoethoxy)ethoxy)acetyl)piperidine-3-carbonyl)piperazin-1-yl)-2-(1-(4-chlorophenyl)cyclohexyl)quinazolin-4-yl)amino)ethyl)carbamate (69.7 mg, 0.132 mmol, 1.0 Eq.) and Sodium sulfate (58.0 mg, 0.408 mmol, 5.0 Eq.), acetic acid (7.58 µl, 0.132 mmol, 1.0 Eq.) were added at room temperature. The suspension was stirred at 23°C for 30 minutes. Sodium cyanoborohydride (26.2 mg, 0.408 mmol, 5.0 eq.) was added and the reaction mixture was stirred at room temperature for 22h. To the resulting yellow-brown cloudy solution 1mL of water was added and left stirring for 10 minutes. The reaction mixture was partitioned between water and ethyl acetate, the layers were separated and extracted with 1x15ml ethyl acetate, washed with 3x10ml water. The organic layers were combined, dried with $Na_2SO_4$, filtered, and concentrated under reduced pressure. The crude product was dissolved in 1mL DMSO/1mL MeOH and purified by using preparative HPLC (Gylson GX-281 with a water-0.1%TFA / Acetonitrile gradient). To the fractions containing the product 200mg $NaHCO_3$ was added, the acetonitrile was evaporated, and the product was extracted with ethyl acetate. The combined-organic phases were dried over $Na_2SO_4$, filtered, and concentrated to dryness to produce the slightly yellow product (50mg, 0.037 mmol, 45.5 %).

* synthesis described in patent WO2021/55295, page 195.

LC-MS: M+H(-)=1329.7/1331.7 m/z

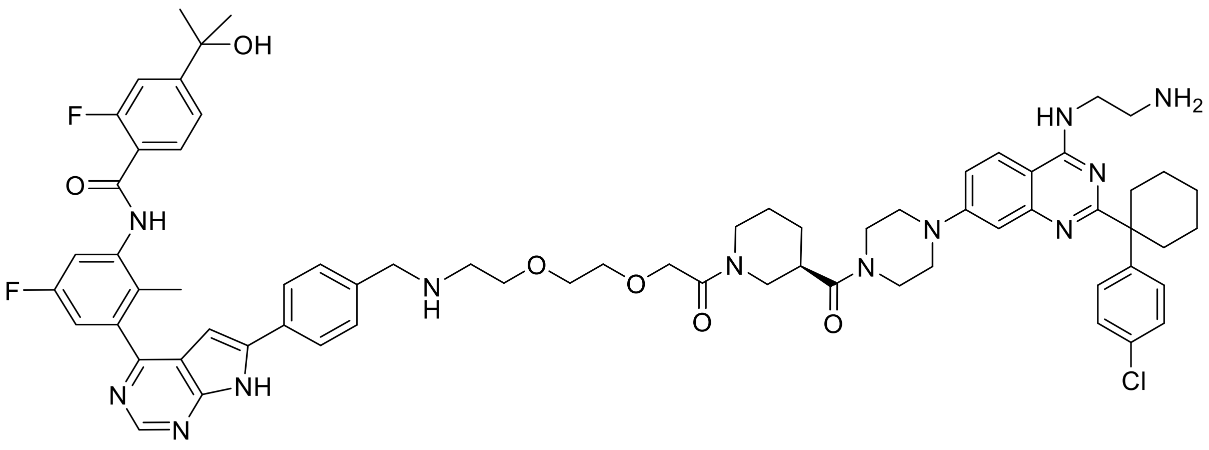

**DBt-6**

**(R)-N-(3-(6-(4-(((2-(2-(2-(3-(4-(4-((2-aminoethyl)amino)-2-(1-(4-chlorophenyl)cyclohexyl)quinazolin-7-yl)piperazine-1-carbonyl)piperidin-1-yl)-2-oxoethoxy)ethoxy)ethyl)amino)methyl)phenyl)-7H-pyrrolo[2,3-d]pyrimidin-4-yl)-5-fluoro-2-methylphenyl)-2-fluoro-4-(2-hydroxypropan-2-yl)benzamide**

A solution of tert-butyl (R)-(2-((2-(1-(4-chlorophenyl)cyclohexyl)-7-(4-(1-(2-(2-(2-((4-(4-(5-fluoro-3-(2-fluoro-4-(2-hydroxypropan-2-yl)benzamido)-2-methylphenyl)-7H-pyrrolo[2,3-d]pyrimidin-6-yl)benzyl)amino)ethoxy)ethoxy)acetyl)piperidine-3-carbonyl)piperazin-1-yl)quinazolin-4-yl)amino)ethyl)carbamate (50 mg, 0.038 mmol, 1.0 Eq.) in 3mL Dichloromethane was cooled in an ice bath before addition of TFA (0.087 ml, 1.126 mmol, 30.0 Eq.). The reaction mixture was stirred at room temperature for 2h resulting in a light-yellow solution. Dichloromethane was evaporated while cooling the reaction mix in an ice bath followed by the addition of 4mL water/Acetonitrile (1:1). The reaction mix was lyophilized resulting in a slightly yellow powder of DBt-6 (60mg, 0.030 mmol, 81 %). The identity and purity of the final compound were assessed both by high resolution MS as well as 1H and 13C NMR:

**$^{1}$H NMR** (600 MHz, DMSO-$D_6$) δ ppm (mixture of rotamers): 1.44 – 1.33 (m, 2H), 1.45 (s, 6H), 1.58 – 1.47 (m, 2H), 1.69 – 1.58 (m, 3H), 1.90 – 1.76 (m, 1H), 2.16 – 2.06 (m, 2H), 2.18 (s, 3H), 2.70 – 2.58 (m, 2H), 2.80 – 2.70 (m, 2H), 3.21 – 3.09 (m, 4H), 3.76 – 3.38 (m, 14H), 3.99 – 3.87 (m, 3H), 4.33 – 4.12 (m, 8H), 6.98 – 6.93 (m, 1H), 7.19 – 7.10 (m, 1H), 7.26 – 7.21 (m, 1H), 7.46 – 7.37 (m, 4H), 7.49 (d, *J* = 8.7 Hz, 2H), 7.60 (d, *J* = 8.0 Hz, 2H), 7.70 – 7.64 (m, 1H), 7.73 (t, *J* = 7.8 Hz, 1H), 8.12 – 8.02 (m, 5H), 8.17 – 8.12 (m, 1H), 8.88 (d, *J* = 3.1 Hz, 1H), 9.07 (s br, 2H), 9.85 (s br, 1H), 9.96 (d, *J* = 2.5 Hz, 1H), 12.83 (s br, 1H), 12.88 (s, 1H).

**$^{13}$C NMR** (150 MHz, DMSO-$D_6$) δ ppm (mixture of rotamers): 171.54, 171.17, 167.41, 165.71, 162.87, 159.42 (d, *J* = 241.6 Hz), 159.30, 159.10 (d, *J* = 248.1 Hz), 156.81 (d, *J* = 7.0 Hz), 156.76 (d, *J* = 2.5 Hz), 154.36, 153.31, 150.91, 142.86, 140.98, 139.35 (d, *J* = 8.6 Hz), 139.21, 138.44 (d, *J* = 10.8 Hz), 132.27, 131.81, 131.11, 130.66, 129.89 (d, *J* = 2.8 Hz), 128.54, 128.39, 126.51 (d, *J* = 3.1 Hz), 125.95, 125.38, 121.32 (d, *J* = 14.5 Hz), 120.76 (d, *J* = 2.8 Hz), 117.94, 116.32, 113.43 (d, *J* = 21.9 Hz), 112.58 (d, *J* = 24.0 Hz), 112.35 (d, *J* = 23.3 Hz), 102.04, 97.13, 70.61, 69.74, 69.51, 68.90 (d, *J* = 15.2 Hz), 65.56, 53.54, 49.75, 48.93, 46.60, 46.38, 45.87, 45.75, 44.61, 43.88, 43.81, 41.79, 40.41, 37.97, 37.91,37.72, 34.06, 31.56, 27.58, 27.27, 24.86, 24.66, 23.77, 22.57, 14.58.

**HR MS:** obs. m/z [M+H$^+$]: calculated for $C_{68}H_{78}N_{12}O_6ClF_2$, 1231.58184; found, 1231.58238; deviation: 0.4 ppm.

**Compound DBt-8 Synthetic Methods:**

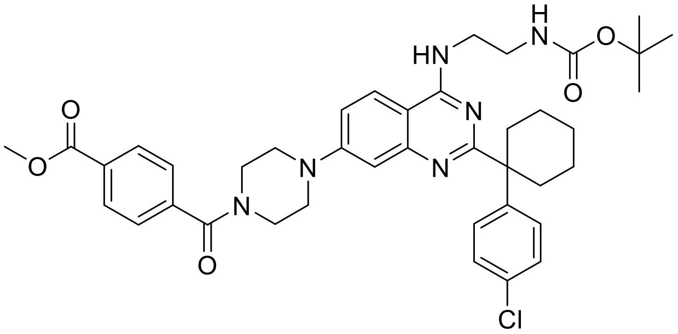

**methyl 4-(4-(4-((2-((tert-butoxycarbonyl)amino)ethyl)amino)-2-(1-(4-chlorophenyl)cyclohexyl)quinazolin-7-yl)piperazine-1-carbonyl)benzoate**

Under a argon atmosphere mono-Methyl terephtalate (111 mg, 0.599 mmol, 1.2 Eq., Fluka #86448), HATU (254 mg, 0.649 mmol, 1.3 Eq.) and DIPEA (0.261 ml, 1.497 mmol, 3.0 Eq.) were dissolved in 2.3ml DMA. The solution was stirred 15min at room temperature and then tert-butyl (2-((2-(1-(4-chlorophenyl)cyclohexyl)-7-(piperazin-1-yl)quinazolin-4-yl)amino)ethyl)carbamate (300 mg, 0.499 mmol, 1.0 Eq.) was added and stirred at room temperature for 1.5h resulting in a yellow-brown solution. The reaction mixture was partitioned between water and 90ml ethyl acetate. The phases were separated, and the organic phases were washed with 1x30ml water and 1x30ml brine. The organic layers were combined, dried with $Na_2SO_4$, filtered and volatiles were removed in vacuo to dryness yielding a brown resin of the crude product. This crude product was dissolved in Dichloromethane and adsorbed onto 3.4g Isolute and was purified by Combi Flash chromatography (detection at 254/280nm on a Combi Flash Rf 200 system with silica) with a gradient of DCM to DCM-MeOH 9-1, fractions containing the product were combined and concentrated under reduced pressure resulting in a crude yellow product. This was followed by an additional step of Combi Flash chromatography (detection at 254/280nm on a Combi Flash Rf 200 system with silica) with a gradient of hexane to DCM-ethyl acetate (90-10 to 80-20). Fractions containing product were evaporated to produce a white solid product (157 mg, 0.214 mmol, 42.8 %).

LC-MS: M+H(+)=728.5 m/z

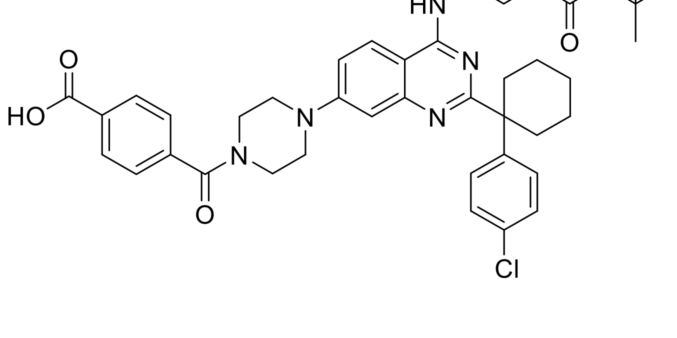

**4-(4-(4-((2-((tert-butoxycarbonyl)amino)ethyl)amino)-2-(1-(4-chlorophenyl)cyclohexyl)quinazolin-7-yl)piperazine-1-carbonyl)benzoic acid**

methyl 4-(4-(4-((2-((tert-butoxycarbonyl)amino)ethyl)amino)-2-(1-(4-chlorophenyl)cyclohexyl)quinazolin-7-yl)piperazine-1-carbonyl)benzoate (157 mg, 0.216 mmol, 1.0 Eq.) was dissolved in 1.7mL THF followed by the addition of Sodium hydroxide 2M (0.173 ml, 0.345 mmol, 1.6Eq.).The reaction mixture was stirred at room temperature for 22h. Additional Sodium hydroxide 2M (0.173 ml, 0.345 mmol, 1.6 Eq.) was added and stirred for another 24h at room temperature. Approx. 20 mL ethyl acetate and about 5mL water were added and the pH was adjusted with about 0.6mL 1N HCl to pH 4. The organic phase was washed with water and separated. The pH of the water phase was adjusted to ph2 with 1N HCl and further product was extracted with 20ml ethyl acetate-MeOH 9-1. The combined organic phase was filtered. The filtrate was dried over phase separator, the solvent partially evaporated under reduced pressure and hexane was added. The slightly yellow product was obtained by filtration and successive wash with hexane (120mg, 0.167mmol, 77%).

LC-MS: M+H(+)=714.5 m/z

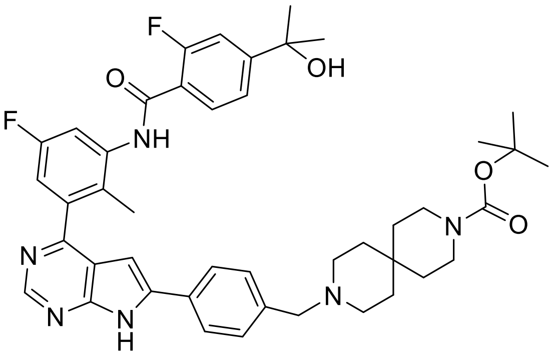


**tert-butyl 9-(4-(4-(5-fluoro-3-(2-fluoro-4-(2-hydroxypropan-2-yl)benzamido)-2-methylphenyl)-7H-pyrrolo[2,3-d]pyrimidin-6-yl)benzyl)-3,9-diazaspiro[5.5]undecane-3-carboxylate**

At room temperature, 3-N-Boc-3,9-diazaspiro-[5,5]undecane HCl (142 mg, 0.475 mmol, 1.0 Eq., CAS: 173405-2), TEA (0.196 mL, 1.405 mmol, 2.96 Eq.)) and 2-fluoro-N-(5-fluoro-3-(6-(4-formylphenyl)-7H-pyrrolo[2,3-d]pyrimidin-4-yl)-2-methylphenyl)-4-(2-hydroxypropan-2-yl)benzamide (250 mg, 0.475 mmol, 1.0 Eq) were suspended in 4mL MeOH. Zinc chloride (0.950 mL, 0.475 mmol, 1.0 Eq.) was added and the olive-green reaction mixture was stirred for 16h at room temperature resulting in an olive-green suspension. Then sodium cyanoborohydride (36.5 mg, 0.570 mmol,1.2 Eq.) was added and the reaction mixture was stirred for 4 days at room temperature producing a brown solution. The reaction mixture was filtered, washed with MeOH and filtrate was evaporated. The evaporated reaction mixture was partitioned between water and 90ml ethyl acetate. The phases were separated, and the organic phases were washed with 1x30ml water and 1x30ml brine. The organic layers were combined, dried with $Na_2SO_4$, filtered and volatiles were removed in vacuo to dryness yielding the yellow crude product (339mg, 0.430 mmol, 91 %).

LC-MS: [M+H]+ = 765.6 m/z

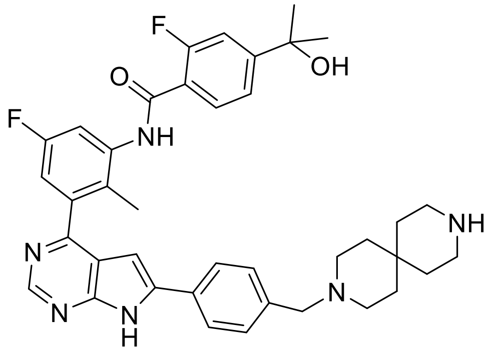


**N-(3-(6-(4-((3,9-diazaspiro[5.5]undecan-3-yl)methyl)phenyl)-7H-pyrrolo[2,3-d]pyrimidin-4-yl)-5-fluoro-2-methylphenyl)-2-fluoro-4-(2-hydroxypropan-2-yl)benzamide**

A solution of tert-butyl 9-(4-(4-(5-fluoro-3-(2-fluoro-4-(2-hydroxypropan-2-yl)benzamido)-2-methylphenyl)-7H-pyrrolo[2,3-d]pyrimidin-6-yl)benzyl)-3,9-diazaspiro[5.5]undecane-3-carboxylate (339 mg, 0.434 mmol, 1.0 Eq.) in 8mL Dichloromethane was cooled in an ice bath before addition of TFA (1.004 mL, 13.03 mmol, 30.0 Eq.). The reaction mixture was stirred at room temperature for 3h resulting in a light yellow solution. Dichloromethane was evaporated while cooling the reaction mix in an ice bath followed by the addition of 4mL MeOH. The crude product was purified by preparative HPLC on a Sunfire preparative C18 column (OBD 5µm. 30 X 100mm, Waters) eluting with acetonitrile in an aqueous solution of TFA (0.1 %). Fractions containing the title compound were combined and after the addition of 0.4 g NaHCO3 the reaction mixture was partitioned between water and 90ml ethyl acetate. The phases were separated, and the organic phases were washed with 1x30ml water and 1x30ml brine. The organic layers were combined, dried with $Na_2SO_4$, filtered and volatiles were removed in vacuo to dryness yielding the slightly yellow title product (168mg, 0.250 mmol, 57.6 %).

LC-MS: [M+H]+ = 665.5 m/z

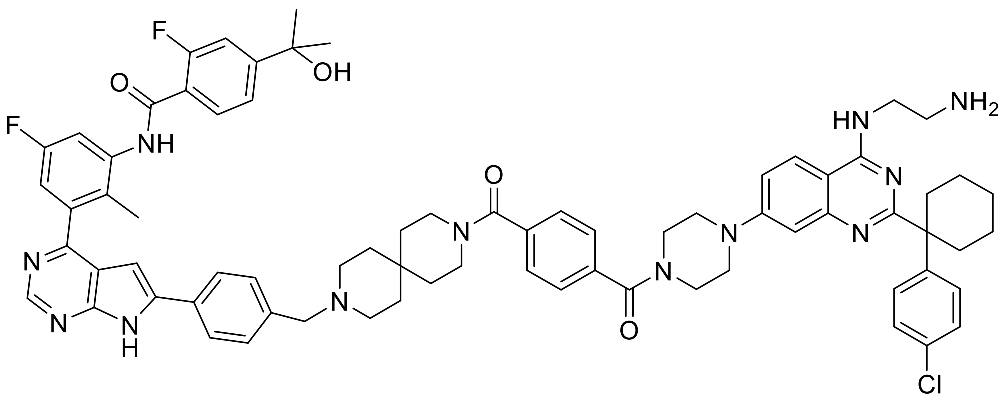


**DBt-8**

**N-(3-(6-(4-((9-(4-(4-(4-((2-aminoethyl)amino)-2-(1-(4-chlorophenyl)cyclohexyl)quinazolin-7-yl)piperazine-1-carbonyl)benzoyl)-3,9-diazaspiro[5.5]undecan-3-yl)methyl)phenyl)-7H-pyrrolo[2,3-d]pyrimidin-4-yl)-5-fluoro-2-methylphenyl)-2-fluoro-4-(2-hydroxypropan-2-yl)benzamide**

Under a N2 atmosphere, 4-(4-(4-((2-((tert-butoxycarbonyl)amino)ethyl)amino)-2-(1-(4-chlorophenyl)cyclohexyl)quinazolin-7-yl)piperazine-1-carbonyl)benzoic acid (115 mg, 0.162 mmol, 1.17 Eq.) was dissolved in 0.8mL DMA and HATU (70.5 mg, 0.180 mmol, 1.3 Eq.) and DIPEA (0.073 ml, 0.415 mmol, 3.0 Eq.) were added. The yellow suspension was stirred at room temperature for 10 minutes. After this activation, a solution of N-(3-(6-(4-((3,9-diazaspiro[5.5]undecan-3-yl)methyl)phenyl)-7H-pyrrolo[2,3-d]pyrimidin-4-yl)-5-fluoro-2-methylphenyl)-2-fluoro-4-(2-hydroxypropan-2-yl)benzamide (92 mg, 0.138 mmol, 1.0 Eq.) in 0.8mL DMA was added and the solution was stirred at room temperature for 90minutes resulting in a yellow solution. The reaction mixture was partitioned between water and 90ml ethyl acetate. The phases were separated, and the organic phases were washed with 1x30ml water and 1x30ml brine. The organic layers were combined, dried with $Na_2SO_4$, filtered and volatiles were removed in vacuo to dryness to yield the yellow resin of the crude title product. This crude product was dissolved in a mixture of 1.5mL DMSO/ 2mL MeOH and was purified by preparative HPLC on a Sunfire preparative C18 column (OBD 5µm. 30 X 100mm, Waters) eluting with acetonitrile in an aqueous solution of TFA (0.1 %). Fractions containing the title compound were combined. After the addition of 65 mg $NaHCO_3$ the reaction mixture was partitioned between water and 90ml ethyl acetate. The phases were separated, and the organic phases were washed with 1x30ml water and 1x30ml brine. The organic layers were combined, dried with $Na_2SO_4$, filtered and volatiles were removed in vacuo to dryness yielding the slightly yellow tert-butyl (2-((2-(1-(4-chlorophenyl)cyclohexyl)-7-(4-(4-(9-(4-(4-(5-fluoro-3-(2-fluoro-4-(2-hydroxypropan-2-yl)benzamido)-2-methylphenyl)-7H-pyrrolo[2,3-d]pyrimidin-6-yl)benzyl)-3,9-diazaspiro[5.5]undecane-3-carbonyl)benzoyl)piperazin-1-yl)quinazolin-4-yl)amino)ethyl)carbamate (108mg, 0.075 mmol, 53.9 %).

This was directly deprotected to yield the final product. Therefore, a solution of tert-butyl (2-((2-(1-(4-chlorophenyl)cyclohexyl)-7-(4-(4-(9-(4-(4-(5-fluoro-3-(2-fluoro-4-(2-hydroxypropan-2-yl)benzamido)-2-methylphenyl)-7H-pyrrolo[2,3-d]pyrimidin-6-yl)benzyl)-3,9-diazaspiro[5.5]undecane-3-carbonyl)benzoyl)piperazin-1-yl)quinazolin-4-yl)amino)ethyl)carbamate (108mg, 0.075 mmol, 1.0 Eq.) in 3mL Dichloromethane was cooled in an ice bath before addition of TFA (0.173 ml, 2.239 mmol, 30.0 Eq.). The reaction mixture was stirred at room temperature for 2.5h resulting in a light-yellow solution. Dichloromethane was evaporated while cooling the reaction mix in an ice bath followed by the addition of 1.5mL acetonitrile. This was further purified by preparative HPLC on a Sunfire preparative C18 column

(OBD 5µm. 30 X 100mm, Waters) eluting with acetonitrile in an aqueous solution of TFA (0.1 %). Fractions containing the title compound were combined. After the addition of 99 mg $NaHCO_3$ the ethyl acetate was evaporated, and the resulting white suspension was filtered. The white solid was washed with water and dried under reduced pressure to yield the title compound DBt-8 (55 mg, 0.043 mmol, 57.9 %). The identity and purity of the final compound were confirmed by high resolution MS as well as 1H and 13C NMR:

**$^{1}$H NMR** (600 MHz, DMSO-$D_6$) δ ppm: 1.44 – 1.19 (m, 2H), 1.46 (s, 6H), 1.78 – 1.47 (m, 6H), 1.92 (d, *J* = 13.6 Hz, 2H), 2.16 – 2.06 (m, 2H), 2.18 (s, 3H), 2.76 (d, *J* = 13.0 Hz, 2H), 3.20 – 2.96 (m, 4H), 3.39 – 3.20 (m, 5H), 3.70 – 3.39 (m, 6H), 3.78 (s, 3H), 3.94 (s, 2H), 4.37 (s, 2H), 6.99 (d, *J* = 2.2 Hz, 1H), 7.17 (s, 1H), 7.25 (dd, *J* = 8.8, 2.8 Hz, 1H), 7.55 – 7.37 (m, 7H), 7.60 (d, *J* = 7.9 Hz, 2H), 7.67 (d, *J* = 10.1 Hz, 1H), 7.73 (t, *J* = 7.8 Hz, 1H), 8.09 – 7.99 (m, 3H), 8.11 (d, *J* = 7.9 Hz, 2H), 8.15 (d, *J* = 9.4 Hz, 1H), 8.90 (s, 1H), 9.73 (s br, 1H), 9.84 (s br, 1H), 9.96 (d, *J* = 2.5 Hz, 1H), 12.83 (s br, 1H), 12.92 (s, 1H).

**$^{13}$C NMR** (150 MHz, DMSO-$D_6$) δ ppm: 168.58, 168.17, 165.75, 162.88, 159.40 (d, J = 250 Hz), 159.32, 159.11 (d, J = 248 Hz), 158.63, 158.28, 156.84, 156.79, 154.41, 153.34, 150.95, 139.28 (d, J = 9.0 Hz), 139.10, 138.45 (d, J = 10.8 Hz), 137.49, 136.38, 131.80, 131.64, 130.20, 129.90 (d, J = 2.9 Hz), 128.56, 128.39, 127.21, 126.78, 126.53 (d, J = 2.6 Hz), 126.11, 125.44, 121.33 (d, J = 14.5 Hz), 120.77 (d, J = 2.8 Hz), 117.94, 116.40, 113.46 (d, J = 21.5 Hz), 112.63 (d, J = 23.8 Hz), 112.36 (d, J = 23.3 Hz), 97.43, 70.62 (d, J = 1.4 Hz), 58.28, 48.95, 47.19, 46.31, 42.84, 37.92, 37.20, 34.08, 31.56, 31.43, 29.17, 24.86, 22.57, 14.59.

**HR MS:** obs. m/z [M+H$^+$]: calculated for $C_{73}H_{78}N_{12}O_4ClF_2$, 1259.59201; found, 1259.59128; deviation: 0.6 ppm.

**Compound DBt-10 Synthetic Methods:**

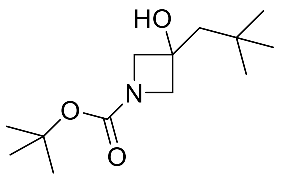

**tert-butyl 3-hydroxy-3-neopentylazetidine-1-carboxylate**

Lanthanium(III)chlorid bis(lithiumchlorid)complex solution in THF (170 mL, 1.0 Eq., 102 mmol) was slowly added to 1-Boc-azetidin-3-on (17.7999g, 1.0 Eq., 102 mmol) under Argon atmosphere at room temperature and reaction was left stirring for 1.5h at room temperature. 2,2-Dimethylpropylmagnesiumchlorid (107 mL, 1.05 Eq., 107 mmol) was slowly added over 30 min and stirred for 1h. reaction was stopped by addition of 100mL aqueous, saturated NH4Cl solution and 100ml H2O on ice and left stirring for 15min. After dilution with H2O and Ethyl acetate and product extracted with Ethyl acetate, washed with brine, dried over sodium sulfate and concentrated under reduced pressure. After chromatographic separation silica gel column chromatography with a ethyl acetate- heptane gradient, tert-butyl 3-hydroxy-3-neopentylazetidine-1-carboxylate was isolated as white product (18.185g, 75mmol, 73%).

**LCMS**: m/z 242.3 [M-H]-

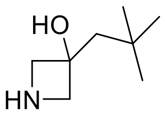

**3-neopentylazetidin-3-ol**

To a solution of tert-butyl 3-hydroxy-3-neopentylazetidine-1-carboxylate (13.183g, 1.0 Eq, 54.2 mmol,) in DCM (182.4 ml) TFA was slowly added (91.2 ml, 1.0 Eq., 54.2 mmol) at 0°C. The clear colorless solution was stirred for 30min at rt. TLC (Heptane/EA 3:1 with Ninhydrine) showed the complete conversion to product. TFA and DCM was evaporated on rotavapor and afterwards the yellow oil was solved two times in toluene and evaporated to dryness. The resulting yellow oil was triturated with ethyl ether and cyclohexylamine at 0°C, afterwards the white crystals of 3-neopentylazetidin-3-ol were filtered and washed two times with cyclohexylamine (13.62g, 52.9 mmol, 98 % yield).

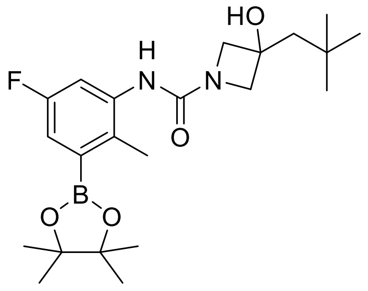

**N-(5-fluoro-2-methyl-3-(4,4,5,5-tetramethyl-1,3,2-dioxaborolan-2-yl)phenyl)-3-hydroxy-3-neopentylazetidine-1-carboxamide**

To a solution of 5-fluoro-2-methyl-3-(4,4,5,5-tetramethyl-1,3,2-dioxaborolan-2-yl)aniline (3.6 g, 1.0 Eq., 14.34 mmol, Merck) and DIPEA (10.02 mL, 4.0 Qu., 57.3 mmol) in $CH_2Cl_2$ (Volume: 40 mL, Ratio: 1.000) Phosgen 20% in toluene was added (9.05 mL,1.2 Eq., 17.20 mmol) slowly at 0 °C. The resulting solution was stirred for 10 min at 0°C, then transferred to a stirring solution of 3-neopentylazetidin-3-ol (4.06 g, 1.1 Eq., 15.77 mmol) in $CH_2Cl_2$ (Volume: 40.0 mL, Ratio: 1.000) at 0 °C. The resulting mixture was stirred for 1 h at 0 °C. The reaction was monitored by LC-MS and indicated a complete conversion. The crude product was triturated in $CH_2Cl_2$/TBME 1:5, then filtered off and dried at high vacuum. The remaining material was concentrated and crystallized again in TBME which resulted in additional product. N-(5-fluoro-2-methyl-3-(4,4,5,5-tetramethyl-1,3,2-dioxaborolan-2-yl)phenyl)-3-hydroxy-3-neopentylazetidine-1-carboxamide was isolated as white crystals (5g, 11.90mmol, 83%).

**1H NMR:** (400 MHz, DMSO-d6) δ 7.73 (s, 1H), 7.40 - 7.22 (m, 1H), 7.14 - 6.98 (m, 1H), 5.45 (s, 1H), 3.83 (dd, J = 45.7, 8.5 Hz, 4H), 2.29 (s, 3H), 1.64 (s, 2H), 1.29 (s, 13H), 0.96 (s, 9H).

LCMS: m/z 421 [M+H]+, m/z 419 [M-H]-

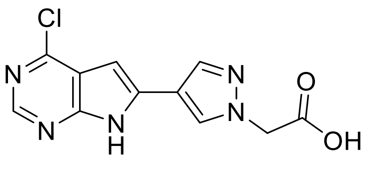

**2-(4-(4-chloro-7H-pyrrolo[2,3-d]pyrimidin-6-yl)-1H-pyrazol-1-yl)acetic acid**

4-chloro-6-iodo-7H-pyrrolo[2,3-d]pyrimidine (1.50 g, 1.0 Eq., 5.37 mmol, CAS number: 876343-10-1, Merck) was dissolved in 50mL Dioxan:H2O (1:1). -(Ethoxycarbonylmethyl)pyrazole-4-boronic acid PE (1.804 g, 1.2Eq., 6.44 mmol),

Cesium carbonate (4.36 g, 2.5 Eq., 13.42 mmol) and $PdCl_2$(PPh2)ferrocene in DCM (0.438 g, 0.1 Eq., 0.537 mmol) were added and reaction was left stirring at 100°C for 1h. After cooling down, approx. 80mL ethyl acetate was added and the water phase was separated, of which the pH was subsequently adjusted to pH2-3 with 0.1M HCl. A further extraction was performed with ethyl acetate, the organic phase was washed once with H2O and then evaporated completely under reduced pressure resulting in the 2-(4-(4-chloro-7H-pyrrolo[2,3-d]pyrimidin-6-yl)-1H-pyrazol-1-yl)acetic acid product. An additional extraction with Butanol of the remaining aqueous phase produced further product and was combined with the already extracted product resulting in the final slightly brown colored 2-(4-(4-chloro-7H-pyrrolo[2,3-d]pyrimidin-6-yl)-1H-pyrazol-1-yl)acetic acid (1.815 g, quantitative, 6.55 mmol)

LCMS: m/z 278.1 [M+H]+, m/z 276.0 [M-H]-

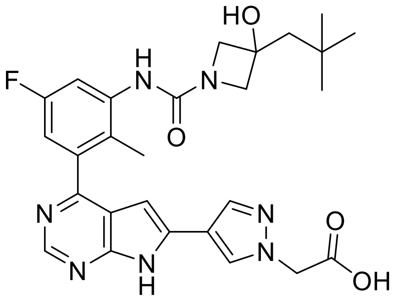


**2-(4-(4-(5-fluoro-3-(3-hydroxy-3-neopentylazetidine-1-carboxamido)-2-methylphenyl)-7H-pyrrolo[2,3-d]pyrimidin-6-yl)-1H-pyrazol-1-yl)acetic acid**

2-(4-(4-chloro-7H-pyrrolo[2,3-d]pyrimidin-6-yl)-1H-pyrazol-1-yl)acetic acid (0.792 g, 1.05 Eq., 2.423 mmol) was dissolved in 25mL Acetonitrile, N-(5-fluoro-2-methyl-3-(4,4,5,5-tetramethyl-1,3,2-dioxaborolan-2-yl)phenyl)-3-hydroxy-3-neopentylazetidine-1-carboxamide (1.0 g, 1.0 Eq., 2.308 mmol), Potassium carbonate (8.31 ml, 3.6 Eq., 8.31 mmol) and Pd(dppf)$Cl_2$ (0.253 g, 0.15 Eq., 0.346 mmol) were added and stirred at 100℃ for 30 minutes. After cooling down, approx. 40mL of Ethyl acetate were added and the aqueous phase was separated, which contained all the product. The pH of the aqueous phase was adjusted to 2-3 with 0.1M HCl and the product was extracted with ethyl acetate twice, the combined organic phases were washed with brine (50 mL), dried over sodium sulfate and concentrated resulting in 2-(4-(4-(5-fluoro-3-(3-hydroxy-3-neopentylazetidine-1-carboxamido)-2-methylphenyl)-7H-pyrrolo[2,3-d]pyrimidin-6-yl)-1H-pyrazol-1-yl)acetic acid (1.112g, 0.903 mmol, 39%)

LCMS: m/z 536.3 [M+H]+, m/z 534.2 [M-H]-

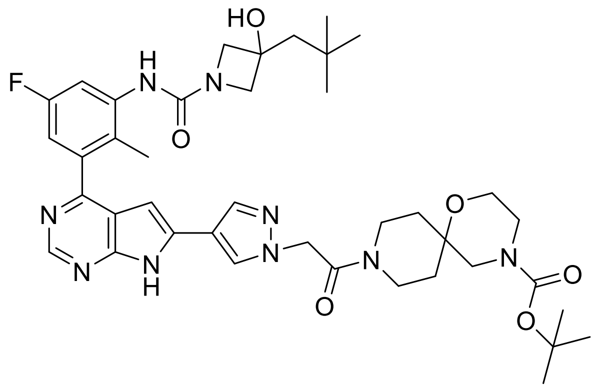

**tert-butyl 9-(2-(4-(4-(5-fluoro-3-(3-hydroxy-3-neopentylazetidine-1-carboxamido)-2-methylphenyl)-7H-pyrrolo[2,3-d]pyrimidin-6-yl)-1H-pyrazol-1-yl)acetyl)-1-oxa-4,9-diazaspiro[5.5]undecane-4-carboxylate**

2-(4-(4-(5-fluoro-3-(3-hydroxy-3-neopentylazetidine-1-carboxamido)-2-methylphenyl)-7H-pyrrolo[2,3-d]pyrimidin-6-yl)-1H-pyrazol-1-yl)acetic acid (300 mg, 1.0 Eq., 0.504 mmol), HATU (237 mg, 1.2 Eq., 0.605 mmol) and DIEA (0.308 ml, 3.5 Eq., 1.764 mmol) were pre-activated in 3mL DMA. Than a solution of tert-butyl 1-oxa-4,9-diazaspiro[5.5]undecane-4-carboxylate (166 mg, 1.1 Eq., 0.555 mmol, CAS: 1023595-11-0) in 2 ml DMA was added to the pre-activated solution and stirred at room temperature, after 30 minutes additional 50% HATU and DIEA added (0.6 and 1.75 Eq., respectively) and stirred again for 10 minutes. Afterwards approx. 50mL of Ethyl acetate were added and the organic phase was twice extracted with H2O. The combined organic phases were subsequentially evaporated, and the product was further purified using a Combi-Flash system (40g silica column) with a gradient of 90% DCM + 10% Methanol and DCM. The fractions containing the product were combined and evaporated under reduced pressure resulting in tert-butyl 9-(2-(4-(4-(5-fluoro-3-(3-hydroxy-3-neopentylazetidine-1-carboxamido)-2-methylphenyl)-7H-pyrrolo[2,3-d]pyrimidin-6-yl)-1H-pyrazol-1-yl)acetyl)-1-oxa-4,9-diazaspiro[5.5]undecane-4-carboxylate (0.2g, 0.128 mmol, 25%).

LCMS: m/z 774.5 [M+H]+, m/z 772.4 [M-H]-

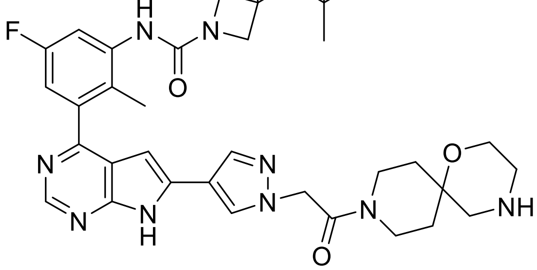

**N-(5-fluoro-2-methyl-3-(6-(1-(2-oxo-2-(1-oxa-4,9-diazaspiro[5.5]undecan-9-yl)ethyl)-1H-pyrazol-4-yl)-7H-pyrrolo[2,3-d]pyrimidin-4-yl)phenyl)-3-hydroxy-3-neopentylazetidine-1-carboxamide**

Tert-butyl 9-(2-(4-(4-(5-fluoro-3-(3-hydroxy-3-neopentylazetidine-1-carboxamido)-2-methylphenyl)-7H-pyrrolo[2,3-d]pyrimidin-6-yl)-1H-pyrazol-1-yl)acetyl)-1-oxa-4,9-diazaspiro[5.5]undecane-4-carboxylate (200 mg, 1.0 Eq., 0.258 mmol) was dissolved in 1mL DCM, mixed with 2,2,2-trifluoroacetic acid (0.594 ml, 30.0 Eq., 7.75 mmol) and stirred at room temperature for 60 minutes. The product was isolated by the addition of approx. 50mL Ethyl acetate and extraction with once a 5% bicarbonate solution and twice with H2O. The aqueous phase was additionally extracted with Butanol. All organic phases were combined, and the solvent evaporated under reduced pressure. The product was directly used for the following synthesis steps without further purification (210mg, quantitative)

LCMS: m/z 674.5 [M+H]+, m/z 672.3 [M-H]-

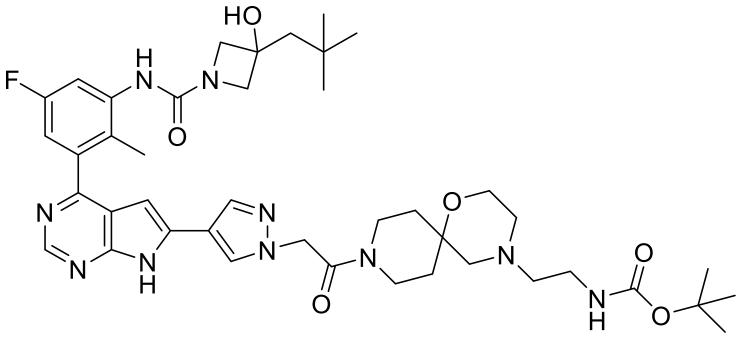


**tert-butyl (2-(9-(2-(4-(4-(5-fluoro-3-(3-hydroxy-3-neopentylazetidine-1-carboxamido)-2-methylphenyl)-7H-pyrrolo[2,3-d]pyrimidin-6-yl)-1H-pyrazol-1-yl)acetyl)-1-oxa-4,9-diazaspiro[5.5]undecan-4-yl)ethyl)carbamate**

N-(5-fluoro-2-methyl-3-(6-(1-(2-oxo-2-(1-oxa-4,9-diazaspiro[5.5]undecan-9-yl)ethyl)-1H-pyrazol-4-yl)-7H-pyrrolo[2,3-d]pyrimidin-4-yl)phenyl)-3-hydroxy-3-neopentylazetidine-1-carboxamide (174 mg, 1.0 Eq., 0.258 mmol) was dissolved in 2mL MeOH/acetic acid and mixed with N-Boc-2-Aminoacetaldehyde (51.9 mg, 1.20Eq., 0.310 mmol, CAS: 89711-08-0) and stirred for 1h at room temperature. Sodium cyanoborohydride (24.32 mg, 1.5 Eq., 0.387 mmol) was added and stirred overnight at room temperature. The reaction was mixed with approx. 50mL Ethyl acetate and extracted twice with water. the combined organic phases were washed with brine (200 mL), dried over sodium sulfate and concentrated under reduced pressure and the resulting product was directly used for the following reaction without further purification (0.246g, 0.139 mmol, 45%).

LCMS: m/z 817.5 [M+H]+, m/z 815.4 [M-H]-

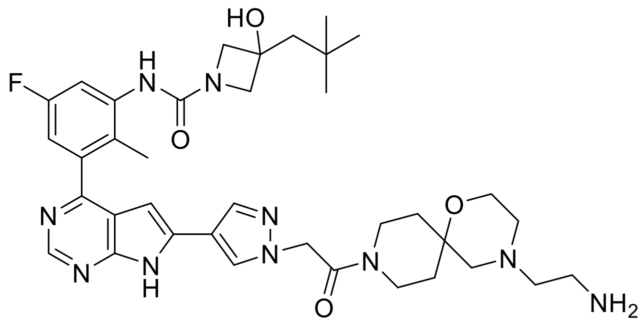

**N-(3-(6-(1-(2-(4-(2-aminoethyl)-1-oxa-4,9-diazaspiro[5.5]undecan-9-yl)-2-oxoethyl)-1H-pyrazol-4-yl)-7H-pyrrolo[2,3-d]pyrimidin-4-yl)-5-fluoro-2-methylphenyl)-3-hydroxy-3-neopentylazetidine-1-carboxamide**

Tert-butyl (2-(9-(2-(4-(4-(5-fluoro-3-(3-hydroxy-3-neopentylazetidine-1-carboxamido)-2-methylphenyl)-7H-pyrrolo[2,3-d]pyrimidin-6-yl)-1H-pyrazol-1-yl)acetyl)-1-oxa-4,9-diazaspiro[5.5]undecan-4-yl)ethyl)carbamate (246 mg, 1.0 Eq., 0.301 mmol) was dissolved in 1mL DCM, mixed with 2,2,2-trifluoroacetic acid (0.692 ml, 30 Eq., 9.03 mmol) and stirred at room temperature for 30 minutes. After the addition of approx. 50mL Butanol the mixture was extracted twice with water. The combined organic phases were washed with brine (50 mL), dried over sodium sulfate, and concentrated under reduced pressure and the resulting product was used without further purification for the following synthesis step (0.204g, 0.285 mmol)

LCMS: m/z 717.6 [M+H]+, m/z 715.4 [M-H]-

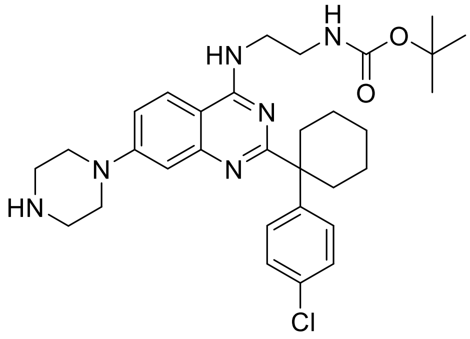

**tert-butyl (2-((2-(1-(4-chlorophenyl)cyclohexyl)-7-(piperazin-1-yl)quinazolin-4-yl)amino)ethyl)carbamate**

Tert-butyl (2-((7-(4-acetylpiperazin-1-yl)-2-(1-(4-chlorophenyl)cyclohexyl)quinazolin-4-yl)amino)ethyl)carbamate (4.074 g, 1.0 Eq., 5.57 mmol) was dissolved in 10mL Ethanol, mixed with 2M Potassium hydroxide (27.8 ml, 10 Eq., 55.7 mmol) and stirred at 80°C for 48h. The reaction was mixed with approx. 80mL Ethyl acetate and the organic phase was extracted twice with water followed by the evaporation of the organic solvents under reduced pressure. The product was purified using a Combi-

Flash chromatography system with a gradient of 90% DCM + 10% Methanol and DCM. Fractions containing the product were pooled and the solvent was evaporated under reduced pressure resulting in slightly brown powder (1.78g, 3.15 mmol, 57%)

LCMS: m/z 565.3 [M+H]+, m/z 563.3 [M-H]-

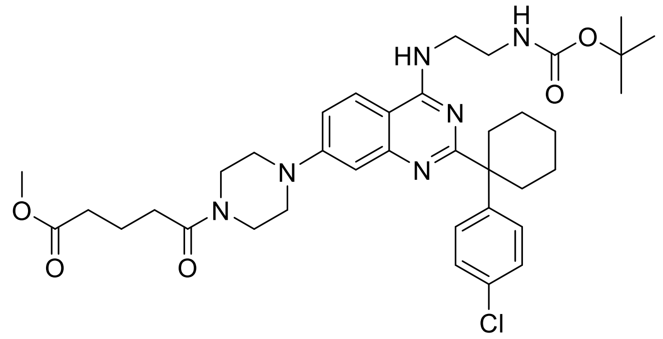

**methyl 5-(4-(4-((2-((tert-butoxycarbonyl)amino)ethyl)amino)-2-(1-(4-chlorophenyl)cyclohexyl)quinazolin-7-yl)piperazin-1-yl)-5-oxopentanoate**

Tert-butyl (2-((2-(1-(4-chlorophenyl)cyclohexyl)-7-(piperazin-1-yl)quinazolin-4-yl)amino)ethyl)carbamate (200 mg, 1.0 Eq., 0.354 mmol) and Triethylamine (0.075 ml, 1.5 Eq., 0.531 mmol) were dissolved in 2mL DCM followed by the addition of methyl 5-chloro-5-oxopentanoate (0.054 ml, 1.1 Eq., 0.389 mmol) (CAS-Nr: 1501-26-4). This reaction was stirred at room temperature for 1h. The reaction was mixed with approx. 50mL Ethyl acetate and the organic phase was extracted twice with water followed by the evaporation of the organic solvents under reduced pressure. The product was used further without additional purification steps. (0.240g, 0.346 mmol, 98%)

LCMS: m/z 693.0 [M+H]+

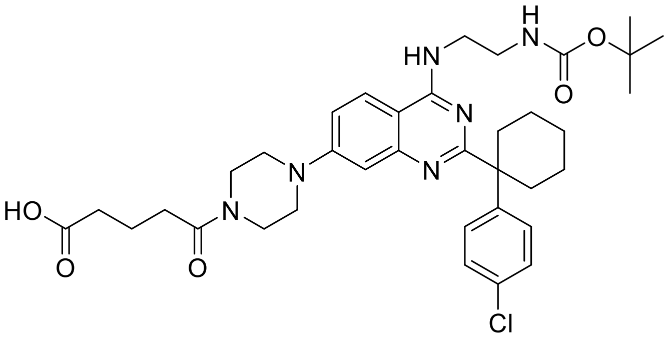

**5-(4-(4-((2-((tert-butoxycarbonyl)amino)ethyl)amino)-2-(1-(4-chlorophenyl)cyclohexyl)quinazolin-7-yl)piperazin-1-yl)-5-oxopentanoic acid**

methyl 5-(4-(4-((2-((tert-butoxycarbonyl)amino)ethyl)amino)-2-(1-(4-chlorophenyl)cyclohexyl)quinazolin-7-yl)piperazin-1-yl)-5-oxopentanoate (240 mg, 1.0 Eq., 0.329 mmol) was dissolved in THF and mixed with Sodium hydroxide (0.247 ml, 1.5 Eq., 0.493 mmol) and stirred at room temperature for 6h. The reaction was mixed with approx. 50mL Ethyl acetate and the organic phase was extracted with water, however the product remained in the aqueous phase. Therefore, the pH of the aqueous phase was adjusted to pH 2-3 with 0.1M HCl and extracted with Butanol. After washing the organic phase once with water, it was evaporated under reduced pressure. The free acid was used without further purification steps (0.224g, 0.159 mmol, 48%)

LCMS: m/z 679.4 [M+H]+, m/z 677.3 [M-H]-

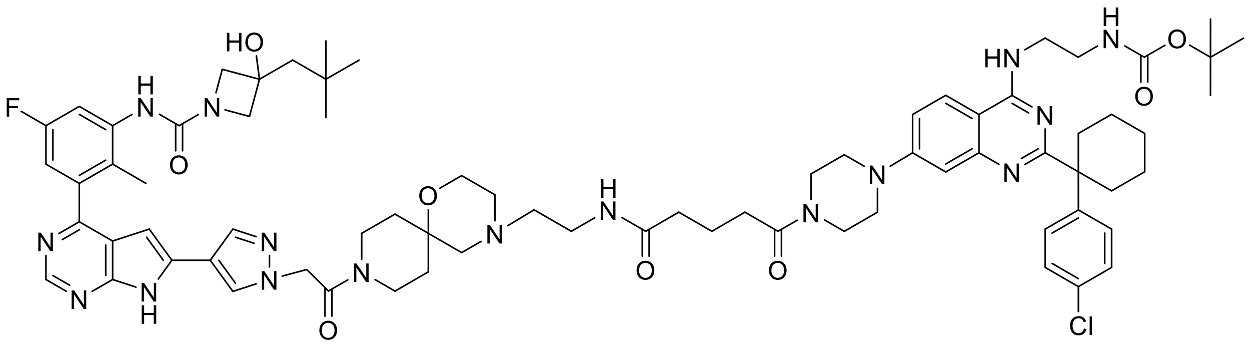

**tert-butyl (2-((2-(1-(4-chlorophenyl)cyclohexyl)-7-(4-(5-((2-(9-(2-(4-(4-(5-fluoro-3-(3-hydroxy-3-neopentylazetidine-1-carboxamido)-2-methylphenyl)-7H-pyrrolo[2,3-d]pyrimidin-6-yl)-1H-pyrazol-1-yl)acetyl)-1-oxa-4,9-diazaspiro[5.5]undecan-4-yl)ethyl)amino)-5-oxopentanoyl)piperazin-1-yl)quinazolin-4-yl)amino)ethyl)carbamate**

N-(3-(6-(1-(2-(4-(2-aminoethyl)-1-oxa-4,9-diazaspiro[5.5]undecan-9-yl)-2-oxoethyl)-1H-pyrazol-4-yl)-7H-pyrrolo[2,3-d]pyrimidin-4-yl)-5-fluoro-2-methylphenyl)-3-hydroxy-3-neopentylazetidine-1-carboxamide (80 mg, 1.0 Eq., 0.113 mmol), HATU (66.5 mg, 1.5 Eq., 0.170 mmol) and DIEA (0.079 ml, 4.0 Eq., 0.452 mmol) were preactivated in 2mL DMA for 5 minutes. Then a solution of 5-(4-(4-((2-((tert-butoxycarbonyl)amino)ethyl)amino)-2-(1-(4-chlorophenyl)cyclohexyl)quinazolin-7-yl)piperazin-1-yl)-5-oxopentanoic acid (81 mg, 1.0 Eq., 0.113 mmol) in 1mL DMA was added and stirred for 60 minutes. Since the reaction was not complete, additional 50% HATU and DIEA (0.75 Eq. and 2.0 Eq., respectively) were added and the reaction was stirred overnight at room temperature. The reaction was mixed with approx. 50mL Ethyl acetate and the organic phase was extracted twice with water, the combined organic phases were washed with brine (50 mL), dried over sodium sulfate followed by the evaporation of the organic solvents under reduced pressure. The product was purified using a Gilson preparative HPLC system with a C18 column and a gradient of 5% - 20 Min - 95%B (A=H2O + 5% TFA , B=Acetonitrile). The fractions containing the product were pooled and the acetonitrile was removed by distillation. The pH of the remaining aqueous solution was adjusted to 9 by addition of a 5% solution of bicarbonate followed by an extraction with Ethyl acetate. The organic phase was washed twice with water, combined organic phases were washed with brine (20 mL), dried over sodium sulfate and subsequentially evaporated under reduced pressure resulting in the product (0.092g, 0.068 mmol, 59%)

LCMS: m/z 1379.6 [M+H]+, m/z 1377.6 [M-H]-

DBt-10

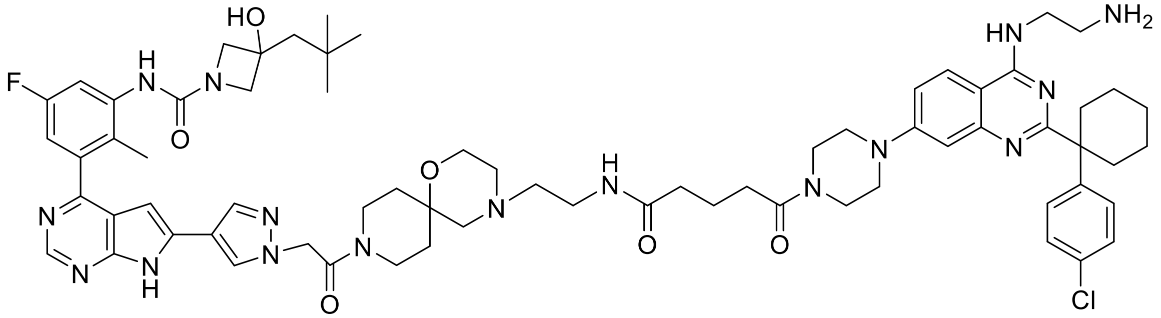

**DBt-10**

**N-(3-(6-(1-(2-(4-(2-(5-(4-(4-((2-aminoethyl)amino)-2-(1-(4-chlorophenyl)cyclohexyl)quinazolin-7-yl)piperazin-1-yl)-5-oxopentanamido)ethyl)-1-oxa-4,9-diazaspiro[5.5]undecan-9-yl)-2-oxoethyl)-1H-pyrazol-4-yl)-7H-pyrrolo[2,3-d]pyrimidin-4-yl)-5-fluoro-2-methylphenyl)-3-hydroxy-3-neopentylazetidine-1-carboxamide**

Tert-butyl (2-((2-(1-(4-chlorophenyl)cyclohexyl)-7-(4-(5-((2-(9-(2-(4-(4-(5-fluoro-3-(3-hydroxy-3-neopentylazetidine-1-carboxamido)-2-methylphenyl)-7H-pyrrolo[2,3-d]pyrimidin-6-yl)-1H-pyrazol-1-yl)acetyl)-1-oxa-4,9-diazaspiro[5.5]undecan-4-yl)ethyl)amino)-5-oxopentanoyl)piperazin-1-yl)quinazolin-4-yl)amino)ethyl)carbamate (92 mg, 1.0 Eq., 0.067 mmol) was dissolved in 1mL DCM, mixed with 2,2,2-trifluoroacetic acid (0.256 ml, 50 Eq., 3.34 mmol) and stirred for 60 minutes at room temperature followed by the evaporation of the solvents. The product was purified using a Gilson preparative HPLC system with a C18 column and a gradient of 5% - 20 Min - 70%B (A=H2O + 5% TFA , B=Acetonitrile). The fractions containing the product were pooled and the acetonitrile was removed by distillation. The pH of the remaining aqueous solution was adjusted to 9 by addition of a 5% solution of bicarbonate followed by an extraction with Ethyl acetate. The organic phase was washed with brine (50 mL), dried over sodium sulfate and concentrated resulting in DBt-10 (58mg, 0.045mmol, 68%)

The identity and purity of the final compound were confirmed both by high resolution MS as well as 1H and 13C NMR:

**$^1$H NMR** (600 MHz, DMSO-D$_6$) δ ppm: 0.98 (s, 9H), 1.34 - 1.57 (m, 4H), 1.59 – 1.65 (m, 2H), 1.66 (s, 2H), 1.76 (p, J = 7.5 Hz, 2H), 2.07 (s, 3H), 2.09 – 2.15 (m, 2H), 2.18 (t, J = 7.5 Hz, 2H), 2.38 (t, J = 7.3 Hz, 2H), 2.70 – 2.80 (m, 2H), 2.83 – 3.22 (m, 10H), 3.32 – 3.67 (m, 12H), 3.68 – 3.77 (m, 2H), 3.79 – 4.05 (m, 10H), 5.19 (d, J = 16.1 Hz, 1H), 5.26 (d, J = 16.1 Hz, 1H), 6.53 (s, 1H), 7.05 (dd, J = 8.8, 2.9 Hz, 1H), 7.14 (s br, 1H), 7.37 – 7.42 (m, 2H), 7.42 – 7.50 (m, 4H), 7.92 (s, 1H), 8.01 – 8.10 (m, 4H), 8.13 (d, J = 9.3 Hz, 1H), 8.20 (s br, 1H), 8.22 (s, 1H), 8.81 (s, 1H), 9.81 (s, 1H), 12.72 (s, 1H), 12.80 (s, 1H).

$^{13}$**C NMR** (150 MHz, DMSO-$D_6$) δ ppm: 173.09, 170.63, 165.69, 164.98, 159.35 (d, J = 240.6 Hz), 156.76, 154.65, 154.36, 152.79, 149.34, 142.85, 140.91, 139.86 (d, J = 11.0 Hz), 138.58, 137.12, 134.30, 131.83, 129.85, 128.56, 128.38, 125.57, 125.39, 118.15, 116.28, 113.67, 111.74 (d, J = 24 Hz), 111.55 (d, J = 22 Hz), 101.98, 97.96, 94.21, 69.90, 69.44, 64.27, 57.04, 56.47, 52.97, 51.27, 50.18, 48.92, 46.08, 45.84, 43.84, 39.38, 39.24, 37.92, 36.73, 34.51, 34.06, 33.24, 31.65, 31.28, 30.54, 24.85, 22.56, 20.60, 14.56.

**HR MS:** obs. m/z [M+H$^+$]: calculated for $C_{68}H_{87}N_{16}O_6ClF$, 1277.66616; found, 1277.66550; deviation: 0.5 ppm.

**NMR spectra**

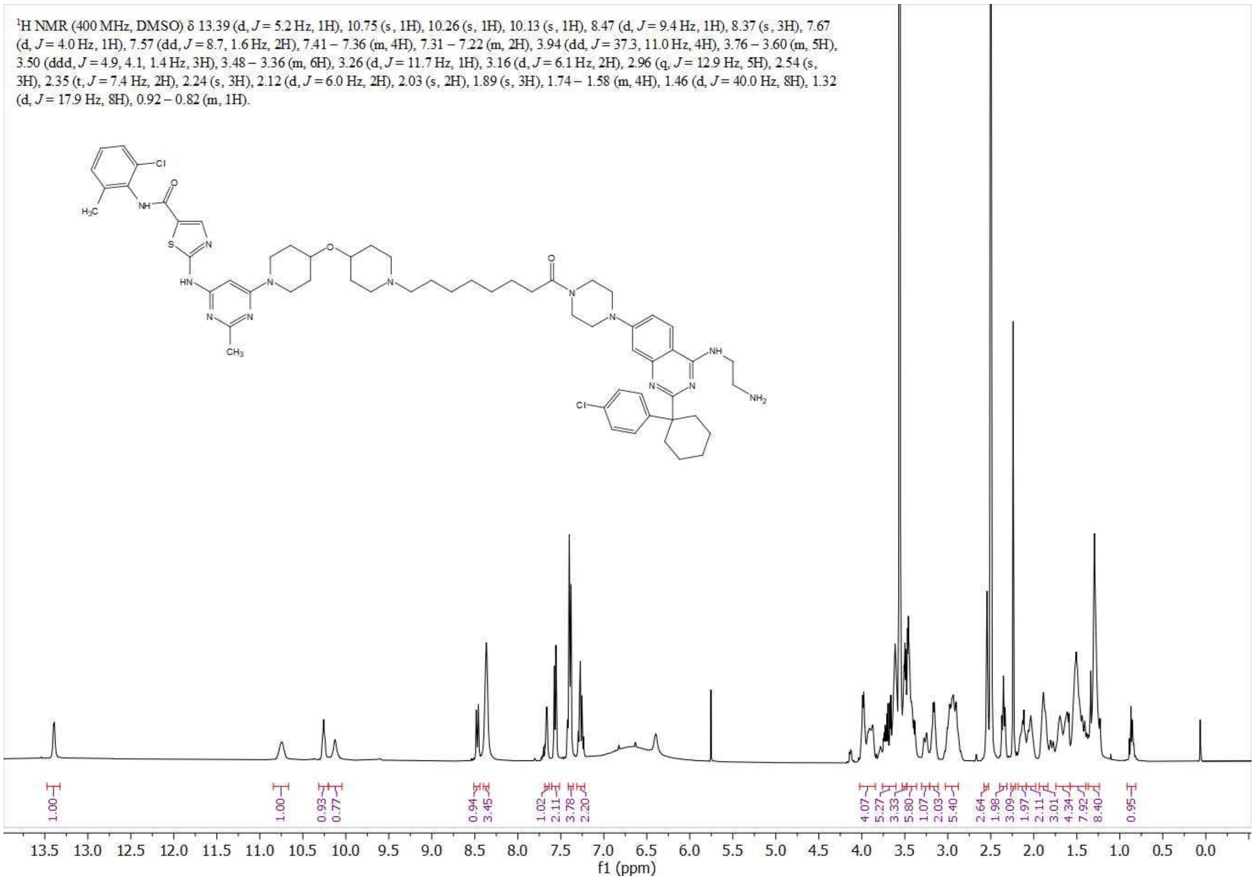


Supplementary spectrum 1: 1H-NMR spectrum of **DDa-1**

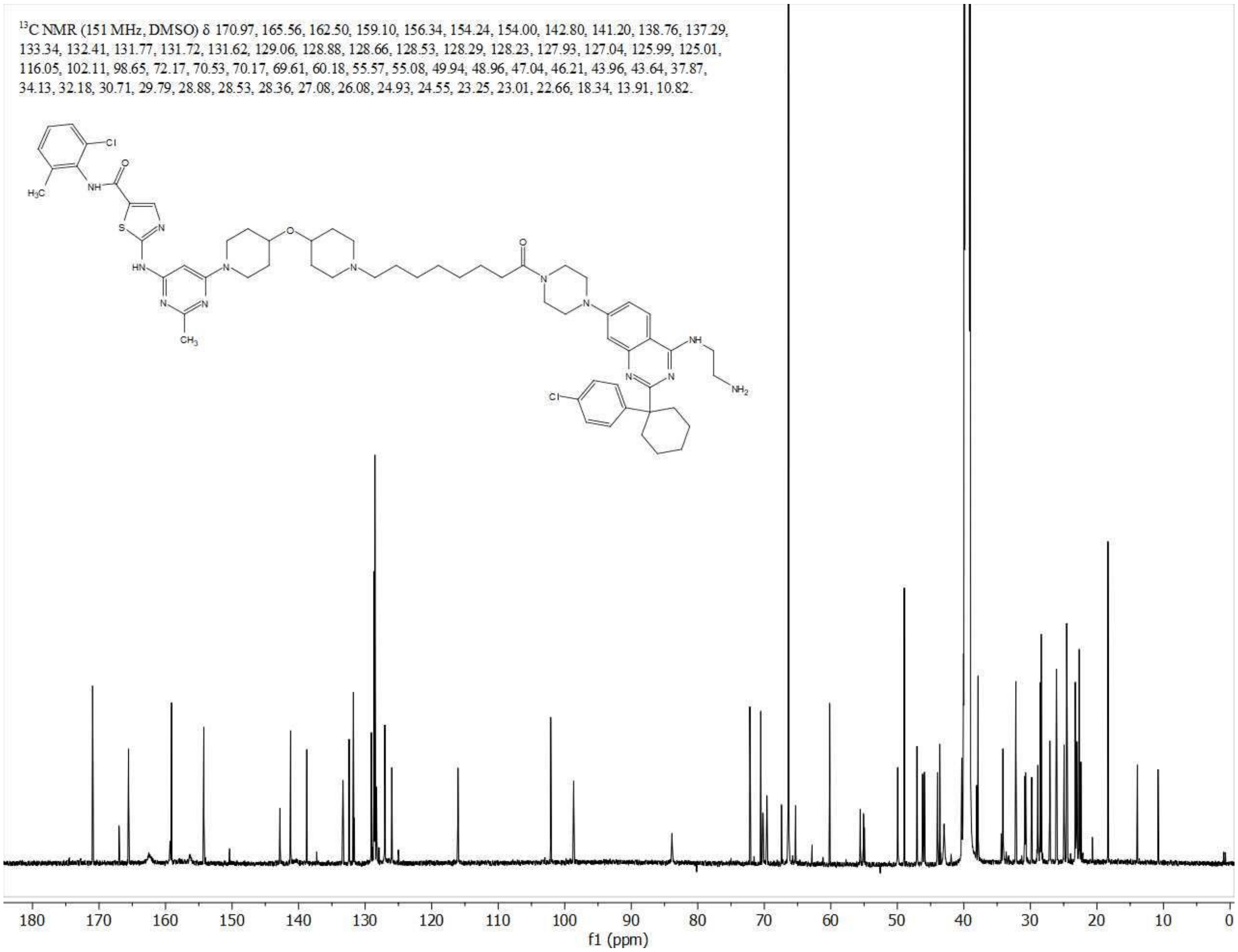


Supplementary spectrum 2: 13C-NMR spectrum of **DDa-1**

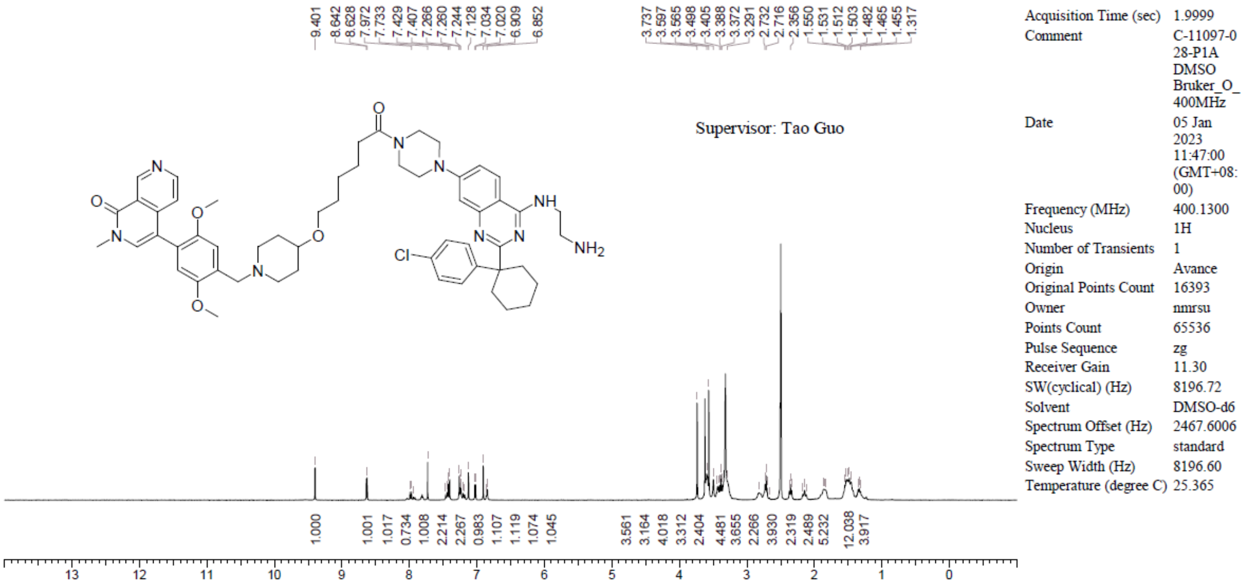


Supplementary spectrum 3: 1H-NMR spectrum of **DBr-1**

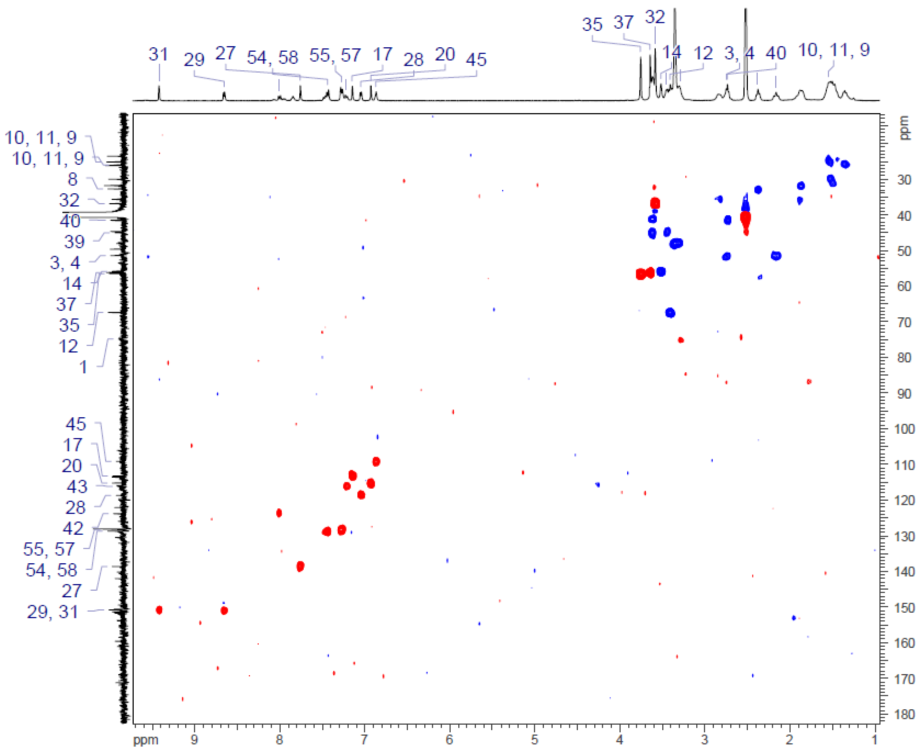


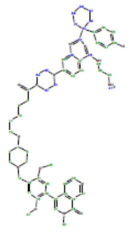

| | |
|---|---|
| *Acquisition Time (sec)* | (0.1828, 0.0064) |
| *Comment* | C-11097-028-P1B DMSO |
| *Date* | 06 Jan 2023 02:55:18 |
| *Frequency (MHz)* | (399.9340, 100.5635) |
| *Nucleus* | (1H, 13C) |
| *Number of Transients* | 4 |
| *Original Points Count* | (1024, 128) |
| *Points Count* | (2048, 512) |
| *Pulse Sequence* | hsqc_edetgp |
| *Solvent* | DMSO-d6 |
| *Spectrum Type* | HSQC-DEPT |
| *Sweep Width (Hz)* | (5598.27, 20074.71) |
| *Temperature (degree C)* | 21.350 |
| *Title* | C-11097-028-P1B DMSO |

Supplementary spectrum 4: HSQC-NMR spectrum of **DBr-1**

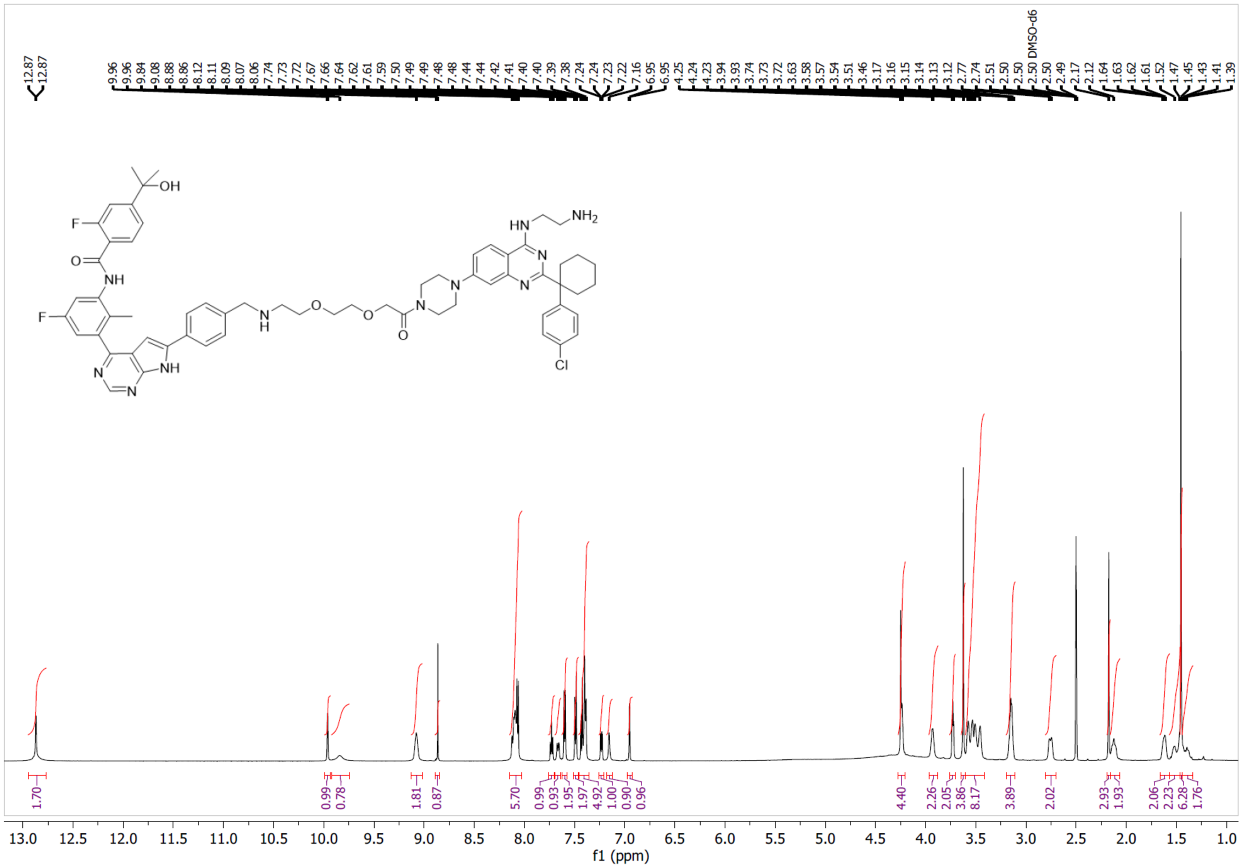

Supplementary spectrum 5: 1H-NMR spectrum of **DBt-3**

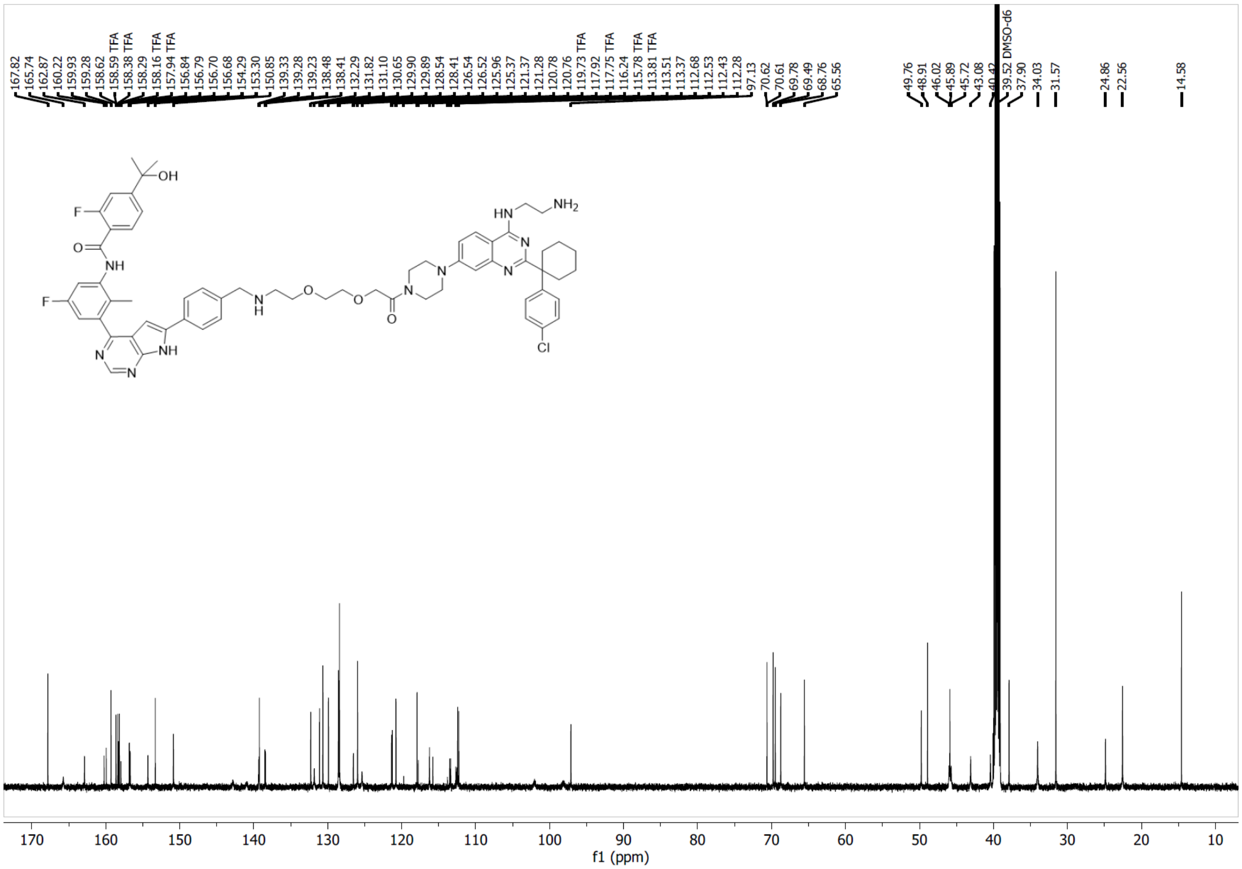

Supplementary spectrum 6: 13C-NMR spectrum of **DBt-3**

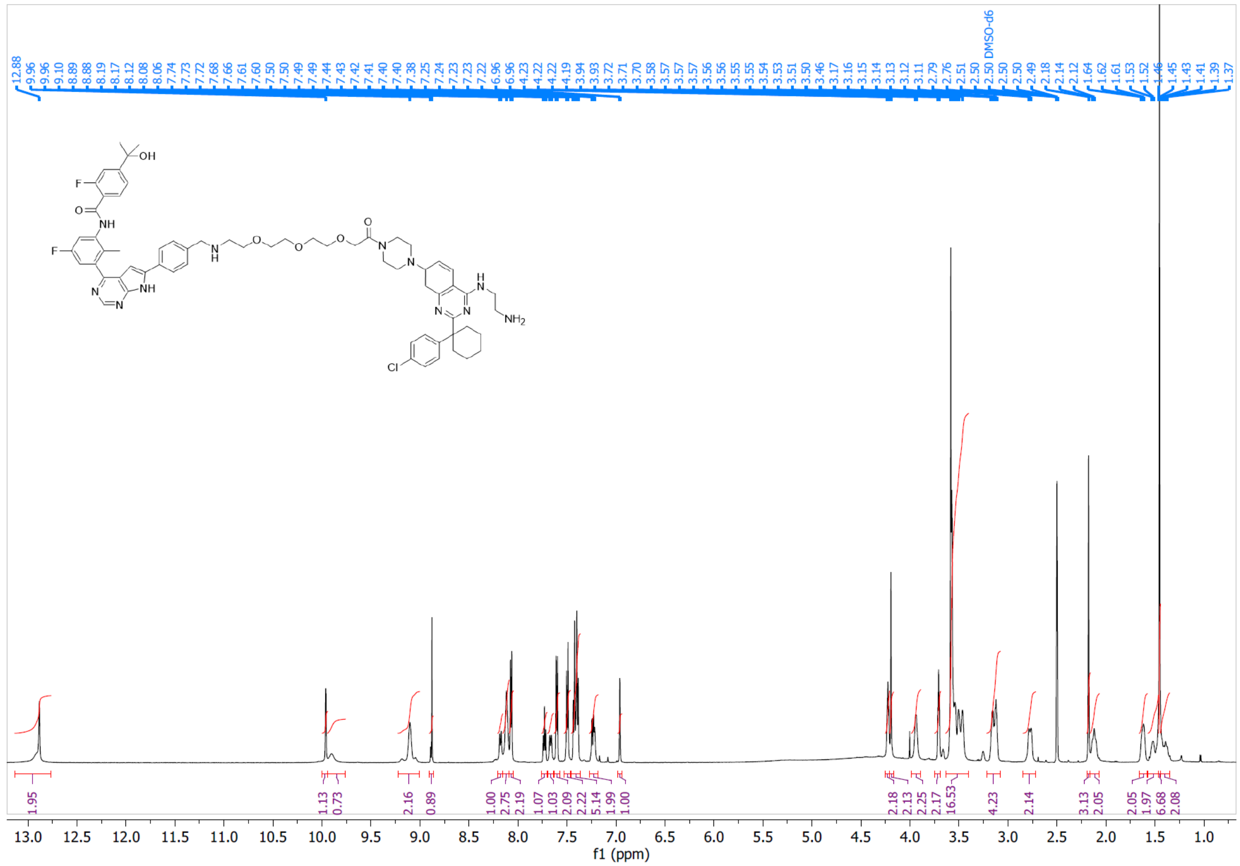

Supplementary spectrum 7: 1H-NMR spectrum of **DBt-4**

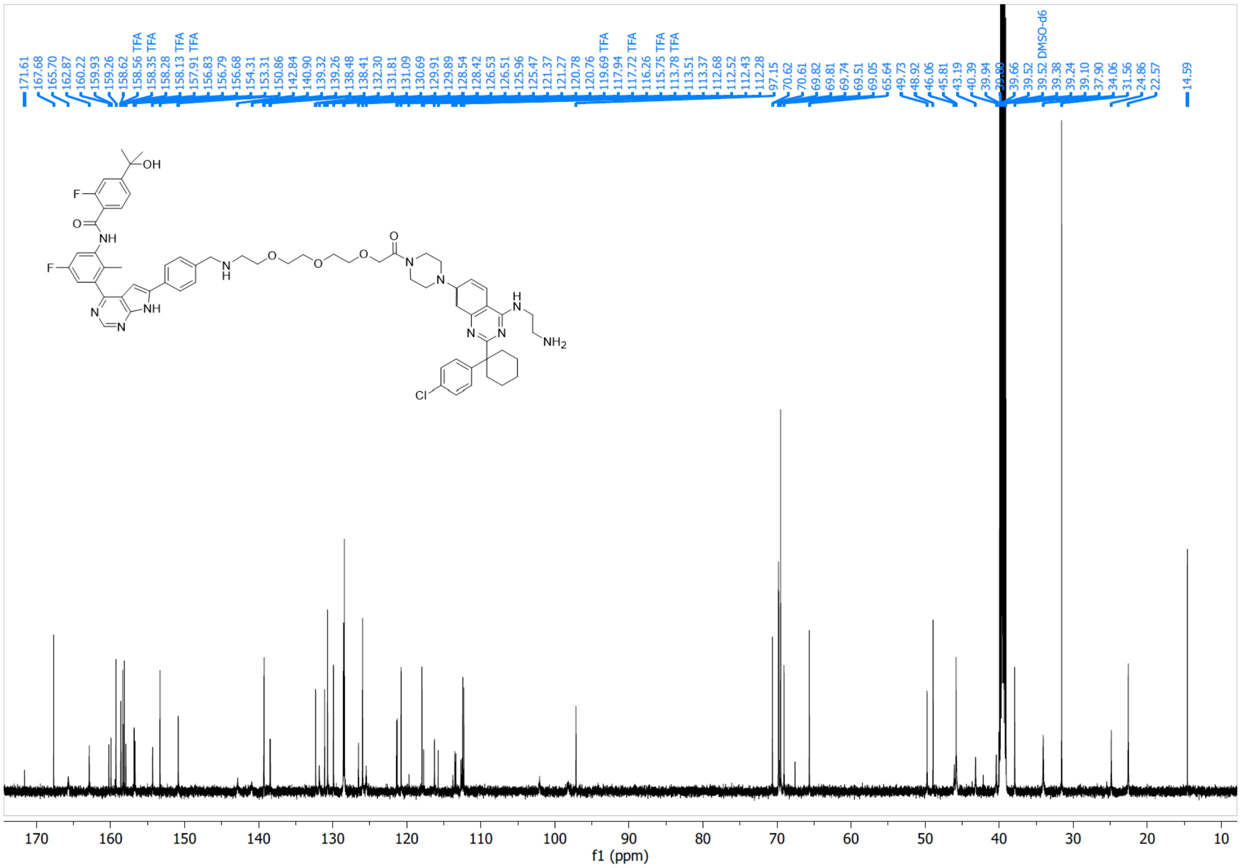


Supplementary spectrum 8: 13C-NMR spectrum of **DBt-4**

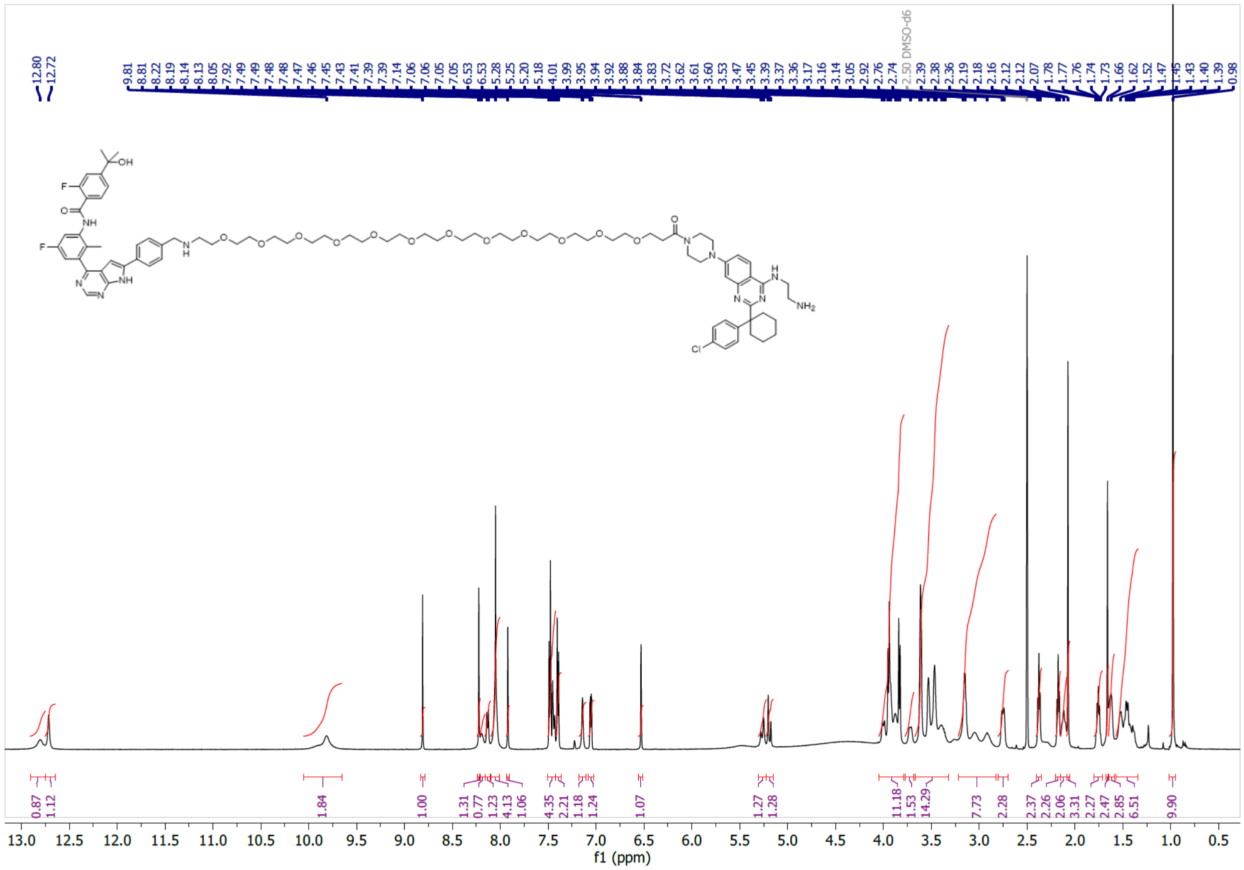

Supplementary spectrum 9: 1H-NMR spectrum of **DBt-5**

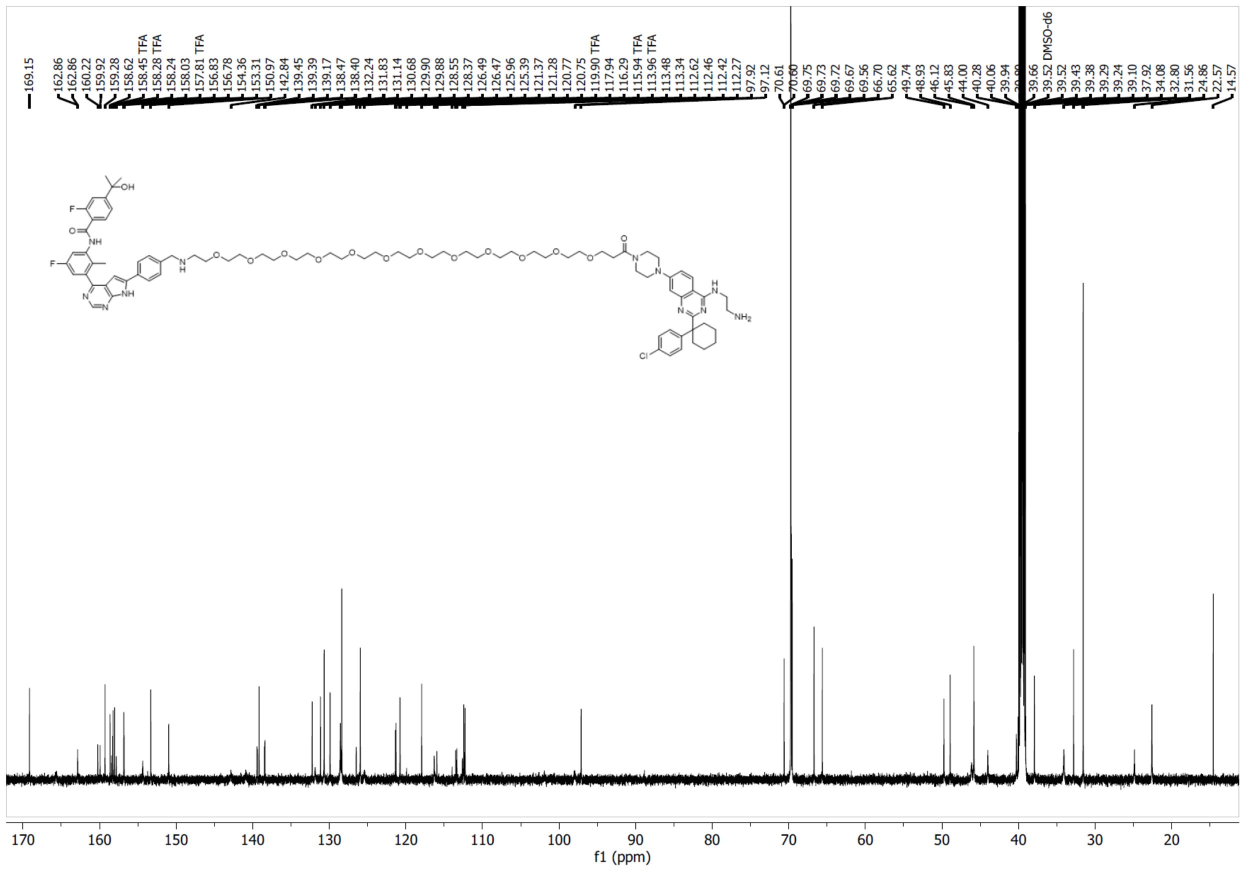

Supplementary spectrum 10: 13C-NMR spectrum of **DBt-5**

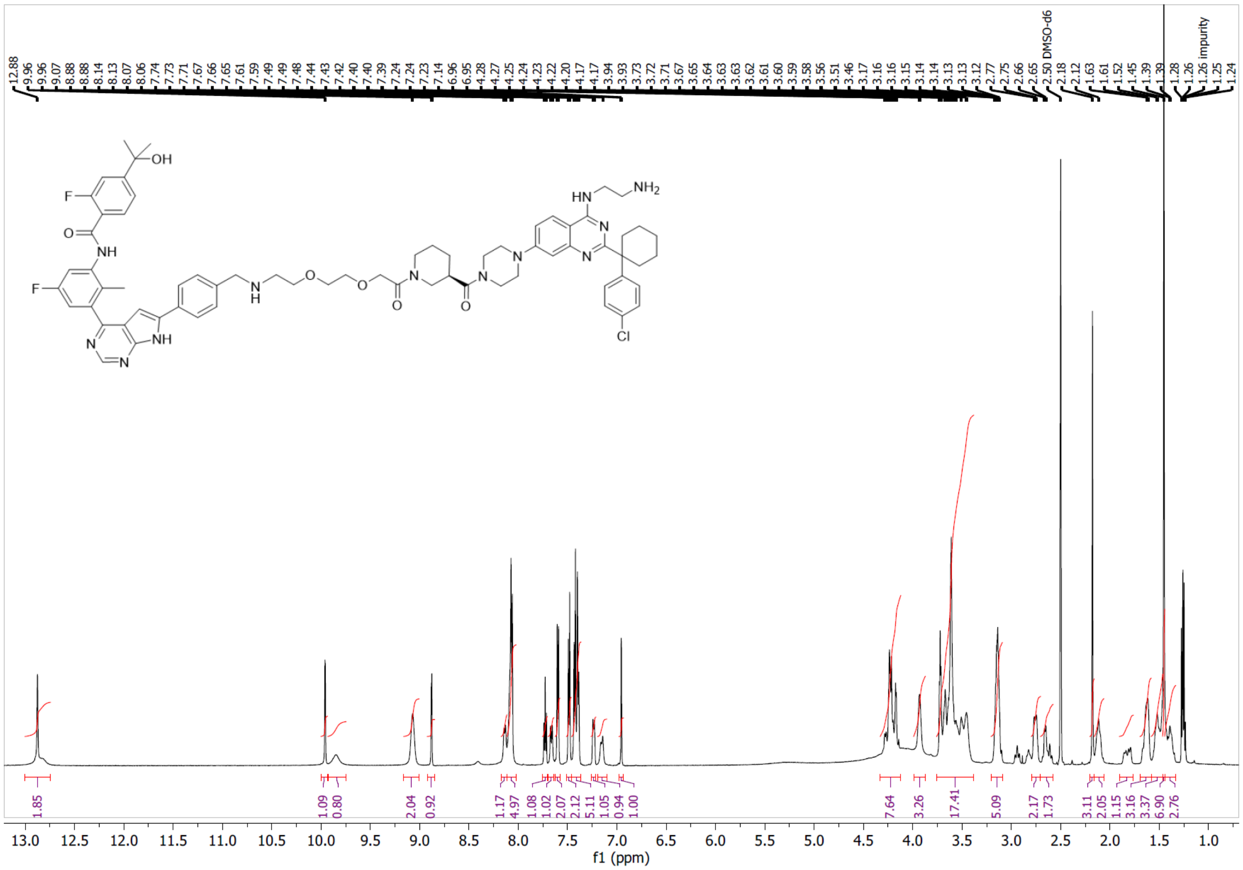

Supplementary spectrum 11: 1H-NMR spectrum of **DBt-6**

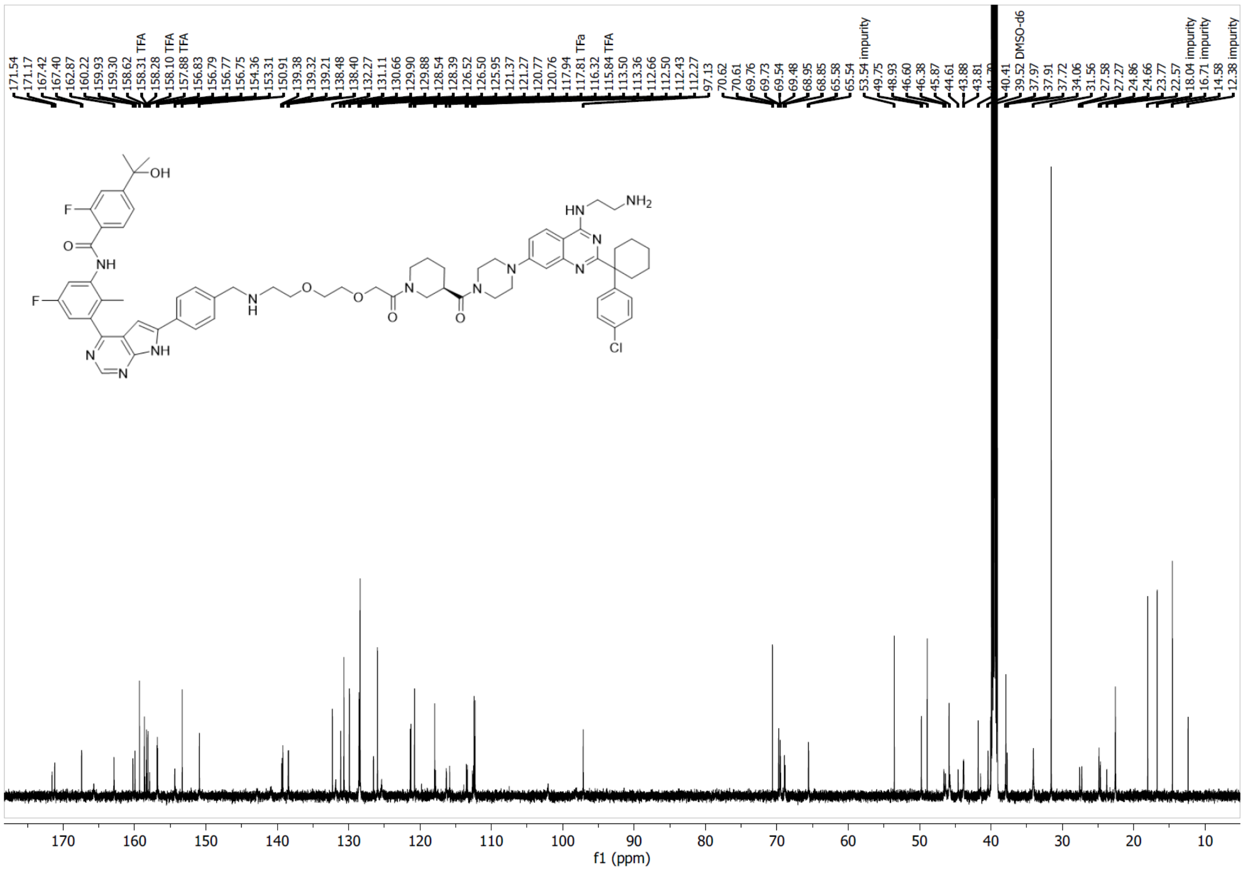

Supplementary spectrum 12: 13C-NMR spectrum of **DBt-6**

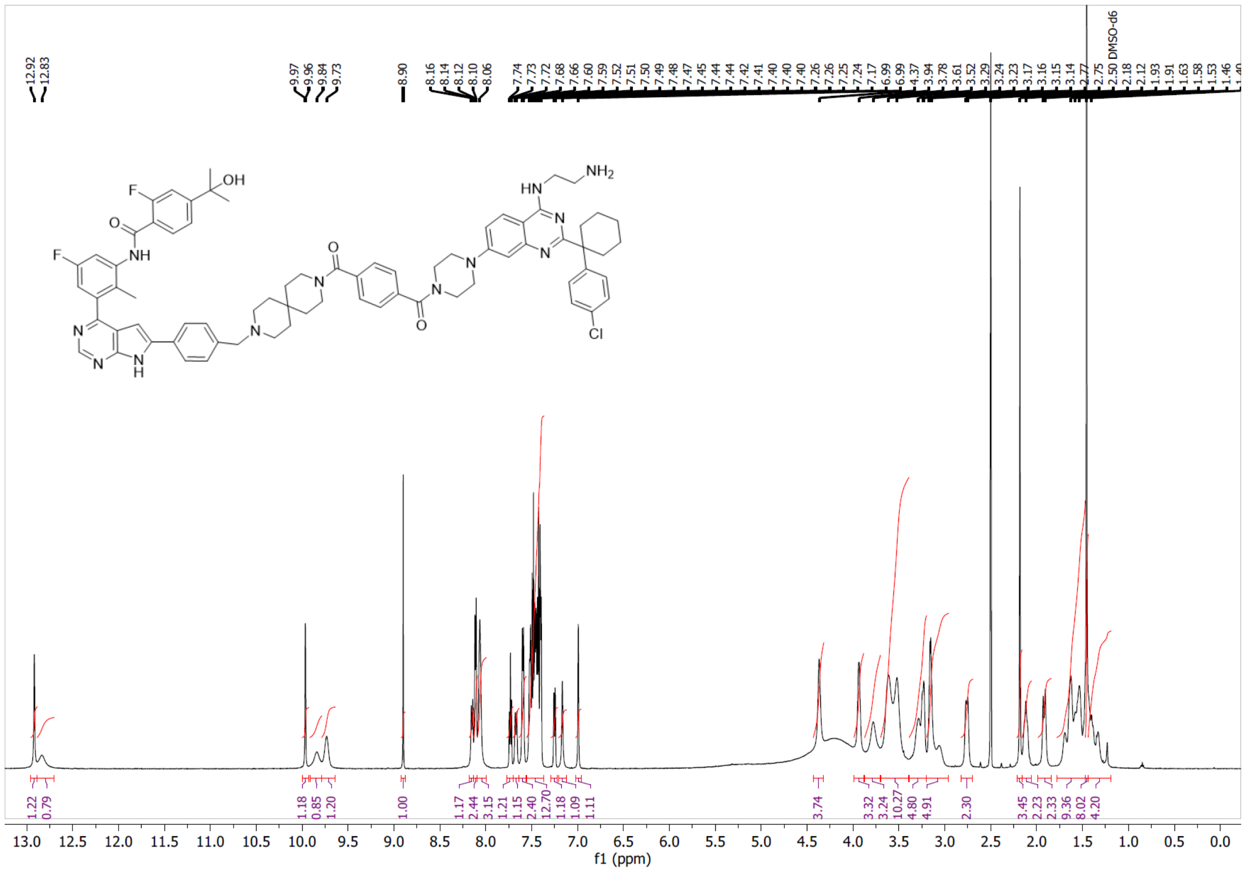

Supplementary spectrum 13: 1H-NMR spectrum of **DBt-8**

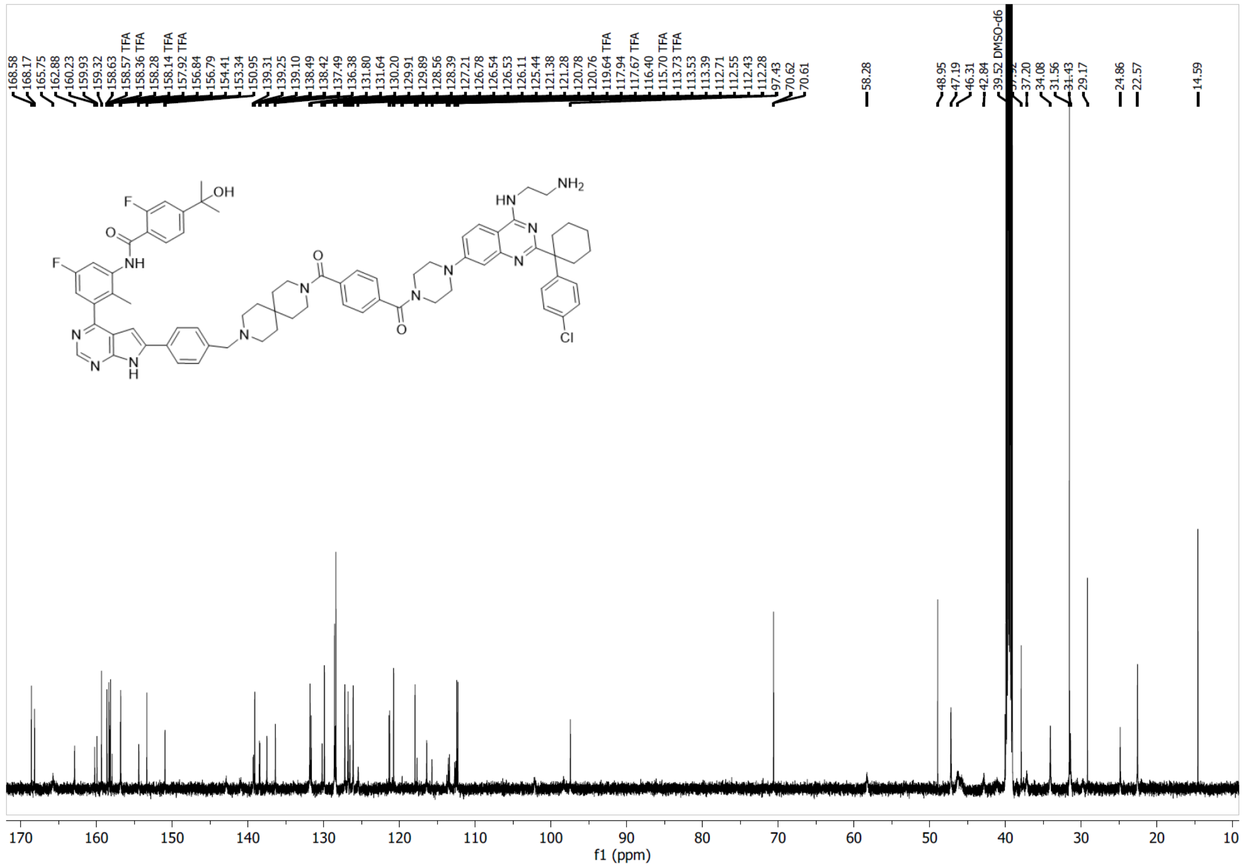

Supplementary spectrum 14: 13C-NMR spectrum of **DBt-8**

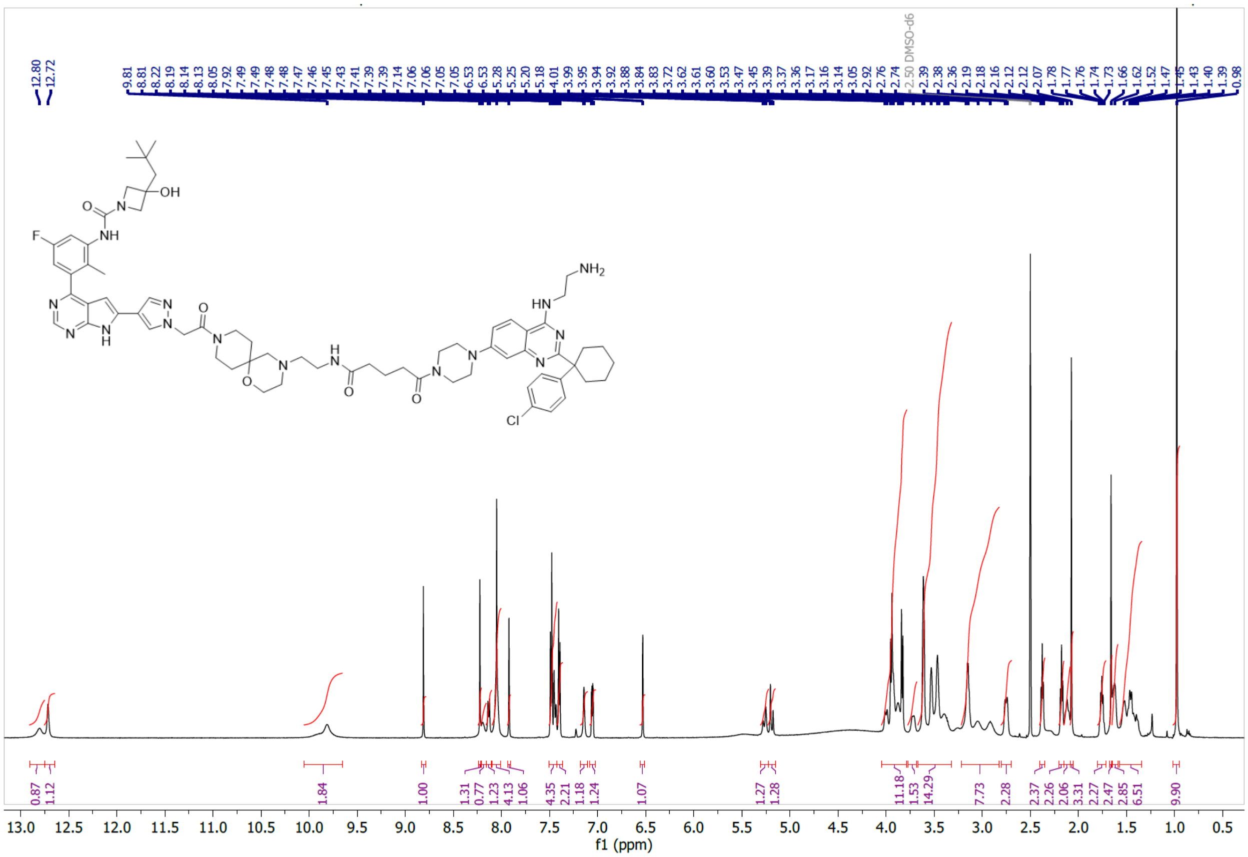

Supplementary spectrum 15: 1H-NMR spectrum of **DBt-10**

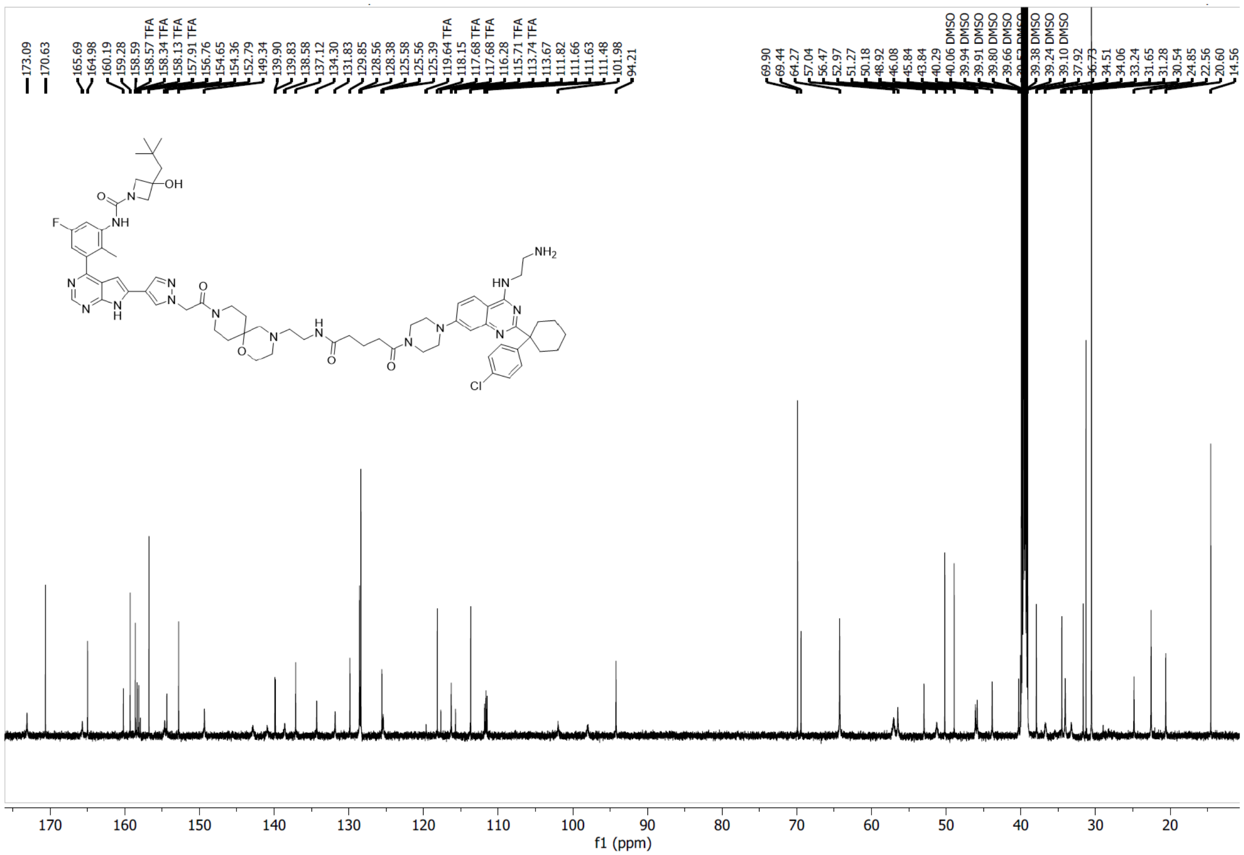


Supplementary spectrum 16: 13C-NMR spectrum of **DBt-10**

### High resolution MS spectra

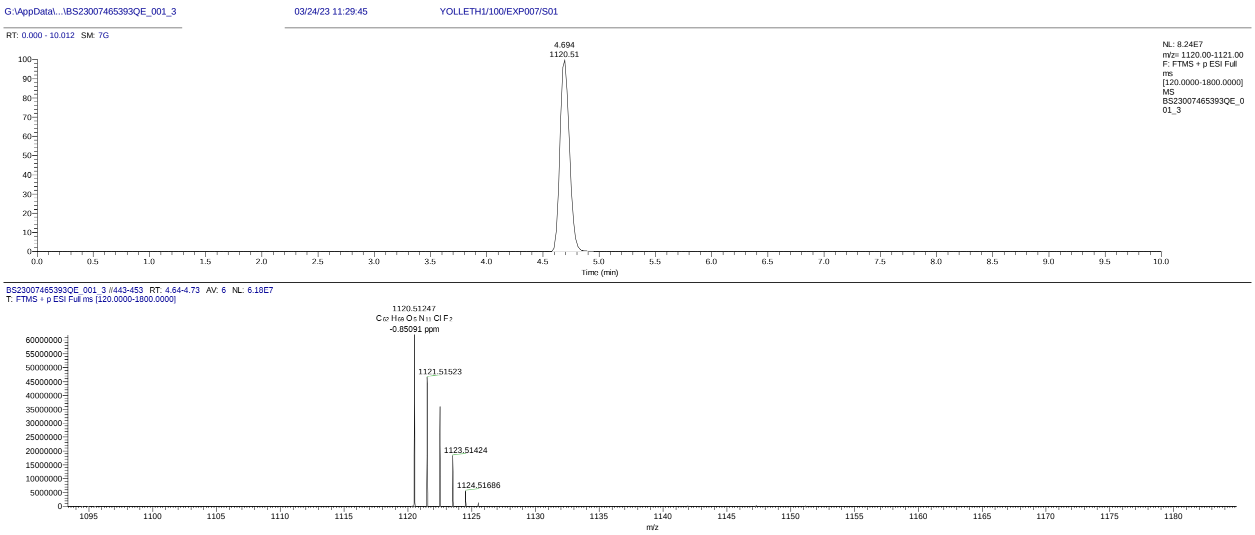


Supplementary spectrum 17: LC-ESI total ionization count spectrum and m/z spectrum of **DBt-3**

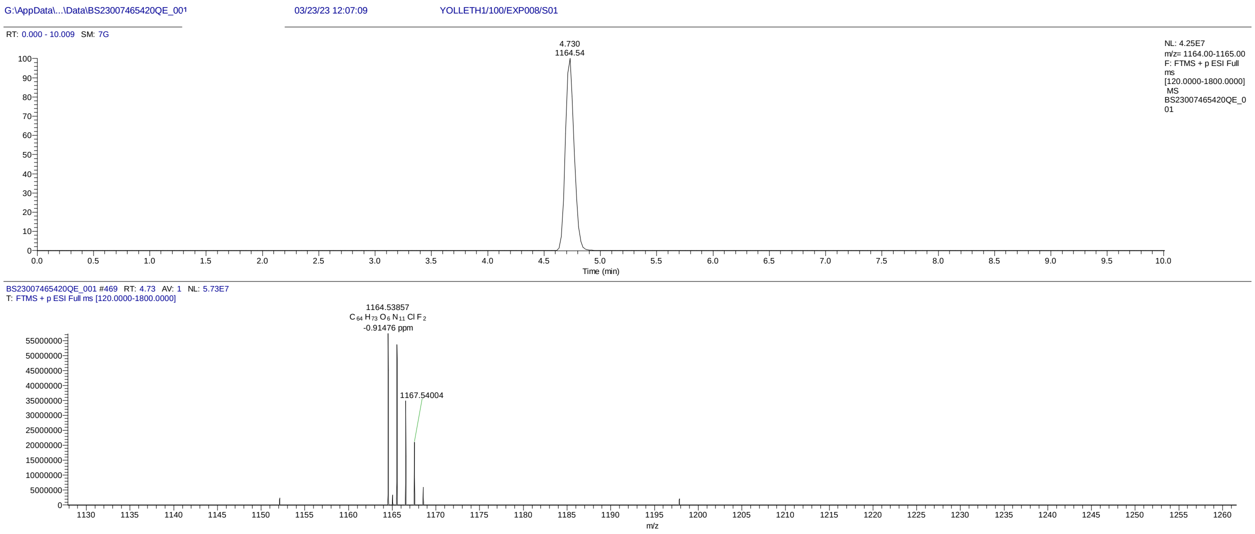


Supplementary spectrum 18: LC-ESI total ionization count spectrum and m/z spectrum of **DBt-4**

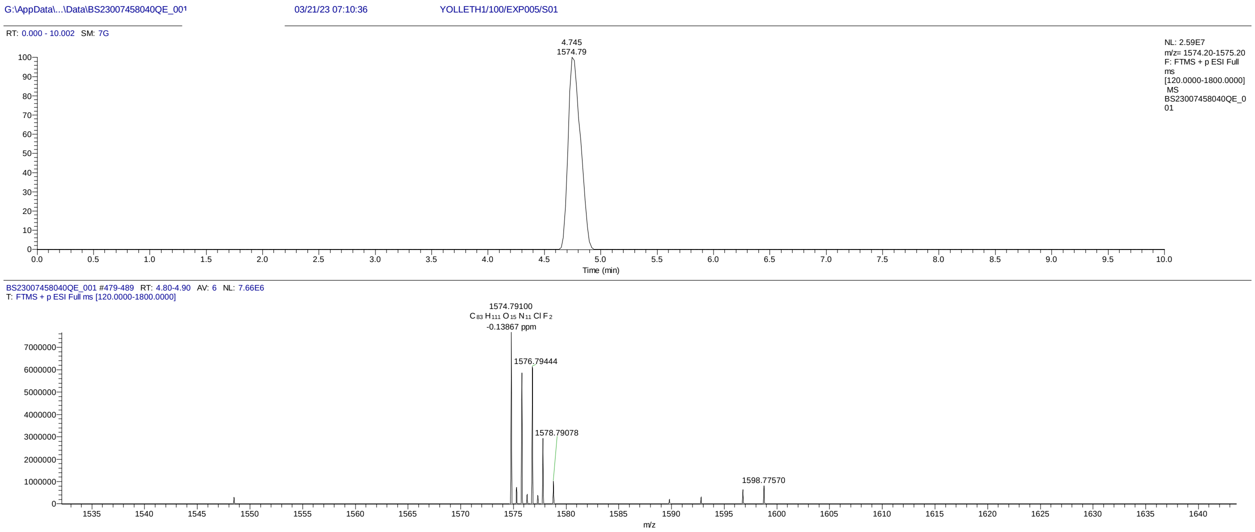


Supplementary spectrum 19: LC-ESI total ionization count spectrum and m/z spectrum of **DBt-5**

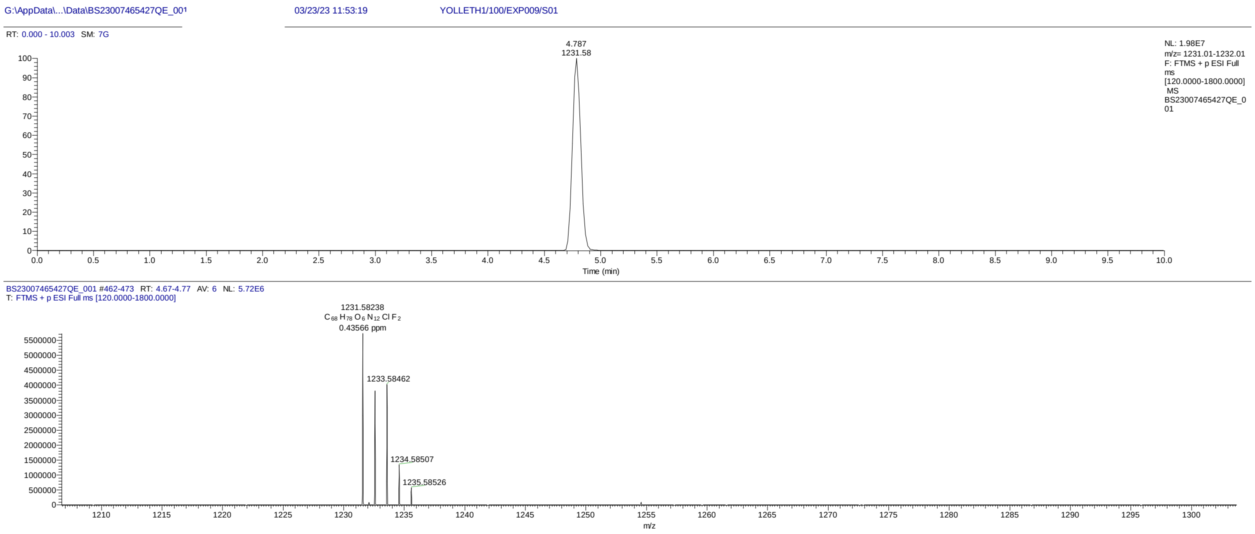


Supplementary spectrum 20: LC-ESI total ionization count spectrum and m/z spectrum of **DBt-6**

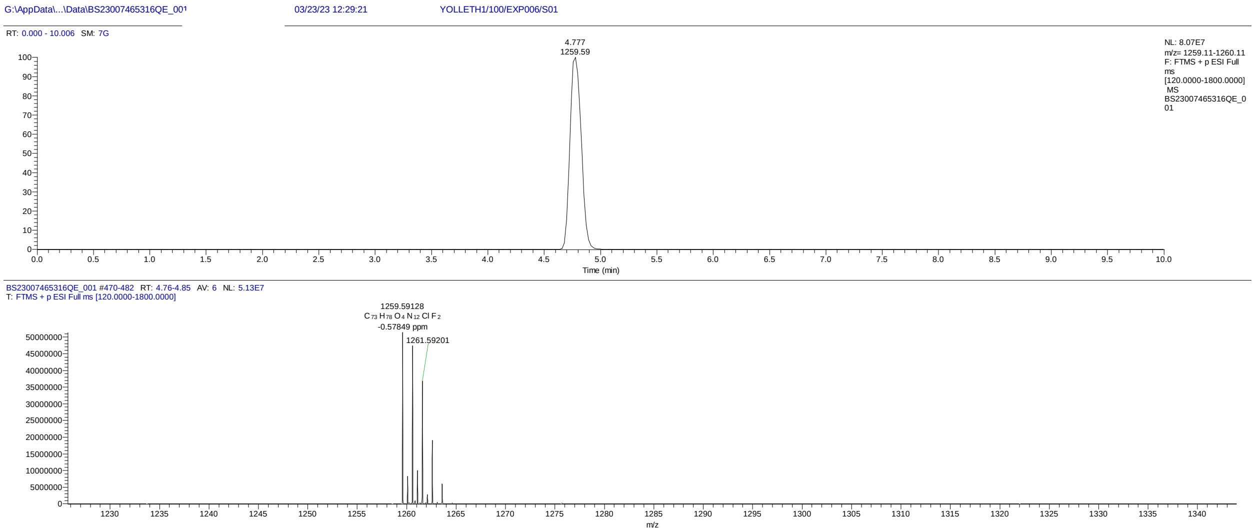


Supplementary spectrum 21: LC-ESI total ionization count spectrum and m/z spectrum of **DBt-8**

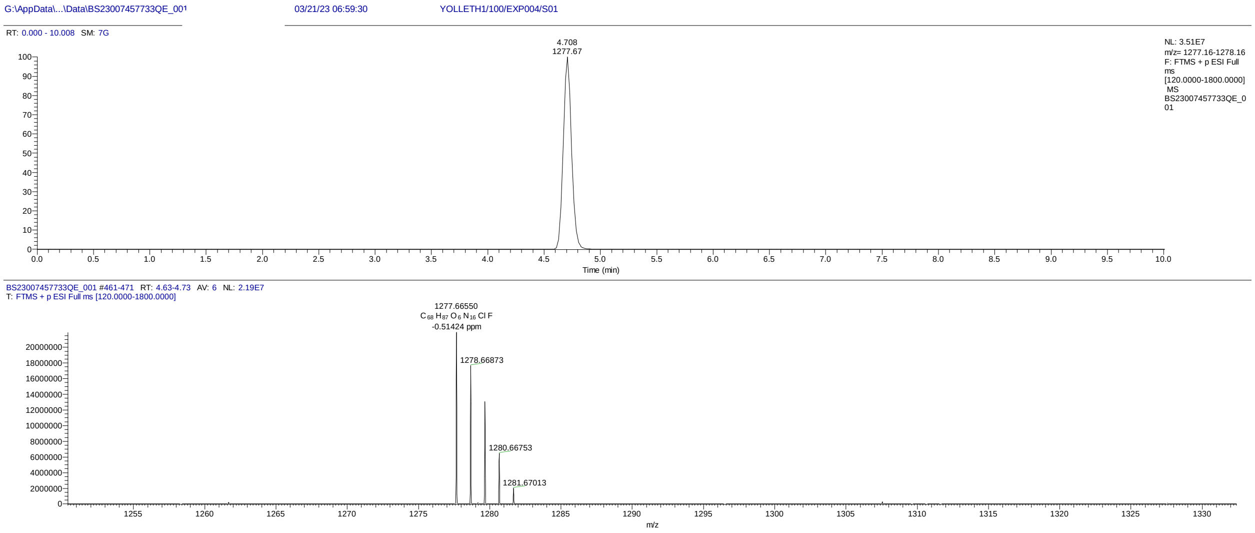


Supplementary spectrum 22: LC-ESI total ionization count spectrum and m/z spectrum of **DBt-10**
